# Integrated single cell analysis reveals co-evolution of malignant B cells and the tumor microenvironment in transformed follicular lymphoma

**DOI:** 10.1101/2022.11.17.516951

**Authors:** Clémentine Sarkozy, Shaocheng Wu, Katsuyoshi Takata, Tomohiro Aoki, Susana B Neriah, Katy Milne, Talia Goodyear, Celia Strong, Tashi Rastogi, Daniel Lai, Laurie H Sehn, Pedro Farinha, Brad H Nelson, Andrew Weng, David W Scott, Jeffrey W Craig, Christian Steidl, Andrew Roth

## Abstract

Follicular lymphoma (FL) is the most common indolent form of non-Hodgkin lymphoma. Histological transformation of FL to a more aggressive form of lymphoma occurs with a linear incidence of 2-3% per year and is associated with poor outcome. Divergent clonal evolution and an altered tumour microenvironment (TME) have both been implicated in the transformation process. However, the phenotypic consequences of this evolution and its implication in reshaping the TME remain unknown. To address this knowledge gap we performed single cell whole genome (scWGS) and single cell whole transcriptome sequencing (scWTS) of paired pre/post transformation samples of 11 FL patients. We further performed scWTS analysis of additional 11 FL samples from patients that had not undergone transformation within 7 years. Our comprehensive single cell analysis revealed the evolutionary dynamics of transformation at unprecedented resolution. Computational integration of scWGS and scWTS allowed us to identify gene programs upregulated and positively selected during evolution. Furthermore, our scWTS analysis revealed a shifting TME landscape, with an exhausted CD8 T cell signature emerging during transformation. Using multi-color immunofluorescence we transferred these findings to a novel TME based biomarker of transformation, subsequently validated in 2 independent cohorts of pretreatment FL samples. Taken together, our results provide a comprehensive view of the combined genomic and phenotypic evolution of malignant cells during transformation, and the shifting cross-talk between malignant cells and the TME.

**Graphical abstract:** 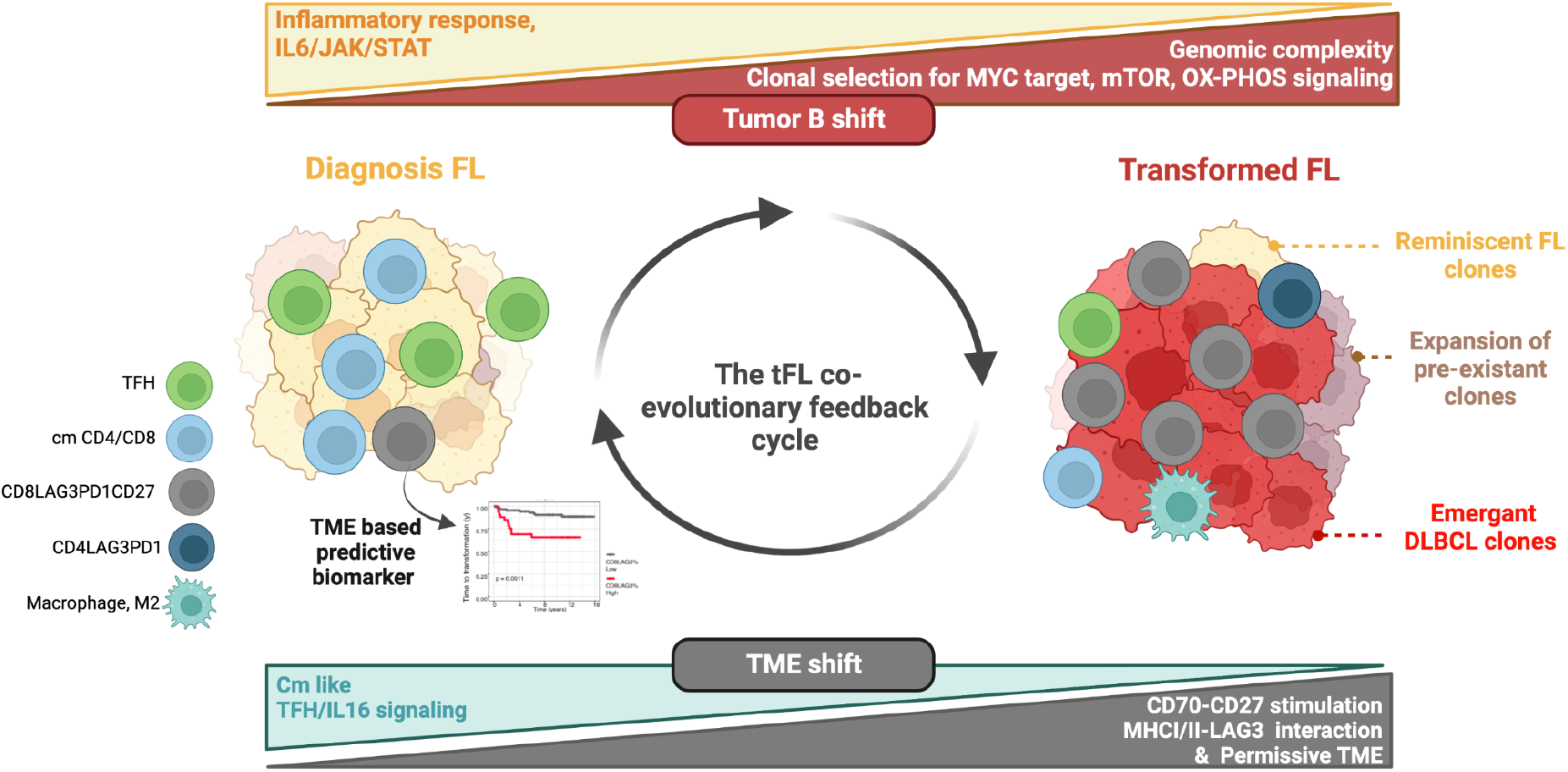

## Introduction

Follicular lymphoma (FL) accounts for approximately 20% of all non-Hodgkin lymphoma (NHL), making it the most common indolent form, and is associated with a median overall survival (OS) that exceeds 10 years (Bachy et al., 2019). The disease is characterized by a highly heterogeneous clinical evolution where some patients remain under observation and free of treatment for decades, while others transform into an aggressive and life-threatening form of B-cell NHL. Transformation manifests most frequently as diffuse large B-cell lymphoma (DLBCL) (Swerdlow et al., 2016), with a linear incidence of 2-3% per year and associated poor outcome (Al-Tourah et al., 2008; Federico et al., 2018; Sarkozy et al., 2016; Wagner-Johnston et al., 2015). The temporal aspect of the transformation process, including an indolent followed by an aggressive form of disease, makes FL an excellent model for studying cancer dynamics and evolution.

Initiating events in FL involve the t(14;18) translocation alongside epigenetic modifier mutations (Green, 2018; Sarkozy et al., 2019). The biological steps leading in transformation in DLBCL have been characterized as an evolutionary shift using phylogenetic analysis of sequential bulk biopsies (Carlotti et al., 2009; Kridel et al., 2016; Okosun et al., 2014; Pasqualucci et al., 2014). These bulk sequencing studies have shown that transformation is very likely the consequence of a branched evolution evolution from a common progenitor or ancestral clone (CPC), based on the existence of shared genetic lesions between primary and transformed FL (tFL) biopsies that precede the acquisition of time-point-specific FL and tFL secondary genomic abnormalities. An increased complexity and mutational burden was also shown in transformed biopsies, but without a unifying molecular driver mechanism (Bouska et al., 2014, 2017). Whether the technical detection thresholds inherent to bulk sequencing hampered the detection of CPCs as well as transformed clones at the time of initial FL diagnosis remains an open question that cannot be addressed without single-cell sequencing using paired pre and post-transformed biopsies. In addition, the relationship between clonal evolution (genetic) and disease evolution (phenotypic) has yet to be demonstrated. In particular, whether the transformation phenotype is governed primarily by clonal (genetic) evolution or other (epigenetic) events is an area of intense interest and investigation. To date, bulk gene expression studies have demonstrated high inter- and intra-cohort heterogeneity, and have not been able to show a clear “road to transformation” (Brodtkorb et al., 2014; Davies et al., 2007; Elenitoba-Johnson et al., 2003; Gentles et al., 2009; Lossos et al., 2002; Martinez-Climent et al., 2003). Several clinico-biological risk models (Huet et al., 2018; Jurinovic et al., 2016) attempting to predict early transformation or relapse in FL have been developed, but show limited reproducibility (Bolen et al., 2021).

Studies have described the potential roles of several TME components during transformation (Carreras et al., 2006, 2009; Dave et al., 2004; Farinha et al., 2010; Glas et al., 2007; Tobin et al., 2019; Tzankov et al., 2008); however, the results are somewhat discordant, likely due to non-exhaustive bulk characterization. The full range and dynamics of TME/tumor B cell interactions during transformation have not been fully explored.

By applying an unprecedented series of high-dimensional single cell RNA and DNA profiling techniques, we here characterize the clonal and phenotypic co-evolution of tumor B cells and their TME in transformation providing avenues for the development of innovative immunotherapies. Using a control cohort of long-term non-transformed FL cases (non-tFL), we additionally searched for potential predictive markers of transformation, both within tumor B cells and the TME.

## Results

### Single cell resolution profiling of the FL microenvironment

To characterize the microenvironmental determinants of transformation we assembled a cohort of 11 FL patients with pre/post transformation (tFL-FL/tFL-DLBCL) paired biopsies, alongside an additional 11 FL patients who had no disease progression within six years (non-TFL), and two reactive lymph node (RLN) samples which served as healthy controls. Combined single cell whole transcriptome and BCR sequencing (scWTS+BCR) was performed for all 35 samples; 129,142 live cells were recovered, with a median of 1,290 genes detected per cell. To identify the evolutionary dynamics of transformation in the 11 transformation pairs, we performed single cell whole genome sequencing (scWGS) using the recently developed direct library plus (DLP+) platform (Laks et al., 2019). We obtained 13,458 high quality single cell genomes after quality control. Finally, to explore the spatial organization of these samples we constructed a tissue microarray (TMA) for the entire tumor cohort (including 2 cores for each sample) and performed multi-colour immunohistochemistry (MC-IHC). An overview of the study workflow is presented in Figure 1. Clinical and biological characteristics of the 11 tFL-pairs and 11 non-tFL patients are presented in Table 1 and Supplementary Figure 1, Supplementary Figure 2A-B.

**Figure 1.**
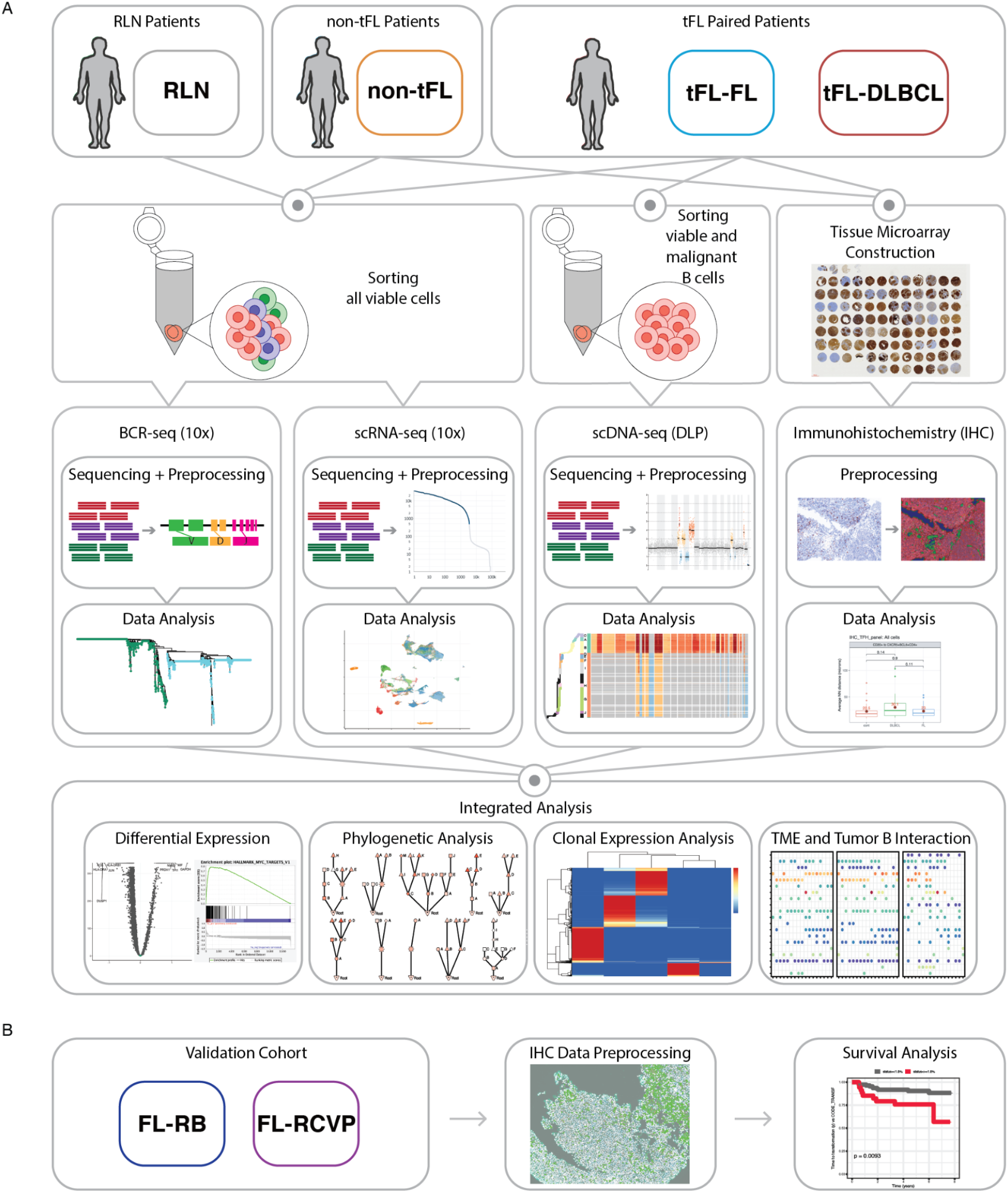
Overview of the workflow. **Panel A:** To characterize the transcriptional and genomic co-evolutionary process occurring in TME and tumor B cells during FL transformation, we performed scRNAseq (5’ and BCR sequencing) and DNAseq on single-cell suspensions collected from paired lymph nodes of 11 patients with documented transformed FL in DLBCL (within pairs: tFL-FL samples are color-coded in blue and tFL-DLBCL samples in red). As a control, we performed scRNAseq on single-cell suspensions of lymph nodes FL samples collected at diagnosis from 11 patients with a non-progressed FL (so called non-tFL, color-coded in orange). In parallel, 2 RLN (ie non-malignant, color coded in grey) were also processed with scRNA. To avoid batch effect, each pair (including tFL-FL and tFL-DLBCL timepoints) was processed with a non-tFL sample and RLN were not included in the same batch. A tissue microarray (TMA) was built from FFPE samples for all the patients and timepoints. Multicolor immunohistochemistry was used to assess distance analysis. **Panel B**: For outcome prediction, we used 2 independent cohorts of FL patients with available samples and TMA: one treated with R-Bendamustine (R-B) and previously published (Freeman et al., 2019) and treated with R-CVP (rituximab, cyclophosphamide, vincristine and prednisone), previously published (Kridel et al., 2016).

### Single cell analysis reveals the cellular composition of FL and patient specific malignant phenotypes

To perform a systematic comparative analysis of the 11 tFL-pairs, 11 non-tFL and 2 RLN, we merged the scWTS expression data from all cells (35 samples) and performed batch correction and normalization. Removal of batch effects (caused by single-cell isolation and library preparation in different experimental runs) resulted in improved mixing of cells across samples, as demonstrated by a significant increase in cell entropy (Wilcoxon–Mann–Whitney P < 0.001; Supplementary Figure 3).

Unsupervised clustering followed by visualization in UMAP space identified 39 expression-based cell clusters that were annotated as non-B TME cell clusters (T cell, macrophage, NK cell and dendritic cell clusters), or B cell clusters based on expression of key markers (Supplementary Figure 4). The B cells were then re-clustered independently (N=25 clusters) and further classified as “tumor B” or “normal B” (cluster 1-3-9 and 22) based on the ratio of kappa/lambda gene expression and BCR sequencing analysis (Supplementary Figure 5). A unique clonal V-D-J recombination was identified in all tumor samples, except in tFL-pair-9 where no productive clonal V-D-J could be found, likely due to the abundance of somatic hypermutation (SHM). No clonal V-D-J recombination events were detected among B cells from the 2 RLN samples (Supplementary Figure 5C; Supplementary Table 2).

The tumor B cells were then re-clustered, leading to 23 independent clusters that segregated strongly by patient but not by timepoint. Tumor sample distance analysis also emphasized the similarity within pairs (Supplementary Figure 6D). Of note, the patient related clustering effect was even stronger when reclustering only the malignant B cells from pairs (Supplementary Figure 7).

### Single cell genome sequencing precisely defines the evolutionary history of transformation

We next sought to determine the evolutionary features of malignant cells associated with transformation. Using scWGS we inferred single cell copy number profiles, which were then used to construct phylogenetic trees showing the genomic clonal evolution from tFL-FL to tFL-DLBCL (Figure 2). In general, tFL-DLBCL samples presented a greater genomic complexity with more subclones and copy number variants (CNVs) than tFL-FL samples (Supplementary Table 3).

**Figure 2.**
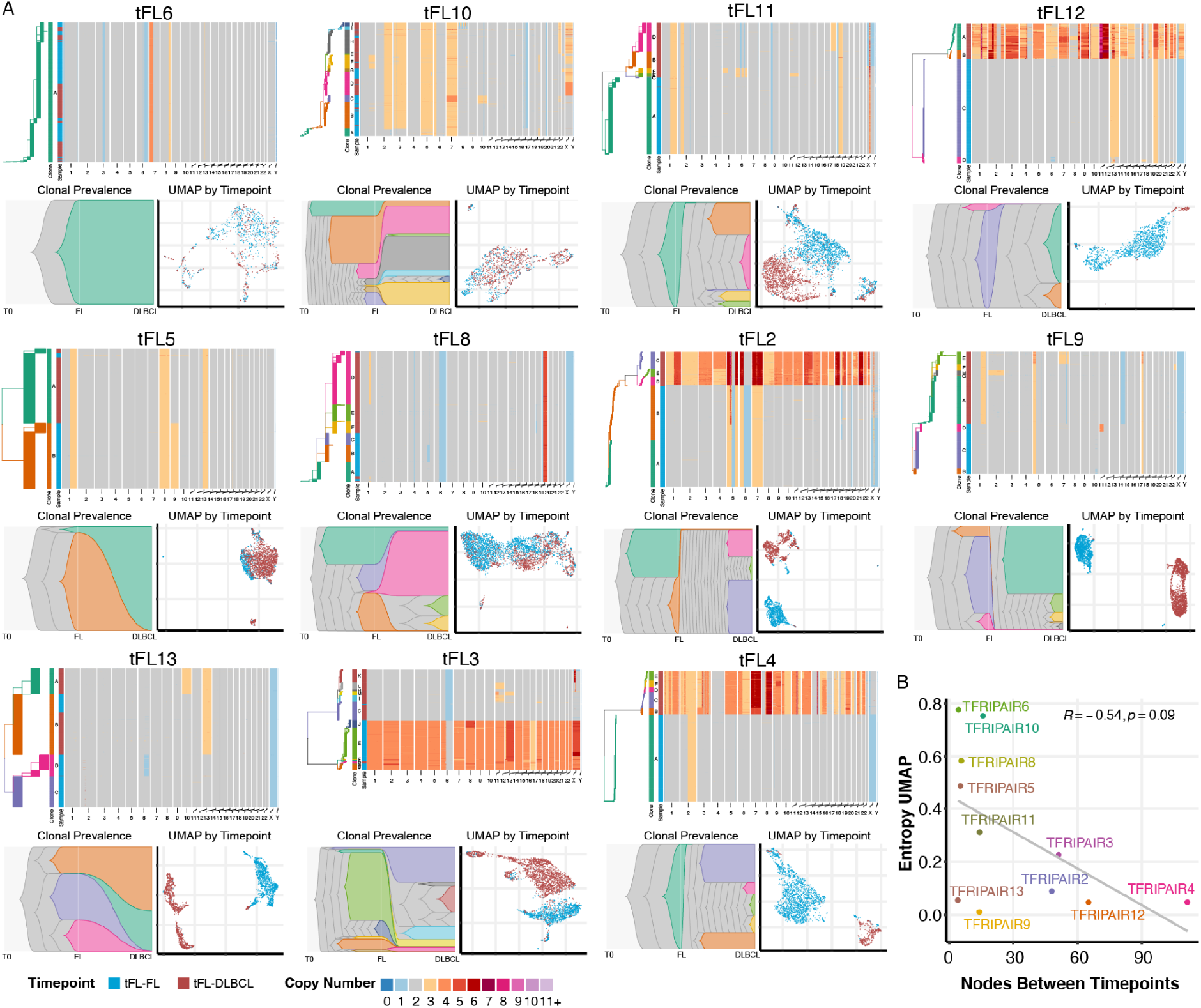
Phenotypic and genomic analysis of tumor B cells in paired transformed follicular lymphoma samples. **Panel A:** Each sub-panel includes from left to right pair level: phylogenetic tree from scDNA-seq data colored by clone (tree scales are different among pairs: wider tips represent smaller scales, and narrower tips larger scales), a vertical bar representing the clonal composition of the pair, aligned another line showing the timepoint composition (ie tFL-FL in blud and tFL-DLBCL in red) and a copy number heatmap that shows the total CN profiles (not allele specific), with CN value color coded from blue (loss) to brown (amplification). The lower part includes a timescape figure representing the clonal evolution from tFL-FL to tFL-DLBCL timepoints (lower left sub-panel), and a UMAP visualization of the 10X scRNA-seq data by pair, blue dots representing tFL-FL cells and red dots tFL-DLBCL timepoint cells (lower right sub-panel). tFL-pairs are sorted based on the level of similarity between tFL-FL and DLBCL timepoint, i.e presence or absence of “mixed” DLP clones and/or fully novel clones in DLBCL biopsies: from a high level of similarity, on the left, to a low level, on the right. tFL-FL timepoint cells could be observed in tFL-DLBCL timepoint clones in half of the pairs (tFL-pair-6, clone D tFL-pair-10, clone D tFL-pair-8, clone A and B tFL-pair-13, clone A tFL-pair-5, clone D&C&M tFL-pair-3, called mixed DLBCL clones). Conversely, in 70% of the pairs, tFL-DLBCL timepoint cells were present in tFL-FL timepoint clones (called mixed-FL clones: clone A tFL-pair-8, clone B tFL-pair-2, clone B tFL-pair-3, clone B tFL-pair-10, clone A tFL-pair-11, clone C&D tFL-pair-13, clone tFL-pair-5). **Panel B:** Scatter plot representing the UMAP entropy (calculated based on the proportion of tFL-FL cells and tFL-DLBCL in the 20 nearest neighbors for each cell) versus number of nodes within DLP trees between the 2 timepoints, for each tFL-pair, to assess the correlation between the phenotypic and genomic magnitude of evolution during transformation.

The evolution from tFL-FL to tFL-DLBCL was not uniform across all pairs, but rather showed important variability in the nodal distance between cells from the two timepoints (number of nodes), as well as the fraction of mixed clonal composition within each tFL-pair (i.e fraction of tFL-FL versus tFL-DLBCL cells in each DLP clone) (Figure 2). Indeed, at one extreme, some tFL-pairs (11-2-4-9-12) contained tFL-DLBCL clones without any tFL-FL cells, supporting a completely divergent evolution from FL to DLBCL. On the other extreme, some tFL-pairs (6-5-13) yielded only tFL-DLBCL clones that were already present at the genetic level at time of FL biopsy (i.e. mixed clones), with a subsequent expansion. tFL-pair-6 is the most extreme example with only one clone detected in the FL and DLBCL samples, in accordance with the pathological data showing little distinction between the FL and transformed biopsies (DLBCL cells comprised just 5% of the total section in the transformed biopsy). Finally, some tFL-pairs (10-8-3) presented with a mixed mode of evolution at the time of DLBCL biopsy, with both an expansion of minor clones present at time of FL biopsies and the emergence of new clones. All in all, some tFL-FL timepoint cells could be observed in tFL-DLBCL timepoint clones in half of the pairs. Conversely, in 70% of the pairs, tFL-DLBCL timepoint cells were present in tFL-FL timepoint clones (called mixed-FL clones). The relative abundance of tFL-FL versus tFL-DLBCL timepoints cells and their position on the DLP trees suggest that the mixed DLBCL clones contain cells from the tFL-FL timepoint that might be early transformed cells or precursor cells of transformation. However, whether these clones present an FL or DLBCL-like phenotype is an important point to be assessed by looking at the phenotype at the clone level.

Finally, in some cases (tFL-pairs 8, 10, or 2), a clone emerging before any divergence and inferred to be a common progenitor clone (CPC) could be identified in tFL-FL samples. In tFL-pair-8, this clone was also present at the time of tFL-DLBCL.

### Phenotypic and genomic divergence reflect a co-evolutionary process associated with time between biopsies

We next quantified the phenotypic and genotypic evolution occurring during transformation. We performed re-clustering of the scWTS data at the pair level (Figure 2) and computed a per cell entropy using the 20 nearest neighbors in expression space. We took the average of this entropy score across cells in a pair to generate a dissimilarity score between FL and DLBCL timepoints within tFL-pairs (a low entropy meaning a high dissimilarity and vice-versa, Supplementary Figure 8A). Our results reveal a continuum of phenotypic evolution during transformation. But strikingly, in each pair, a minimum level of similarity between tumor B cells from tFL-FL and tFL-DLBCL timepoints was found, even in those presenting a very low entropy and distinct timepoint-related clusters (tFL-pairs 2-3-4-9-12-13): some tFL-FL cells were found within the tFL-DLBCL clusters (labeled as ‘DLBCL-like cells’) and some tFL-DLBCL cells were in the tFL-FL cluster.

A side-by-side analysis of genomic and phenotypic data highlighted, in the majority of the pairs, a co-evolutionary process where samples with low genomic changes present a high level of entropy (tFL-pairs 11-10-5-6 and 8), and vice-versa (tFL-pairs 4-3-12-2) (Figure 2 panel B, Supplementary Figure 8A). However, this correlation was not apparent in all tFL-pairs. In particular, tFL-pair 13 showed minimal clonal evolution, with the predominance of mixed clones composed on both tFL-FL and tFL-DLBCL cells, but a high degree of phenotypic evolution with 2 different tFL-FL and tFL-DLBCL clusters. As well, the strict co-evolution of malignant B-cell genotypes and phenotypes cannot account for the five tFL-pairs (11-2-4-12-9) with more extreme clonal divergence but the presence, at the phenotypic level in tFL-FL time point biopsies, of cells clustering with the tFL-DLBCL time point cells. These “DLBCL-like” phenotypic cells do not have a DLBCL-like genotype, but the full genomic divergence associated with the presence of tFL-FL-specific CNV (as loss in 5q33.3-34 and 9p22.1-21.3 in PAIR4 tFL-FL cells, absent in tFL-DLBCL cells) suggests that other factors such as epigenetic or TME related events account for this phenotype, rather than uncaptured genomic events such as SNV or structural variants. Finally, in tFL-pair-3, the CNV profile revealed complex changes in the tFL-FL sample with a high number of chromosomal gain (confirmed by FISH). Interestingly, FISH analysis identified a *MYC* translocation in the tFL-DLBCL sample, absent in the tFL-FL biopsy and very likely a co-driver of the phenotypic evolution.

Intriguingly, the phenotypic, and to a lesser extent the genomic, evolution through transformation, tended to be related to the time interval between the two biopsies (Supplementary Figure 8B-C-D) rather than the presence of chemotherapy or radiation exposure in the interval.

### Single cell analysis reveals upregulated MYC target gene signaling as a feature of transformation

Selecting only the tumor B cells, we performed a differential expression (DE) analysis between the 11 tFL-FL and 11 tFL-DLBCL samples, using timepoint, patient, and batch as covariate terms. Gene set enrichment analysis (GSEA) of the DE results was then performed using the Hallmark database (Subramanian et al., 2005). The “MYC targets V1”, “oxidative phosphorylation”, “mTORC1” and “glycolysis” pathways were significantly enriched in tFL-DLBCL cells versus “IL6-JAK-STAT”, “IL2-STAT5”, “inflammatory response”, “complement cascade” in tFL-FL cells (Supplementary Figure 9). These results suggest a shift in tumor B cell biology during transformation, with the hallmark of TME/tumor B cells interactions more present in tFL-FL as compared to tFL-DLBCL cells that carry a distinct metabolism reflecting proliferation. Interestingly, when we used all the cells as a pseudo-bulk DE, our analysis did not highlight any significant pathways enriched in tFL-FL, and the significance of those enriched in tFL-DLBCL was much lower (Supplementary Figure 10) suggesting that the precision afforded by single cell methods, to focus analysis on just the malignant cell compartment, is critical.

We calculated a “MYC_target_score” (MTS), as well as “mTORC1” and “OXPHOS” score per cells and confirmed that in each tFL-pair these scores were higher in tFL-DLBCL samples as compared to tFL-FL, except in tFL-pair-5 and tFL-pair-6 (Figure 3A; Supplementary Figure 11), in concordance with the pathological and genomic data. Importantly, this significant increase was not related to cell cycle state (Figure 3B, Supplementary Figure 11A). Using a 75% quantile threshold, we also showed an increase in the proportion of cells with a high MTS, mTORC1 and OXPHOS score from tFL-FL to tFL-DLBCL timepoints related to the interval of time between the 2 biopsies (Figure 3C, Supplementary Figure 11B and D). Of note, the evolution was not related to acquisition of rearrangements in the *MYC* gene locus in the majority of cases (assessed by FISH break-apart probes, see Supplementary Table 4). While we observed a high incidence of low-level *MYC* locus gains, these are likely not related to the phenotype as shown in a previous study (Collinge et al., 2021).

**Figure 3:**
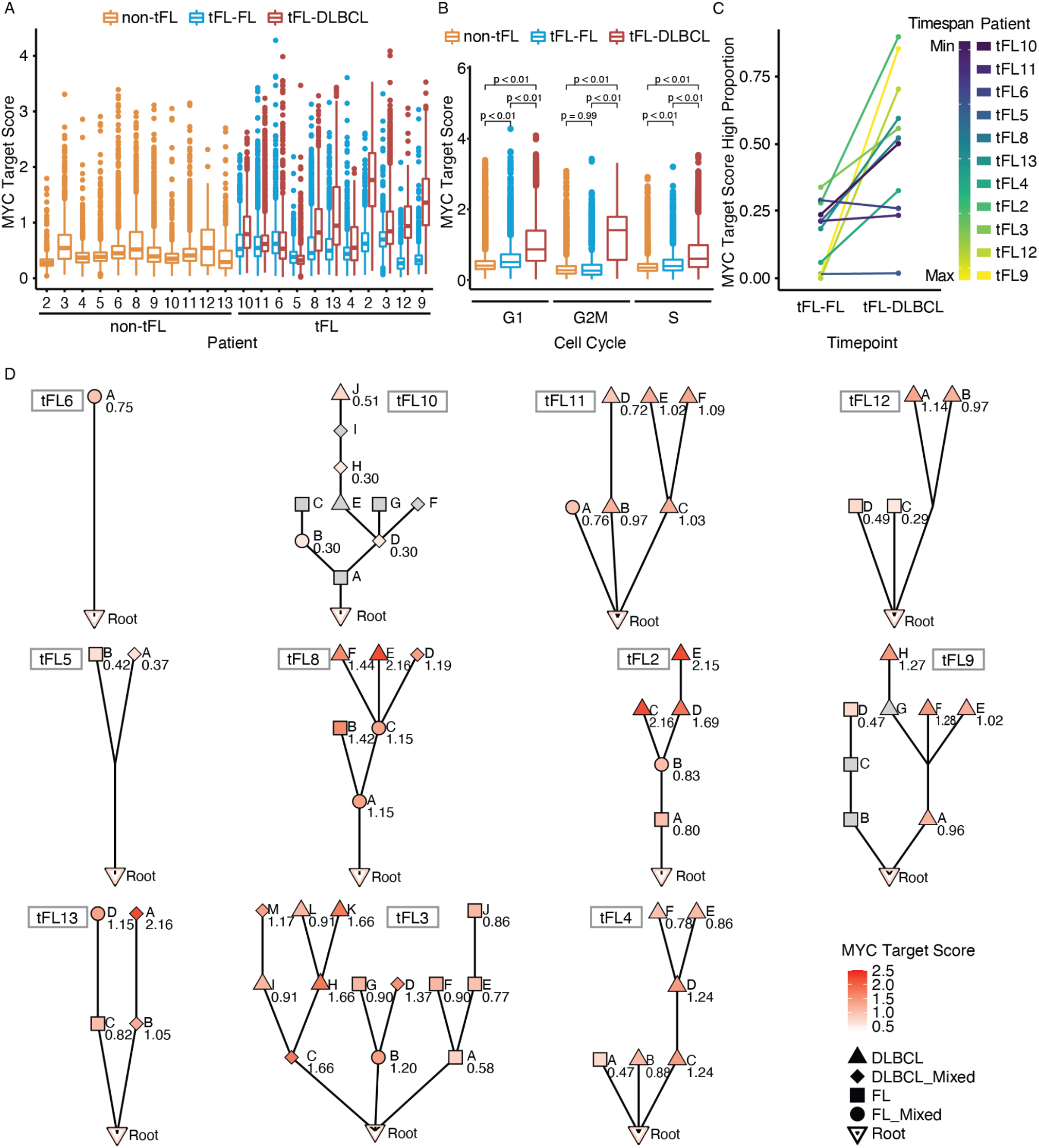
Simultaneous measurement of phenotypic and genomic evolution at the single cell level reveals the role of MYC pathway within the clonal structure of tFL. **Panel A**: Boxplot representing the MYC target score (MTS) within each sample: 11 non-tFL (yellow), 11 tFL-pairs (tFL-FL timepoint in blue and tFL-DLBCL in red). tFL-pairs are ordered based on time elapsed between tFL-FL and tFL-DLBCL biopsies. **Panel B:** Boxplot representing the MTS per cell cycle state (G1, G2/M, S) and within each timepoint. **Panel C:** Line plot representing the evolution of proportion of cells with a high MTS within each pair-timepoint, from tFL-FL to tFL-DLBCL. tFL-pairs are colored based on time elapsed between tFL-FL and tFL-DLBCL biopsies. **Panel D**: Genomic clones aligned with phenotypic status. Each sub-panel represents pair level genomic phylogenetic trees based on CNV (DLP) data and colored after clone-alignment based on the MYC target score (MTS) per clone. tFL-pairs are ordered as in Figure 2 panel A. Squares represent clones composed only of tFL-FL timepoint cells only, circles represent mixed-clones composed of a majority of tFL-FL cells and less tFL-DLBCL timepoint cells, triangles represent clones composed of only tFL-DLBCL timepoint cells and diamonds represent mixed clones composed of a majority of tFL-DLBCL cells with less tFL-FL timepoint cells. Dark red represents a high MTS, white a low MTS. Clones where clonealign failed to assign cells to 10X gene expression data (based on Supplementary Figure 11 B-C) were colored in gray.

Finally, non-tFL cells had lower mean MTS, mTORC1 and OXPHOS scores and a lower proportion of cells with a high score, as compared to tFL-FL cells, independently from cell cycle state (Figure 3B, Supplementary Figure 11A-C). However, the presence of some cells with high scores in these indolent cases suggest that other factors influence the final phenotype such as interactions between the tumor B cells and a more or less permissive TME in FL.

### Integrated scWGS and scWTS reveals dynamics of biological pathway regulation during evolution

We next explored the phenotypic evolution at the level of clones. As the scWTS and scWGS data was derived from the same cell suspensions for each case, we applied a computational method to assign single cell expression profiles to genomic clones (Campbell et al., 2019) (see method section, Supplementary Figure 12A). To validate the computational assignment of cells, we checked that the proportion of cells from the scWTS assigned to clones matched the inferred prevalences from scWGS (Supplementary Figure 12B-C). To assess the relationship between clonal evolution and phenotypic evolution, we calculated the median MTS for each DLP clone and represented the results in the context of the phylogenetic trees (Figure 3D). Strikingly, the tFL-DLBCL timepoint clones had a significantly higher MTS score than the one from tFL-FL biopsies (Supplementary Figure 13, Supplementary Table 5). Moreover, we observed an increase of the MTS along the DLBCL branches of the trees. This suggests that a MYC-related target signature is under ongoing selection within the clonal populations, and that the phenotypic transition during transformation relies, at least partially, on an ongoing genomic clonal evolution.

### Single cell analysis reveals the co-evolution of TME during transformation

Given that transformation largely appeared to be driven by clones which were not detectable at the time of FL diagnosis, we next sought to determine whether features of the TME may provide indications of transformation risk. To that end we performed reclustering of the scWTS data from the non-B TME cells from all 35 samples. We identified 18 clusters and annotated them using key marker genes as: T-cell (15 subsets), macrophage, NK, and plasmacytoid dendritic cell (Supplementary Figure 14, Supplementary Table 6). Further annotation, based on the expression of selected marker genes, allowed us to curate the 15 T cell clusters as naïve/central-memory-like (N=5, T_cm), exhausted/regulatory-like (N=4, T_exh), cytotoxic (N=2), T follicular helper-like (TFH-like, N=3), and a proliferation cluster (high Ki67 expression, N=1) (Supplementary Figure 15).

Single cell resolution allowed us to determine the evolution of the non-B cell TME composition (Figure 4) in the transformation pairs. We observed a consistent decrease in T_cm and TFH clusters, as opposed to an increase in T_exh and cytotoxic clusters during transformation (Figure 4, Supplementary Figure 16, 17). Interestingly, when adding the RLN and non-tFL data-set to this comparison, the decrease in T_cm clusters versus increase in T_exh clusters showed a continuous gradient from RLN to non-tFL, tFL-FL and tFL-DLBCL (Figure 4A, Supplementary Figure 17A). These variations were confirmed at the patient level (Figure 4B), and correlated with the time between tFL-FL and tFL-DLBCL biopsies (Supplementary Figure 18A-B). Moreover, the magnitude of the TME shift was correlated with the magnitude of the malignant B-cell phenotypic evolution (Supplementary Figure 18D-E), suggesting that the crosstalk between the TME and tumor B cell is also an important player in the transformation process.

**Figure 4:**
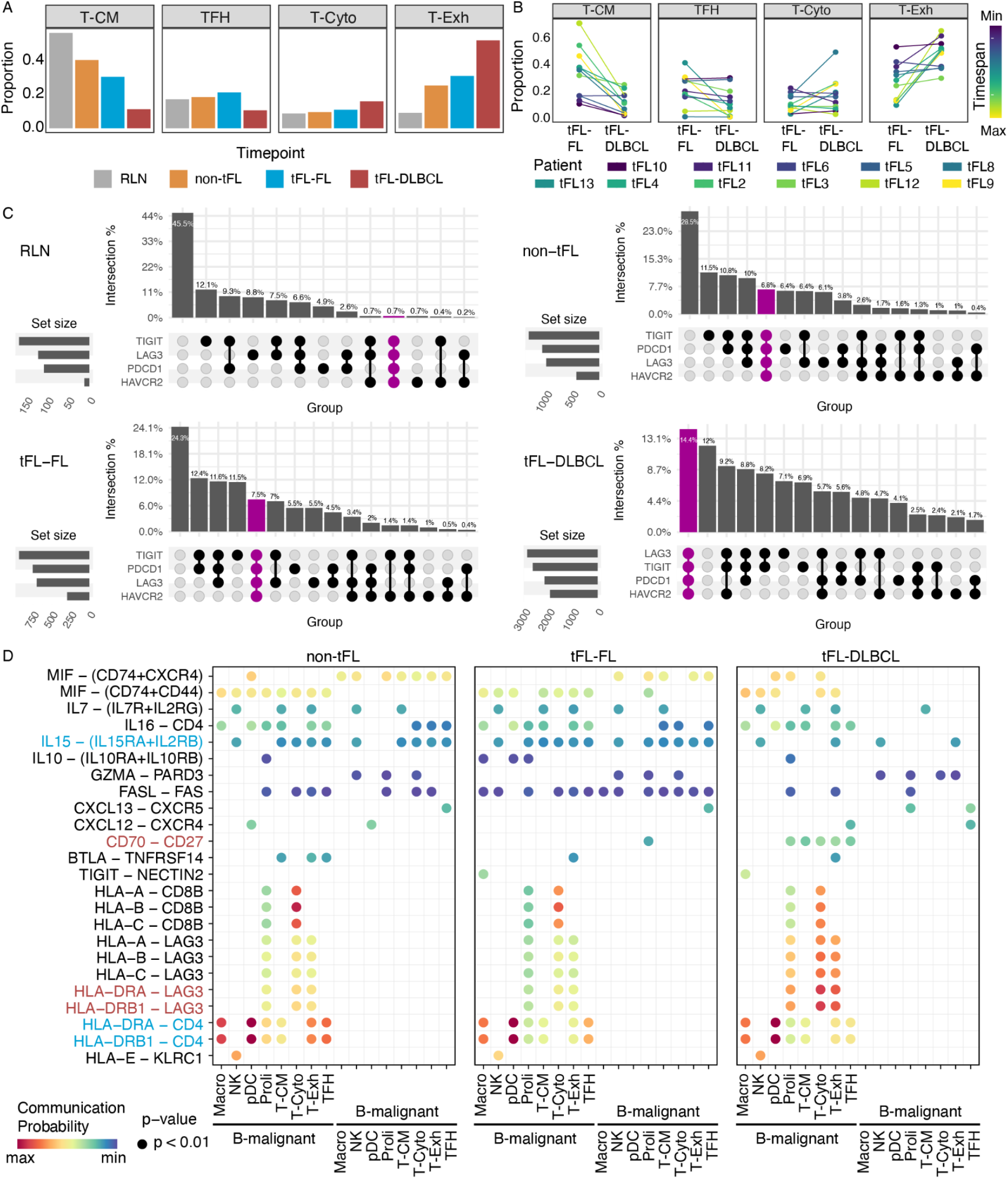
Tumor microenvironment evolution in the transformation process and its interaction with tumor B cells. **Panel A**: Barplot representing proportion of cells within each timepoint assigned to a TME cell type. Data are based on the UMAP representation of re-clustering of TME cells (ie non-B cells from the 2 RLN and non-B and non-tumoral cells from the 11 non-tFL, 11 tFL-pairs (11 tFL-FL and 11 tFL-DLBCL are included in this analysis). T-cm represent cells included in cluster with T central memory markers identified (ie cluster 4,6,8,10,16), TFH cells included in cluster with TFH markers identified (ie cluster 3,11,15), T-cyto cells included in cluster with T cytotoxic markers identified (ie cluster 5, 9), T-exh cells included in cluster with exhaustion or regulatory markers identified (ie cluster 1,2,7,13). Cells included in cluster 12 (characterized by high MKI67 expression level), NK (natural killer cells cluster 14), Macro (macrophage cells cluster 17) and pDC (plasmacytoid dendritic cells cluster 18) are shown in Supplementary Figure 16. **Panel B**: Line plots representing the evolution of the proportion of cells assigned to a cell type within tFL-FL and tFL-DLBCL timepoints. tFL-pairs are colored by time elapsed between tFL-FL and tFL-DLBCL timepoint. Cells included in cluster 12 (characterized by high MKI67 expression level), NK (natural killer cells cluster 14), Macro (macrophage cells cluster 17) and pDC (plasmacytoid dendritic cells cluster 18) are shown in Supplementary Figure 16. **Panel C:** Co-expression plot of regulatory/exhaustion marker genes based on 10X scRNA seq data. Within the TME clusters, CD8+ cells were selected, and double positive cells for CD8 and CD4 were removed. Data are shown in RLN (top), nontFL, tFL-FL and tFL-DLBCL (bottom) samples. **Panel D**: TME cell and malignant B cells interaction. Representation of the probability of interaction between tumor B cells and TME cell-types, within each cohort: non-tFL (left panel), tFL-FL (middle panel), tFL-DLBCL (right panel). Interaction probability has been represented for pathways of interest (left) and colored by the level of probability (higher with red dot and lower with blue dot, absence of interaction is represented by the absence of dot). Pathways enriched in tFL-DLBCLs samples are highlighted in red, those in FLs in blue. Abbreviations: Malig: malignant, Macro: macrophage, NK: natural killer, pDC: plasmacytoid dendritic cells, Proli: proliferation cluster, T-CM: T central memory cells clusters, T-cyto: T cytotoxic cells clusters, T-reg: T regulatory cells clusters, TFH: T-follicular helper cells clusters.

We next looked at the co-expression profile of exhaustion/regulatory genes. T cells from FL samples showed a distinct profile compared to T cells in the RLN (healthy controls) (Figure 4C, Supplementary Figure 19 and 20). The plots highlight a shift in the co-expression of *LAG3, TIGIT, PDCD1, HAVCR2* with a strong increase in the number and proportion of CD8 cells with a “full” exhausted phenotype in tFL-DLBCLs versus tFL-FLs. These cells are almost absent in RLN samples, and their proportion within CD8+ cells increased gradually from non-tFL to tFL-FL to tFL-DLBCL. At the patient level, the proportion of CD8 cells co-expressing *LAG3/PDCD1/TIGIT/HAVCR2* increased, except in the tFL-pair-5 and tFL-pair-6 (Supplementary Figure 21), consistent with the tumour B phenotype/genotype and pathological data and tFL-pair-4. tFL-pair-4 rather showed an increase in a regulatory cluster composed of classical CD4 T reg cells (Cluster 1 T_reg, Supplementary Figure 17B) suggesting a distinct process in this sample that appeared to be the one with the highest level of nodal evolution between tFL-FL and tFL-DLBCL timepoints.

### Tumor B and TME cell interactions reveal a regulatory eco-system

Using cell-cell interaction methods (Jin et al., 2021), we looked at the inferred communication between tumor B cells and TME cells, within the non-tFL, tFL-FL and tFL-DLBCL data-sets, based on co-evolutionary expression of ligands and receptors in selected pathways (Figure 4D). This analysis showed a similar pattern between non-tFL and tFL-FL, but highlighted striking dissimilarities with tFL-DLBCLs. The level of MHCII-CD4 interaction was lower in TFH cells from tFL-DLBCLs when compared to FLs. In addition, the IL15-IL15RA/IL2RB signalling axis between tumor B cells and T_cm, TFH or cytotoxic T clusters was present in non-tFL and tFL-FL samples but absent in tFL-DLBCLs. In contrast, even though the MHCI-CD8 interaction profile was very similar between the FLs and DLBCLs, we observed a strong enrichment in MHC(I/II)-LAG3 interaction in T-exh and T cytotoxic cell types in tFL-DLBCLs samples versus FLs, further emphasizing the role of the B and TME crosstalk during transformation. Furthermore, significant CD70-CD27 interactions were inferred in the tFL-DLBCL samples between tumor B cells and all TME T clusters, but not within tFL-FL or non-tFL samples. Importantly, in our scWTS data, *CD27* was higher in the cytotoxic-LAG3+ and T-exh clusters, than in T_cm clusters, suggesting a role of CD27-CD70 axis in T cell exhaustion in the context of transformation (Supplementary Figure 22). Moreover, almost all CD8+LAG3+ cells co-expressed CD27, with an increase in the proportion of triple positive cells within CD8 cells from RLN, to non-tFL, tFL-FL and tFL-DLBCL (Supplementary Figure 23A), alongside with an increase in CD70 expression on tumor B cells after transformation (Supplementary Figure 24).

### Spatial analysis confirms a shift in the TME-B crosstalk during transformation

To validate our predictions from disaggregated single cell analysis we performed multi-color immunofluorescence analysis (MC-IF). Our analysis confirmed the increase in the co-expression of LAG3 and PD1 on CD8 cells during transformation (Figure 5B-D, Supplementary Figure 25A-C), with a vast majority of LAG3 positive cells being PD1 positive as well. Of note, non-tFL samples had a significantly lower proportion of LAG3+ and LAG3+PD1+ CD8 cells as compared to tFL-FL samples, suggesting that the LAG3 proportion within the TME cells could be used as a predictive marker for transformation. A spatial analysis showed that CD8+LAG3+PD1+ cells were significantly closer to B cells than CD8+LAG3+PD1-cells (Supplementary Figure 26A). Furthermore, even though the B cells and not further subtyped CD8 cells were equidistant in tFL-FL and tFL-DLBCL samples, CD8+LAG3+ and CD8+LAG3+PD1+ cells tended to be closer to B cells after transformation than in FL samples (Supplementary Figure 26A). These cells, with an exhausted phenotype, were significantly enriched within the close neighborhood of B cells after transformation (defined as 125 microns) (Figure 5E, Supplementary Figure 26B). Our analysis also showed an increase in the proportion of CD4+LAG3+PD1 cells in the neighborhood of B cells (Supplementary Figure 27 and 28). By contrast, TFH cells were closer to B cells in the tFL-FL and in non-tFL samples as compared to tFL-DLBCL (Supplementary Figure 29), confirming the qualitative shift in the TME composition and tumor B cell crosstalk during transformation. As in the scWTS data, the majority of CD8+LAG3+ cells also expressed CD27 by IF and these cells were closer to CD70 expressing B cells than the CD27-population, and in a greater proportion in the tFL-DLBCL samples as compared to paired FL samples (Figure 5C, Supplementary Figure 30).

**Figure 5:**
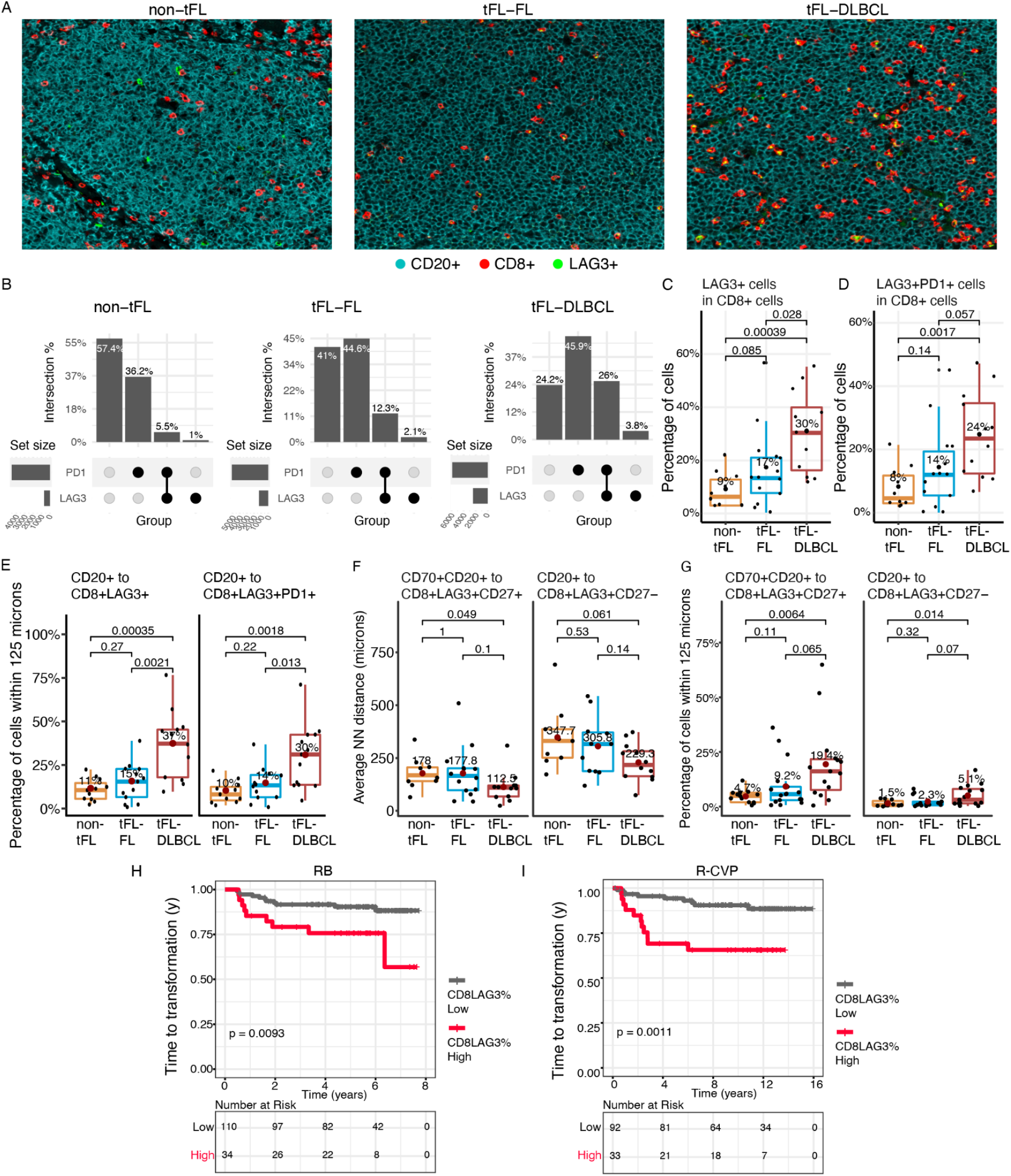
Validation of the tumor microenvironment shift using multicolor Immuno-histochemestry. This figure is based on multicolor Immunohistochemistry analysis, using the single-cell cohort (panel A-B-C-D) and validation cohort for survival analysis (panel H and I) **Panel A: Multi-color IHC** Picture highlighting CD20 cells (blue), CD8 cells (red) and LAG3+ cells (green) in an example of (from left to right) non-tFL, tFL-FL and tFL-DLBCL (Magnification 250X). **Panel B:** co-expression plot for LAG3 and PD1 expression, within CD8+ (and CD4-CD20-) cells, in MC-IHC in the 3 different cohorts (non-tFL, tFL-FL and tFL-DLBCL) **Panel C**: IHC expression of LAG3 in CD8+ cells (and CD4-CD20-) in the 3 different cohorts (non-tFL, tFL-FL and tFL-DLBCL). The proportion of LAG3 + cells among CD8+ cells is represented. **Panel D**: IHC expression of LAG3 and PD1 in CD8+ (and CD4-CD20-) cells in the 3 different cohorts (non-tFL, tFL-FL and tFL-DLBCL). The proportion of LAG3+PD1+co-expressing cells among CD8 + cells is represented. **Panel E**: Multicolor IHC neighborhood analysis. The first sub-panel represents the percentage of CD8+ cells that express LAG3 within 125 microns from CD20+ cells, in each timepoint. The second sub-panel represents the percentage of CD8+ cells that express LAG3 and PD1 within 125 microns from CD20+ cells. **Panel F:** Distance between CD8+LAG3+CD27+ and to CD20+CD70+ cells, and CD8+LAG3+CD27- to CD20+ cells in each timepoint. **Panel G:** The percentage of CD8+LAG3+CD27+ cells within 125 microns from CD70+CD20+ cells, and percentage of CD8+LAG3+CD27-cells within 125 microns from CD20+ cells, in each timepoint. **Panel H and I:** Transformation free survival curves in 2 independent cohorts of follicular lymphoma patients homogeneously treated with Rituximab Bendamustine (H) or R-CVP (I).

### Co-measurement of CD8 and LAG3 within the TME cell population defines a biomarker of transformation risk

Given the strong interaction observed between LAG3 and MHC-I within the cytotoxic and CD8-exh TME clusters and to assess the outcome predictive value of CD8+LAG3+ cell abundance, we used two independent cohorts of FL patients treated with rituximab-bendamustine (R-B) and R-CVP (cyclophosphamide, vincristine and prednisone), respectively. The percentage of CD8+LAG3+ out of CD20-cells was strongly associated with time to transformation in both R-B and R-CVP cohorts. When applying the threshold of the fourth quartile, patients with a high CD8+LAG3+ cell infiltrate had a shorter time to transformation, to progression and disease specific survival in both R-CVP and R-B cohorts (Figure 5G-H, Supplementary Figure 31-32, and Supplementary Table 7). Within a multivariate Cox model including FLIPI score and FL histological grading, CD8+LAG3+ infiltration alone remained significantly associated with transformation risk in both R-CVP and R-Benda cohorts (Supplementary Figure 33-34).

## Discussion

By applying high-dimensional scRNA and scDNA profiling techniques in paired FL and transformed biopsies, we were able to observe, for the first time, the relationships between cellular genotype and phenotype during transformation, and describe co-evolution of malignant cells within a qualitatively changing TME. We provide direct single cell evidence that genomic divergence, previously only inferred from bulk sequencing data (Green et al.,2013; Kridel et al., 2016; Okosun et al., 2014; Pasqualucci et al., 2014), co-develops with phenotypic divergence during transformation highlighting a MYC target signature in malignant B cells. As a specific feature of co-evolution, an exhausted cytotoxic T cell phenotype in the TME emerged as a potential biomarker predictive of transformation risk and survival outcomes.

Recently, single cell techniques have allowed a deeper characterization of FL tumor B cells, revealing strong inter-patient heterogeneity (Andor et al., 2019; Haebe et al., 2021; Han et al.,2022; Roider et al., 2020) and plasticity of tumor B cells resulting in loss of germinal center specific gene expression synchrony (Milpied et al., 2018). In contrast to our work, the topic of transformation was not addressed in these studies, which did not include pairs of pre- and post-transformation FL biopsies.

We observed different patterns and degrees of clonal divergence during transformation. Common patterns include the less frequent expansion of a precursor population present at the time of diagnosis with a transformed genotype and a more frequently observed emergence of new clones at transformation that were absent at diagnosis. In only a subset of cases, we were able to identify rare cells with a transformed genotype and phenotype in the pre transformed tFL-FL sample, up to 6 years before transformation.

Our single cell approach allowed us to study the biological pathways affected during the phenotypic transition from tFL-FL to tFL-DLBCL tumor B cells with unprecedented accuracy, which contrasts with previous bulk sequencing studies that could not reliably separate phenotypic measurements of malignant cells from TME components. Pathways reflecting tumor B cell autonomous survival, metabolism and proliferation, were enriched after transformation and importantly, the signal strength in MTS, OXPHOS, and mTOR signatures were independent of cell cycle state. This corroborates and provides additional context to previous work using unpaired bulk gene expression data highlighting expression signatures of embryonic stem cells that were not merely a proxy for proliferation differences between FL and transformed DLBCL (Gentles et al., 2009). Previously, DE analysis based on bulk sequencing, reported 2 different groups of transformed FL with opposite evolution of the expression level of *MYC* and its target genes (Lossos et al., 2002; Martinez-Climent et al.,2003). Pasqualucci et al reported *MYC* translocations and low-level copy number gains as enriched in transformed samples (Pasqualucci et al., 2014). Consistent with these data, we observed MYC rearrangements in the majority of the cases. However, the CNA were mostly gains that were already present at diagnosis (4/11). The new acquisition of a translocation during transformation was found in only one tFL-pair suggesting that alternative genetic and epigenetic mechanisms are driving a MYC target signature in the majority of cases.

The integration of genomic and transcriptomic information through the alignment of cells from scWTS and scWGS analysis, revealed the evolutionary dynamics of MYC target genes, OXPHOS as well as mTOCRC1 pathways during transformation, with an ongoing clonal selection process for clones with a high signature score. Furthermore, the level of phenotypic changes, and to a lesser extent genomic changes, were more related to time between the biopsies, than treatment received, meaning that transformation is a continuous process related to the ongoing acquisition of genomic aberrations driven by positive selection for a phenotype.

Our data also demonstrate that genomic features are not the only determinants of phenotypic evolution. Indeed, the observed correlation between the TME composition and B cell phenotypic shift suggests that a permissive TME in the vicinity of the tumor cells impacts fitness and survival, as seen in other lymphomas (Chiron et al., 2016; Mondello et al., 2021). The mechanistic role for escape of immune surveillance in the transformation process is supported in the literature by presence of mutations in genes with immune-related functions in tumours from tFL patients, such as *B2M (Kridel et al., 2016)*. In our data, a significant qualitative shift was observed within the overall T cell compartment, from cells with a TFH and central memory phenotype in FL samples, to cells with a cytotoxic, exhausted phenotype in transformed biopsies. More particularly, the CD8+LAG3+PD1+ population more than doubled during transformation and expanded with a fully exhausted phenotype and co-expression of *LAG3, PD1, TIGIT*, and *HAVCR2*. Notably, this exhausted population also expressed *CD27*, known to correlate with T regulatory activity (Duggleby et al., 2007). In particular, our cell to cell communication analysis provided additional evidence for specific interactor cell types and involved receptor-ligand pairs. These interactions include the enrichment of MHC-I/II-LAG3 and CD27-CD70 in DLBCL samples versus MCHII-CD4 in FL. The abundance and spatial relationship of cell types could be validated by imaging analysis, where compared to CD27-cells, CD8+LAG3+CD27+ cells were closer to CD70+ B cellsin tFL-DLBCL. The literature supports the role of the CD27-CD70 pathway in immune regulation, homeostasis and inhibitory function of Tregs, as well as CD8 and NK cell exhaustion (Bowakim et al., 2018; van Gisbergen et al., 2009; Tesselaar et al., 2003; Yang et al., 2007). Through its interaction with CD27, CD70 might function as an activator receptor on malignant B cells and an inhibitor of T cell response, with a reduction of apoptosis of Tregs in vivo (Claus et al., 2012). In NHL, CD70 has been associated with an immunosuppressive TME in mantle cell lymphoma (Balsas et al., 2021; Yang et al., 2007), and poor survival (Claus et al., 2012). Our data are consistent with the hypothesis that within ongoing clonal evolution, CD8 cells in the TME are triggered through the CD70-CD27 axis, providing proliferation and survival signal to the tumor B cell on the one side and an the acquisition of an exhausted phenotype for these CD8 cells on to the other side.

Single cell resolution afforded us the ability to identify phenotypic features that might provide a predictive value for transformation, such as MTS, mTORC1 or OXPHOS expression score. Furthermore, the magnitude of the TME/B interaction shift suggests that TME composition could provide more significant indication for the transformation risk as opposed to tumor B cell features. Using two independent cohorts, we could validate that the level of infiltration of LAG3+CD8+ cells before treatment initiation, within the TME, was a predictive marker of transformation and outcome. Other recent single-cell FL reports have also highlighted a key role of tumour microenvironment (TME) (Abe et al., 2022) B cell plasticity and attempted to develop a classification based on T cell subsets (Han et al., 2022), but our single cell study is the first one to show that implementation of single cell techniques in tFL prediction is valuable. The role of regulatory TME cells in transformation and outcome has already been assessed, primarily using bulk sequencing, and has resulted in inconsistent results on FL prognosis (Alcoceba et al., 2022; Casulo, 2021; Mondello et al., 2021). More recently, Tobin et al reported in bulk genomic studies that a low immune infiltration (characterized by the lower bulk expression of *LAG3, PD-1, FOX-P3* and *TIM-3)* was associated with an early progression in FL after R-CVP treatment (Tobin et al., 2019). High levels of PD1 cell infiltration was also associated with a better outcome in Carreras et al (Carreras et al., 2009), as well as CD8 cells infiltration (Wahlin et al., 2010). The single cell deconvolution of the TME components performed in our study, which provided a precise quantification and characterization of the different cell types, might explain the differences in these results.

Aiming to develop a predictive tFL tools, bulk sequencing studies have yielded little success (Crouch et al., 2021; Green et al., 2013; Kridel et al., 2016; Okosun et al., 2014; Pasqualucci et al., 2014) and our high precision single cell analysis could only identify cells with a transformed genotype in a subset of the pre-transformed FL biopsies. However, our observations suggest that a more promising avenue for tFL prediction might come from exploring phenotypic features of malignant cells and the composition and spatial architecture of the TME.

## Methods

### Patient selection

All samples for research were obtained according to protocols approved by the BCC Research Ethics Board.

For single cell analyses, paired pre- and post-transformation biopsies (pairs cohort) were identified from the BCC data-base where single cell suspensions were available at both time points. The control FL cohort (non-tFL cohort) was selected based on the absence of relapse/progression with more than 7 years of follow-up and availability of a frozen vial at FL diagnosis. All case materials were reviewed by expert hematopathologists to confirm the diagnosis of FL and transformation based on WHO criteria(Swerdlow et al., 2016). Samples were processed by batches (N=11) including one pair and one non-tFL sample (i.e. 3 samples per batch). A total of 11 pairs (FL and DLBCL timepoints, herein referred to as tFL-FL and tFL-DLBCL, respectively) and 11 non-tFL were included in the study, along with 2 reactive lymph node samples (RLN), leading to a total of 35 samples processed for 10X-5’ and BCR single cell RNAseq and 22 for single cell DNAseq using Direct Library Preparation plus platform (DLP+)(Swerdlow et al., 2016; Zahn et al., 2017) (processed for 11 pairs, not done for non-tFL and RLN).

We used tissue microarrays (TMA) from previously published FL cohorts homogeneously treated with rituximab bendamustine (BR) (Freeman et al., 2019) or R-CVP (Kridel et al., 2015) at BCC as validation cohorts (see Supplementary methods section).

### Library preparation for scDNAseq and scRNAseq: see Supplementary method section

For scDNA and scRNA-seq Sample Preparation, see Supplementary section.

For scDNAseq, DLP+ library construction was carried out as previously described (Zahn et al., 2017). Libraries were sequenced at UBC Biomedical Research Centre (BRC) in Vancouver, British Columbia on the Illumina NextSeq 550 (mid- or high-output, paired-end 150-bp reads).

For scRNAseq, in total 7,000 cells per sample were loaded into a Chromium Chip A (PN-1000009) and processed according to the Chromium Single Cell V(D)J Reagent Kit User Guide. 5’Gene expression libraries from 2 samples were pooled and sequenced in a HiSeq2500 instrument (Recipe: 125Cycles Read1, 8 cycles index, 125 cycles Read2). BCR libraries from 18 or 20 libraries were pooled and sequenced in a NextSeq 550 instrument (High Output Flow Cell V2.5 Recipe: 150 cycles Read1, 8 cycles index, 150 cycles Read2).

### Data processing of scDNA-seq and scRNA-seq: see Supplementary method section

Briefly, scDNAseq data were run through our automated pipeline (https://svn.bcgsc.ca/bitbucket/projects/SC/repos/single_cell_pipeline/browse). For scRNA-seq data, BaseCall files after sequencing were used to generate library-specific FASTQ files with cellranger mkfastq (v3.0.2).

#### Phylogenetic analysis for scDNA-seq

The copy number information output from HMMcopy was used to construct the phylogenetic trees with a single cell Bayesian tree reconstruction method called sitka(Salehi et al., 2021).

#### Dimension reduction, clustering analysis for scRNA-seq and differential expression analysis

Normalized log counts for the genes with biological variance >= 0 were used as input into the scanorama R package (Version 1.6) to perform batch correction. Principal components analysis was performed on the resulting batch-corrected expression matrix for the top 1000 most variable genes. The first 50 PCs were used as input for UMAP. Initial and subsequent unsupervised clustering was performed with monocle3 (Version 0.2.3.0). Clusters from PhenoGraph were manually assigned to a cell type by comparing the mean expression of known markers across cells in a cluster. Markers used for cell type annotation are described in Supplementary Table 8. Entropy was computed at cluster or timepoint levels to assess heterogeneity (see Supplementary method section).

#### Association of scDNAseq and scRNAseq

The malignant B cells identified during the clustering and annotation of scRNA-seq data were assigned to the genomic clones defined by copy number information from scDNA-seq data using a statistical method called clonealign(Campbell et al., 2019).

### TMA, FISH and IHC/multicolor-ImmunoFluorescence (MC-IF) analysis

Duplicate 1 mm FFPE cores were used to construct TMA including the 11 tFL-FL and tFL-DLBCL paired biopsies and the 11 non-tFL samples. IHC, IFand FISH were performed.

**Table 1.** clinical and treatment characteristics

|  | PAIRS, N=11 | NON-TFL, N=11 |
| --- | --- | --- |
| Mean Age at diagnosis | 60y/o (34-76) | 59y/o (49-76) |
| Gender | 4F / 7M | 3F / 8M |
| FL grade at diagnosis: |  |  |
| 1. Grade 1-2 | 7 | 9 |
| 2. Grade 3A | 4 | 2 |
| FL biopsy |  |  |
| -diagnostic/before any treatment | 9 | 11 |
| -first relapse after: |  | 0 |
| RAD | 1 |  |
| Chemo | 1 |  |
| DLBCL biopsy |  | NA |
| -first occurrence of DLBCL | 11 |  |
| -second or more | 0 |  |
| DLBCL Biopsy site different than FL | 5/11 | NA |
| B symptoms at FL diagnosis: | /11 | /10 |
| Yes | 2 | 1 |
| No | 9 | 9 |
| Hemoglobin <120g at FL diagnosis | 0/10 | 0/10 |
| LDH>UNL | 0/8 | 0/9 |
| AA stage at FL diagnosis |  |  |
| -1-2 | 5 | 4 |
| -3-4 | 6 | 7 |
| Treatment at FL diagnosis |  |  |
| -Watch and Wait | 5 | 3 |
| -Radiation | 2 | 2 |
| -surgery | 1 | 0 |
| -Rituximab only | 0 | 2 |
| -Rituximab-chemotherapy | 3 | 4 |
| Relapse |  |  |
| -No | 0 | 11 |
| -Yes | 11 | 0 |
| Chemotherapy before transformation |  | NA |
| -no | 5 |  |
| -yes | 6 |  |
| Timing between FL-DLBCL biopsy |  | NA |
| -Mean (range) | 41m (9-114) |  |
| -<24 months | 5 |  |
| ->=24 months | 6 |  |

## Acknowledgements

This study is supported by Program Project Grant funding from the Terry Fox Research Institute (Grant No. 1061 and 1108), Large Scale Applied Research Project funding from Genome Canada (Grant No. 13124), Genome BC (Grant No. 271LYM) and CIHR (Grant No. GP1-155873), a Foundation grant from CIHR (Grant No. 148393), the BC Cancer Foundation and the Paul G. Allen Frontiers Group (Distinguished Investigator award to C.S. Grant No. 12829). In addition, AR receives operating funds from the Natural Sciences and Engineering Research Council of Canada (Grant RGPIN-2022-04378), the V Foundation (Grant V2021-033), and Michael Smith Health Research BC (Grant SCH-2019-2100). We also thank Teva and the LRF team for funding the RB cohort. We acknowledge that the graphical abstract is created with BioRender.com.

## Author contributions

Cohort Design, C.Sa., T.A., K.T; Data Generation, C.Sa., K.T., P.F., S.B.N, T.R; Data Preprocessing, S.W., J.C., D.L., C.St. Data Analysis, C.Sa., S.W., J.C; Manuscript Drafting, C.Sa., S.W; Manuscript Review & Editing, C.Sa., S.W., A.R., C.S., J.C., B.N., A.W., D.S; Supervision, C.S., A.R; Funding Acquisition, C.S., A.R.

## References

Abe, Y., Sakata-Yanagimoto, M., Fujisawa, M., Miyoshi, H., Suehara, Y., Hattori, K., Kusakabe, M., Sakamoto, T., Nishikii, H., Nguyen, T.B., et al. (2022). A single-cell atlas of non-haematopoietic cells in human lymph nodes and lymphoma reveals a landscape of stromal remodelling. Nat. Cell Biol. 24, 565–578..

Alcoceba, M., García-Álvarez, M., Okosun, J., Ferrero, S., Ladetto, M., Fitzgibbon, J., and García-Sanz, R. (2022). Genetics of Transformed Follicular Lymphoma. Hematology 3, 615–633.

Al-Tourah, A.J., Gill, K.K., Chhanabhai, M., Hoskins, P.J., Klasa, R.J., Savage, K.J., Sehn, L.H., Shenkier, T.N., Gascoyne, R.D., and Connors, J.M. (2008). Population-based analysis of incidence and outcome of transformed non-Hodgkin’s lymphoma. J. Clin. Oncol. 26, 5165–5169..

Andor, N., Simonds, E.F., Czerwinski, D.K., Chen, J., Grimes, S.M., Wood-Bouwens, C., Zheng, G.X.Y., Kubit, M.A., Greer, S., Weiss, W.A., et al. (2019). Single-cell RNA-Seq of follicular lymphoma reveals malignant B-cell types and coexpression of T-cell immune checkpoints. Blood 133, 1119–1129..

Bachy, E., Seymour, J.F., Feugier, P., Offner, F., López-Guillermo, A., Belada, D., Xerri, L., Catalano, J.V., Brice, P., Lemonnier, F., et al. (2019). Sustained Progression-Free Survival Benefit of Rituximab Maintenance in Patients With Follicular Lymphoma: Long-Term Results of the PRIMA Study. J. Clin. Oncol. 37, 2815–2824..

Balsas, P., Veloza, L., Clot, G., Sureda-Gómez, M., Rodríguez, M.-L., Masaoutis, C., Frigola, G., Navarro, A., Beà, S., Nadeu, F., et al. (2021). SOX11, CD70, and Treg cells configure the tumor-immune microenvironment of aggressive mantle cell lymphoma. Blood 138, 2202–2215..

Bolen, C.R., Mattiello, F., Herold, M., Hiddemann, W., Huet, S., Klapper, W., Marcus, R., Mir, F., Salles, G., Weigert, O., et al. (2021). Treatment dependence of prognostic gene expression signatures in de novo follicular lymphoma. Blood 137, 2704–2707..

Bouska, A., McKeithan, T.W., Deffenbacher, K.E., Lachel, C., Wright, G.W., Iqbal, J., Smith, L.M., Zhang, W., Kucuk, C., Rinaldi, A., et al. (2014). Genome-wide copy-number analyses reveal genomic abnormalities involved in transformation of follicular lymphoma. Blood 123, 1681–1690..

Bouska, A., Zhang, W., Gong, Q., Iqbal, J., Scuto, A., Vose, J., Ludvigsen, M., Fu, K., Weisenburger, D.D., Greiner, T.C., et al. (2017). Combined copy number and mutation analysis identifies oncogenic pathways associated with transformation of follicular lymphoma. Leukemia 31, 83–91..

Bowakim, N., Acolty, V., Dhainaut, M., Yagita, H., Oldenhove, G., Leo, O., and Moser, M. (2018). Role of the CD27/CD70 pathway in regulatory T cell function. The Journal of Immunology 200, 47.27–47.27..

Brodtkorb, M., Lingjærde, O.C., Huse, K., Trøen, G., Hystad, M., Hilden, V.I., Myklebust, J.H., Leich, E., Rosenwald, A., Delabie, J., et al. (2014). Whole-genome integrative analysis reveals expression signatures predicting transformation in follicular lymphoma. Blood 123, 1051–1054..

Campbell, K.R., Steif, A., Laks, E., Zahn, H., Lai, D., McPherson, A., Farahani, H., Kabeer, F., O’Flanagan, C., Biele, J., et al. (2019). clonealign: statistical integration of independent single-cell RNA and DNA sequencing data from human cancers. Genome Biol. 20, 54..

Carlotti, E., Wrench, D., Matthews, J., Iqbal, S., Davies, A., Norton, A., Hart, J., Lai, R., Montoto, S., Gribben, J.G., et al. (2009). Transformation of follicular lymphoma to diffuse large B-cell lymphoma may occur by divergent evolution from a common progenitor cell or by direct evolution from the follicular lymphoma clone. Blood 113, 3553–3557..

Carreras, J., Lopez-Guillermo, A., Fox, B.C., Colomo, L., Martinez, A., Roncador, G., Montserrat, E., Campo, E., and Banham, A.H. (2006). High numbers of tumor-infiltrating FOXP3-positive regulatory T cells are associated with improved overall survival in follicular lymphoma. Blood 108, 2957–2964..

Carreras, J., Lopez-Guillermo, A., Roncador, G., Villamor, N., Colomo, L., Martinez, A., Hamoudi, R., Howat, W.J., Montserrat, E., and Campo, E. (2009). High numbers of tumor-infiltrating programmed cell death 1-positive regulatory lymphocytes are associated with improved overall survival in follicular lymphoma. J. Clin. Oncol. 27, 1470–1476..

Casulo, C. (2021). Upfront identification of high-risk follicular lymphoma. Hematol. Oncol. 39 Suppl 1, 88–93..

Chiron, D., Bellanger, C., Papin, A., Tessoulin, B., Dousset, C., Maiga, S., Moreau, A., Esbelin, J., Trichet, V., Chen-Kiang, S., et al. (2016). Rational targeted therapies to overcome microenvironment-dependent expansion of mantle cell lymphoma. Blood 128, 2808–2818..

Claus, C., Riether, C., Schürch, C., Matter, M.S., Hilmenyuk, T., and Ochsenbein, A.F. (2012). CD27 signaling increases the frequency of regulatory T cells and promotes tumor growth. Cancer Res. 72, 3664–3676..

Collinge, B., Ben-Neriah, S., Chong, L., Boyle, M., Jiang, A., Miyata-Takata, T., Farinha, P., Craig, J.W., Slack, G.W., Ennishi, D., et al. (2021). The impact of MYC and BCL2 structural variants in tumors of DLBCL morphology and mechanisms of false-negative MYC IHC. Blood 137, 2196–2208..

Crouch, S., Painter, D., Barrans, S., Roman, E., Beer, P., Lacy, S., Cooke, S., Webster, N., Glover, P., Hoppe, S., et al. (2021). MOLECULAR SUBCLUSTERS OF FOLLICULAR LYMPHOMA: A REPORT FROM THE UK’S HAEMATOLOGICAL MALIGNANCY RESEARCH NETWORK. Hematological Oncology 39. https://doi.org/10.1002/hon.40_2879.

Dave, S.S., Wright, G., Tan, B., Rosenwald, A., Gascoyne, R.D., Chan, W.C., Fisher, R.I., Braziel, R.M., Rimsza, L.M., Grogan, T.M., et al. (2004). Prediction of survival in follicular lymphoma based on molecular features of tumor-infiltrating immune cells. N. Engl. J. Med. 351, 2159–2169..

Davies, A.J., Rosenwald, A., Wright, G., Lee, A., Last, K.W., Weisenburger, D.D., Chan, W.C., Delabie, J., Braziel, R.M., Campo, E., et al. (2007). Transformation of follicular lymphoma to diffuse large B-cell lymphoma proceeds by distinct oncogenic mechanisms. Br. J. Haematol. 136, 286–293..

Duggleby, R.C., Shaw, T.N.F., Jarvis, L.B., Kaur, G., and Gaston, J.S.H. (2007). CD27 expression discriminates between regulatory and non-regulatory cells after expansion of human peripheral blood CD4+ CD25+ cells. Immunology 121, 129–139..

Elenitoba-Johnson, K.S.J., Jenson, S.D., Abbott, R.T., Palais, R.A., Bohling, S.D., Lin, Z., Tripp, S., Shami, P.J., Wang, L.Y., Coupland, R.W., et al. (2003). Involvement of multiple signaling pathways in follicular lymphoma transformation: p38-mitogen-activated protein kinase as a target for therapy. Proc. Natl. Acad. Sci. U. S. A. 100, 7259–7264..

Farinha, P., Al-Tourah, A., Gill, K., Klasa, R., Connors, J.M., and Gascoyne, R.D. (2010). The architectural pattern of FOXP3-positive T cells in follicular lymphoma is an independent predictor of survival and histologic transformation. Blood 115, 289–295..

Federico, M., Caballero Barrigón, M.D., Marcheselli, L., Tarantino, V., Manni, M., Sarkozy, C., Alonso-Álvarez, S., Wondergem, M., Cartron, G., Lopez-Guillermo, A., et al. (2018). Rituximab and the risk of transformation of follicular lymphoma: a retrospective pooled analysis. Lancet Haematol 5, e359–e367..

Freeman, C.L., Kridel, R., Moccia, A.A., Savage, K.J., Villa, D.R., Scott, D.W., Gerrie, A.S., Ferguson, D., Cafferty, F., Slack, G.W., et al. (2019). Early progression after bendamustine-rituximab is associated with high risk of transformation in advanced stage follicular lymphoma. Blood 134, 761–764..

Gentles, A.J., Alizadeh, A.A., Lee, S.-I., Myklebust, J.H., Shachaf, C.M., Shahbaba, B., Levy, R., Koller, D., and Plevritis, S.K. (2009). A pluripotency signature predicts histologic transformation and influences survival in follicular lymphoma patients. Blood 114, 3158–3166..

van Gisbergen, K.P.J.M., van Olffen, R.W., van Beek, J., van der Sluijs, K.F., Arens, R., Nolte, M.A., and van Lier, R.A. (2009). Protective CD8 T cell memory is impaired during chronic CD70-driven costimulation. J. Immunol. 182, 5352–5362..

Glas, A.M., Knoops, L., Delahaye, L., Kersten, M.J., Kibbelaar, R.E., Wessels, L.A., van Laar, R., van Krieken, J.H.J.M., Baars, J.W., Raemaekers, J., et al. (2007). Gene-expression and immunohistochemical study of specific T-cell subsets and accessory cell types in the transformation and prognosis of follicular lymphoma. J. Clin. Oncol. 25, 390–398..

Green, M.R. (2018). Chromatin modifying gene mutations in follicular lymphoma. Blood 131, 595–604..

Green, M.R., Gentles, A.J., Nair, R.V., Irish, J.M., Kihira, S., Liu, C.L., Kela, I., Hopmans, E.S., Myklebust, J.H., Ji, H., et al. (2013). Hierarchy in somatic mutations arising during genomic evolution and progression of follicular lymphoma. Blood 121, 1604–1611..

Haebe, S., Shree, T., Sathe, A., Day, G., Czerwinski, D.K., Grimes, S.M., Lee, H., Binkley, M.S., Long, S.R., Martin, B., et al. (2021). Single-cell analysis can define distinct evolution of tumor sites in follicular lymphoma. Blood 137, 2869–2880..

Han, G., Deng, Q., Marques-Piubelli, M.L., Dai, E., Dang, M., Ma, M.C.J., Li, X., Yang, H., Henderson, J., Kudryashova, O., et al. (2022). Follicular Lymphoma Microenvironment Characteristics Associated with Tumor Cell Mutations and MHC Class II Expression. Blood Cancer Discov 3, 428–443..

Huet, S., Tesson, B., Jais, J.-P., Feldman, A.L., Magnano, L., Thomas, E., Traverse-Glehen, A., Albaud, B., Carrère, M., Xerri, L., et al. (2018). A gene-expression profiling score for prediction of outcome in patients with follicular lymphoma: a retrospective training and validation analysis in three international cohorts. Lancet Oncol. 19, 549–561..

Jin, S., Guerrero-Juarez, C.F., Zhang, L., Chang, I., Ramos, R., Kuan, C.-H., Myung, P., Plikus, M.V., and Nie, Q. (2021). Inference and analysis of cell-cell communication using CellChat. Nat. Commun. 12, 1088..

Jurinovic, V., Kridel, R., Staiger, A.M., Szczepanowski, M., Horn, H., Dreyling, M.H., Rosenwald, A., Ott, G., Klapper, W., Zelenetz, A.D., et al. (2016). Clinicogenetic risk models predict early progression of follicular lymphoma after first-line immunochemotherapy. Blood 128, 1112–1120..

Kridel, R., Xerri, L., Gelas-Dore, B., Tan, K., Feugier, P., Vawda, A., Canioni, D., Farinha, P., Boussetta, S., Moccia, A.A., et al. (2015). The Prognostic Impact of CD163-Positive Macrophages in Follicular Lymphoma: A Study from the BC Cancer Agency and the Lymphoma Study Association. Clin. Cancer Res. 21, 3428–3435..

Kridel, R., Chan, F.C., Mottok, A., Boyle, M., Farinha, P., Tan, K., Meissner, B., Bashashati, A., McPherson, A., Roth, A., et al. (2016). Histological Transformation and Progression in Follicular Lymphoma: A Clonal Evolution Study. PLoS Med. 13, e1002197..

Laks, E., McPherson, A., Zahn, H., Lai, D., Steif, A., Brimhall, J., Biele, J., Wang, B., Masud, T., Ting, J., et al. (2019). Clonal Decomposition and DNA Replication States Defined by Scaled Single-Cell Genome Sequencing. Cell 179, 1207–1221.e22..

Lossos, I.S., Alizadeh, A.A., Diehn, M., Warnke, R., Thorstenson, Y., Oefner, P.J., Brown, P.O., Botstein, D., and Levy, R. (2002). Transformation of follicular lymphoma to diffuse large-cell lymphoma: Alternative patterns with increased or decreased expression of c - myc and its regulated genes. Proceedings of the National Academy of Sciences 99, 8886–8891. https://doi.org/10.1073/pnas.132253599.

Martinez-Climent, J.A., Alizadeh, A.A., Segraves, R., Blesa, D., Rubio-Moscardo, F., Albertson, D.G., Garcia-Conde, J., Dyer, M.J.S., Levy, R., Pinkel, D., et al. (2003). Transformation of follicular lymphoma to diffuse large cell lymphoma is associated with a heterogeneous set of DNA copy number and gene expression alterations. Blood 101, 3109–3117.

Milpied, P., Cervera-Marzal, I., Mollichella, M.-L., Tesson, B., Brisou, G., Traverse-Glehen, A., Salles, G., Spinelli, L., and Nadel, B. (2018). Human germinal center transcriptional programs are de-synchronized in B cell lymphoma. Nature Immunology 19, 1013–1024. https://doi.org/10.1038/s41590-018-0181-4.

Mondello, P., Fama, A., Larson, M.C., Feldman, A.L., Villasboas, J.C., Yang, Z.-Z., Galkin, I., Svelolkin, V., Postovalova, E., Bagaev, A., et al. (2021). Lack of intrafollicular memory CD4 + T cells is predictive of early clinical failure in newly diagnosed follicular lymphoma. Blood Cancer J. 11, 130..

Okosun, J., Bödör, C., Wang, J., Araf, S., Yang, C.-Y., Pan, C., Boller, S., Cittaro, D., Bozek, M., Iqbal, S., et al. (2014). Integrated genomic analysis identifies recurrent mutations and evolution patterns driving the initiation and progression of follicular lymphoma. Nat. Genet. 46, 176–181..

Pasqualucci, L., Khiabanian, H., Fangazio, M., Vasishtha, M., Messina, M., Holmes, A.B., Ouillette, P., Trifonov, V., Rossi, D., Tabbò, F., et al. (2014). Genetics of follicular lymphoma transformation. Cell Rep. 6, 130–140..

Roider, T., Seufert, J., Uvarovskii, A., Frauhammer, F., Bordas, M., Abedpour, N., Stolarczyk, M., Mallm, J.-P., Herbst, S.A., Bruch, P.-M., et al. (2020). Dissecting intratumour heterogeneity of nodal B-cell lymphomas at the transcriptional, genetic and drug-response levels. Nat. Cell Biol. 22, 896–906..

Salehi, S., Dorri, F., Chern, K., Kabeer, F., Rusk, N., Funnell, T., Williams, M.J., Lai, D., Andronescu, M., Campbell, K.R., et al. (2021). Cancer phylogenetic tree inference at scale from 1000s of single cell genomes.

Sarkozy, C., Trneny, M., Xerri, L., Wickham, N., Feugier, P., Leppa, S., Brice, P., Soubeyran, P., Gomes Da Silva, M., Mounier, C., et al. (2016). Risk Factors and Outcomes for Patients With Follicular Lymphoma Who Had Histologic Transformation After Response to First-Line Immunochemotherapy in the PRIMA Trial. J. Clin. Oncol. 34, 2575–2582..

Sarkozy, C., Maurer, M.J., Link, B.K., Ghesquieres, H., Nicolas, E., Thompson, C.A., Traverse-Glehen, A., Feldman, A.L., Allmer, C., Slager, S.L., et al. (2019). Cause of Death in Follicular Lymphoma in the First Decade of the Rituximab Era: A Pooled Analysis of French and US Cohorts. Journal of Clinical Oncology 37, 144–152. https://doi.org/10.1200/jco.18.00400.

Subramanian, A., Tamayo, P., Mootha, V.K., Mukherjee, S., Ebert, B.L., Gillette, M.A., Paulovich, A., Pomeroy, S.L., Golub, T.R., Lander, E.S., et al. (2005). Gene set enrichment analysis: a knowledge-based approach for interpreting genome-wide expression profiles. Proc. Natl. Acad. Sci. U. S. A. 102, 15545–15550..

Swerdlow, S.H., Campo, E., Pileri, S.A., Harris, N.L., Stein, H., Siebert, R., Advani, R., Ghielmini, M., Salles, G.A., Zelenetz, A.D., et al. (2016). The 2016 revision of the World Health Organization classification of lymphoid neoplasms. Blood 127, 2375–2390. https://doi.org/10.1182/blood-2016-01-643569.

Tesselaar, K., Arens, R., van Schijndel, G.M.W., Baars, P.A., van der Valk, M.A., Borst, J., van Oers, M.H.J., and van Lier, R.A.W. (2003). Lethal T cell immunodeficiency induced by chronic costimulation via CD27-CD70 interactions. Nat. Immunol. 4, 49–54..

Tobin, J.W.D., Keane, C., Gunawardana, J., Mollee, P., Birch, S., Hoang, T., Lee, J., Li, L., Huang, L., Murigneux, V., et al. (2019). Progression of Disease Within 24 Months in Follicular Lymphoma Is Associated With Reduced Intratumoral Immune Infiltration. J. Clin. Oncol. 37, 3300–3309..

Tzankov, A., Meier, C., Hirschmann, P., Went, P., Pileri, S.A., and Dirnhofer, S. (2008). Correlation of high numbers of intratumoral FOXP3+ regulatory T cells with improved survival in germinal center-like diffuse large B-cell lymphoma, follicular lymphoma and classical Hodgkin’s lymphoma. Haematologica 93, 193–200..

Wagner-Johnston, N.D., Link, B.K., Byrtek, M., Dawson, K.L., Hainsworth, J., Flowers, C.R., Friedberg, J.W., and Bartlett, N.L. (2015). Outcomes of transformed follicular lymphoma in the modern era: a report from the National LymphoCare Study (NLCS). Blood 126, 851–857. https://doi.org/10.1182/blood-2015-01-621375.

Wahlin, B.E., Aggarwal, M., Montes-Moreno, S., Gonzalez, L.F., Roncador, G., Sanchez-Verde, L., Christensson, B., Sander, B., and Kimby, E. (2010). A unifying microenvironment model in follicular lymphoma: outcome is predicted by programmed death-1--positive, regulatory, cytotoxic, and helper T cells and macrophages. Clin. Cancer Res. 16, 637–650..

Yang, Z.-Z., Novak, A.J., Ziesmer, S.C., Witzig, T.E., and Ansell, S.M. (2007). CD70+ non-Hodgkin lymphoma B cells induce Foxp3 expression and regulatory function in intratumoral CD4+CD25 T cells. Blood 110, 2537–2544..

Zahn, H., Steif, A., Laks, E., Eirew, P., VanInsberghe, M., Shah, S.P., Aparicio, S., and Hansen, C.L. (2017). Scalable whole-genome single-cell library preparation without preamplification. Nat. Methods 14, 167–173..

