## Supplemental Information for "Integrated single cell analysis reveals co-evolution of malignant B cells and the tumor microenvironment in transformed follicular lymphoma"

### Supplementary methods

#### Cohorts

For the single cell analysis, including a cohort of transformed FL (PAIRS cohrt) and another one of non-transformed FL (non-tFL cohort), samples were selected from BCC data-base where single cell suspensions generated by mechanical dissociation were stained with the clinical diagnostic flow cytometry panel, and any excess remaining cell suspension material was frozen with DMSO cryoprotectant. Selection was done based on availability of frozen vials with at least 3 millions viably cryopreserved cells at time of FL diagnosis and at time of transformation. The control FL cohort (non-tFL cohort) was selected based on the absence of relapse/progression with more than 7 years of follow-up and availability of a frozen vial at FL diagnosis. All the samples were reviewed by hemato-pathology experts to confirm the diagnosis of FL and transformation based on WHO criteria. Samples were processed by batches (N=11) including one pair and one non-tFL sample (i.e. 3 samples per batch). A total of 11 pairs (FL and DLBCL timepoints, called tFL-FL and tFL-DLBCL in the manuscript) and 11 non-tFL were included in the study, along with 2 reactive lymph nodes samples (RLN, processed in independent batches) leading to a total of 35 samples processed for 10X-5' and BCR single cell RNAseq and 20 for single cell DNAseq using Direct Library Preparation plus platform (DLP+)(Zahn et al., 2017) (processed for 11 pairs, not done for non-tFL and RLN). WGS of paired normal DNA (extracted from non-tumoral PBMC) was performed for 10 out of the 11 tFL patients (no normal DNA available for PAIR2).

#### Validation cohorts

We enrolled previously published follicular lymphoma cohorts from the BCC to study transformation and outcome prediction based on TME features (Supplementary Table 9 and 10). R-CVP TMA (N=125 extracted from (Kridel et al., 2015)), initially treated with R-CVP and with a TMA available, including 20 pre-transformed biopsies (Supplementary Table 9). R-Bendamustine FL TMA (N=144 extracted from (Freeman et al., 2019)) treated with R-Bendamustin and TMA available, including 21 pre-transformed biopsies (Supplementary Table 10).

#### scDNA and scRNA-seq Sample Preparation

Cell suspensions from tumors or RLN were rapidly defrosted at 37°C, washed in 10 mL of RPMI-1640/10% fetal bovine serum (FBS) solution or RPMI 1640/20% FBS solution containing DNase I (Millipore Sigma) and washed in PBS. Cells were resuspended in PBC containing 2% FBS and stained with DAPI for 15 minutes at 4°C in the dark. We used DAPI (Sigma-Aldrich) for live–dead cells discrimination. Viable cells (DAPI-negative) were sorted on a FACS ARIAIII or FACS Fusion (BD Biosciences) using an 85-μm nozzle. Sorted cells were collected in 0.5 mL of medium, centrifuged and diluted in 1× PBS with 0.04% bovine serum albumin (BSA) (Supplementary Figure 1 for flow gating strategy). Cell number was determined using a Countess II Automated Cell Counter whenever possible. For DLP-DNA-seq, tFL-FL and tFL-DLBCL samples were sorted to isolate tumor cells based on tumor cell characteristics defined in the flow clinical report (using kappa/lambda, CD10, CD20, CD3 staining).

#### Library preparation for scDNAseq and scRNAseq

For scDNAseq, DLP+ library construction was carried out as described(Zahn et al., 2017). In brief, single cell suspensions were fluorescently stained using CellTrace CFSE (Life Technologies) and LIVE/DEAD Fixable Red Dead Cell Stain (ThermoFisher) in a PBS solution containing 0.04% BSA (Miltenyi Biotec 130-091-376) incubated at 37 °C for 20 min. Cells were subsequently centrifuged to remove stain, and resuspended in fresh PBS with 0.04% BSA. This single cell suspension was loaded into a contactless piezoelectric dispenser (sciFLEXARRAYER S3, Scienion) and spotted into the open nanowell arrays (SmartChip, TakaraBio) preprinted with unique dual index sequencing primer pairs. Occupancy and cell state were confirmed by fluorescent imaging and wells were selected for single cell CN profiling. In brief, cell dispensing was followed by enzymatic and heat lysis. After lysis, tagmentation mix (14.335 nl TD Buffer, 3.5 nl TDE1, and 0.165 nl 10% Tween-20) in PCR water was dispensed into each well followed by incubation and neutralization. Final recovery and purification of single cell libraries was done after 8 cycles of PCR. Cleaned up pooled single-cell libraries were analysed using the Aglient Bioanalyzer 2100 HS kit. Libraries were sequenced at UBC Biomedical Research Centre (BRC) in Vancouver, British Columbia on the Illumina NextSeq 550 (mid- or high-output, paired-end 150-bp reads) and at the GSC on Illumina HiSeq2500 (paired-end 125- bp reads) and Illumina HiSeqX (paired-end 150-bp reads).

For scRNAseq, in total 7,000 cells per sample were loaded into a Chromium Chip A (PN-1000009) and processed according to the Chromium Single Cell V(D)J Reagent Kit User Guide. Libraries were constructed using the Chromium Single Cell 5' Library and Gel Bead kit (PN-1000006) and Chromium i7 Multiplex kit (PN-120262). Target enrichment from cDNA was performed by using the Human B Cell Single Cell V(D)J enrichment kit (PN-1000016). 5'Gene expression libraries from 2 samples were pooled and sequenced in a HiSeq2500 instrument (Recipe: 125Cycles Read1, 8 cycles index, 125 cycles Read2). BCR libraries from 18 or 20 libraries were pooled and sequenced in a NextSeq 550 instrument (High Output Flow Cell V2.5 Recipe: 150 cycles Read1, 8 cycles index, 150 cycles Read2).

#### Data processing of scDNA-seq and scRNA-seq

scDNA-seq: FASTQ reads for each patient were obtained from NextSeq 550 instruments and were used as input into our automation pipeline (https://svn.bcgsc.ca/bitbucket/projects/SC/repos/single_cell_pipeline/browse). Reads were trimmed with TrimGalore to remove adapters and paired-end single-cell FASTQs were aligned with bwa, aln mode. PCR duplicates were marked using picard MarkDuplicates and read alignment metrics are computed for each cell. Reads were then tabulated for non-overlapping 500k genomic regions or "bins". A modal regression normalization was performed to reduce GC bias. Copy number was called using HMMcopy, under 6 possible ploidy settings and a fit was computed for each ploidy with the best fit returned per cell.

scRNA-seq: BaseCall files after sequencing were used to generate library-specific FASTQ files with cellranger mkfastq (v3.0.2) prior to running cellranger count (v3.0.2) with the GRCh38 reference to produce cell barcode-gene expression matrices using default settings. Cell cycle scores were computed with cyclone from the scran R package (Version 1.16.0). For single-cell VDJ datasets, cellranger vdj (v3.1.0) was run using the refdata-cellranger-vdj_GRCh38_alts_ensembl-3.1.0-3.1.0 reference from 10x Genomics using default settings. Poor quality contigs that either did not map to immunoglobulin chains or were designated incomplete by cellranger were discarded.

After sequencing and processing, gene expression data were analyzed with monocle3 and visualized on UMAP; DLP data were analyzed with a DLP platform (Zahn et al., 2017). Gene expression and genomic data were then integrated thanks to clonealign (Campbell et al., 2019). TME and tumor B cell interaction analysis was performed based on 10X data with CellChat(Jin et al., 2021) (see methods above).

#### Dimension reduction and clustering analysis for scRNA-seq

The resulting count expression matrices were read into SingleCellExperiment objects. Outlier cells according to quality control parameters (>= 3 median absolute deviations from the median) were filtered out using the scater R package (Version 1.16.2). In addition, cells with >= 10% mitochondrial UMIs and >= 60% ribosomal UMIs were removed. Size factors were computed using quickCluster and computeSumFactors from the scran R package (1.16.0). Data normalization was then performed using scater normalize function. Gene-specific variance was calculated using scran trendVar and then was decomposed into biological and technical components. Normalized logcounts for the genes with biological variance >= 0 were used as input into the scanorama (Hie et al., 2019) R package (Version 1.6) to perform batch correction. Principal components analysis was performed on the resulting batch-corrected expression matrix for the top 1000 most variable genes. The first 50 PCs were used as input for UMAP. Entropy of cell expression before and after batch correction was assessed using the method as previously described (cite: single-cell map of diverse immune phenotypes in the breast tumor microenvironment). Briefly, a nearest-neighbor graph was generated using Euclidean distance with either the first 50 PCs from the non-batch-corrected data, or the first 50 PCs from the batch-corrected data. For each cell, the proportion of neighborhood cells from each other samples was calculated, and these were used to compute the Shannon entropy of the cell. The entropy distributions were compared to assess the performance of batch correction (Supplementary Figure 3).

Initial and subsequent unsupervised clustering was performed with monocle3 (Version 0.2.3.0), using the default leiden algorithm and automatic resolution. Cluster-specific gene expressions were extracted using the monocle3 top_markers function. Clusters from PhenoGraph were manually assigned to a cell type by comparing the mean expression of known markers across cells in a cluster. Markers used for cell type annotation are described in the Supplementary Table 8 (Aoki et al., 2020).

The Shannon entropy was computed at cluster level to assess the sample heterogeneity. It is also used to quantify the heterogeneity of tumor subpopulations between the two timepoints after the individual reclustering of malignant B cell populations. For each cell in the UMAP (after reclustering), we find its 20 nearest neighbors, calculate the proportion of tFL-FL cells and tFL-DLBCL cells within these 20 neighbors, and then calculate the entropy for this given cell. Then we take the average of the entropy values and use this averaged value to represent a patient, with greater values representing greater mixing between the two timepoints.

#### Phylogenetic analysis for scDNA-seq

The copy number information output from HMMcopy was used to construct the phylogenetic trees with a single cell Bayesian tree reconstruction method called sitka (Salehi et al.). In brief, the sitka transforms the copy number data into binary breakpoint profile and then performs Markov chain monte carlo algorithms with tree exploration moves to generate the optimal phylogenetic topology. The inferred trees were post-processed to identify clonal populations from major clades. Clones were constructed by identifying connected components (each a clade or a paraphyly) in the phylogenetic tree reconstruction and each clone is composed of cells of sufficient genomic homogeneity. For each tFL-FL cell in a tree, we calculated the average number of nodes traversed to reach each tFL-DLBCL cell, defined as the DLP tree distance.

#### Association of scDNAseq and scRNAseq with clonealign

To prepare the clonealign (Campbell et al., 2019) input from scDNA-seq data, the overlap between gene position and region-based copy number profile was identified and then the matrix of the gene by clone with copy number as entries was generated. Genes were first filtered by only autosomal genes and by genes that were uniquely mapped. Genes with multiple copy numbers that differ by more than 1 within the same clone were removed from the downstream analysis. Pairwise Manhattan distance for clones was calculated and clones with Manhattan distance less than 0.005 were merged as one clone. The clonealign input from scRNA-seq data was a gene expression matrix with row names as the genes and column names as the cells.

These two matrices are then used as input to clonealign, which is a statistical method to assign cells measured with single-cell RNA-seq to clones derived from low-coverage single cell

DNAs-seq based on the probability that the observed gene expression levels are consistent with the copy number profiles of the clones(Campbell et al., 2019). It is based on an assumed positive correlation between copy number and gene expression.

#### Differential expression and enrichment analysis

Differential expression was performed using the limma R package (Version 3.44.3) with covariate terms including timepoint, patient, and batch information. The resulting log fold change values (filtering out ribosomal and mitochondrial genes) were then used as input for gene set enrichment analysis with the pre-ranked mode in GSEA software (v4.0.3), using default parameters with 1000 permutations, and hallmark pathway gene set (The Molecular Signatures Database Hallmark Gene Set Collection). For enrichment of pathways, genes with a significant p value were ranked by log fold change (logFC) in differential expression between 2 timepoints. All reported Q values refer to Benjamini-Hochberg corrected P values for two-sided tests.

#### Cell-cell interaction analysis

Cell-cell interaction analysis was performed using the CellChat R package (Version 1.1.2) for each timepoint. The CellChat ligand-receptor database was modified to include the MHC II - LAG3 ligand-receptor pairs and the signaling pathway categories used were "Secreted Signaling" and "Cell-cell Contact". Signaling pathways with a significant p value were selected and then subjected to manual reviews. The final signaling pathways included in the study were: IL10 - (IL10RA+IL10RB), CXCL13 - CXCR5, MIF - (CD74+CXCR4), IL16 - CD4, IL7 - (IL7R+IL2RG), GZMA - PARD3, FASL - FAS, MIF - (CD74+CD44), IL10 - (IL10RA+IL10RB), IL15 - (IL15RA+IL2RB), CXCL12 - CXCR4, CD70 - CD27, BTLA - TNFRSF14, TIGIT - NECTIN2, HLA-A - CD8B, HLA-B - CD8B, HLA-C - CD8B, HLA-A - LAG3, HLA-B - LAG3, HLA-C - LAG3, HLA-DRA - LAG3, HLA-DRB1 - LAG3, HLA-DRA - CD4, HLA-DRB1 - CD4, and HLA-E - KLRC1.

#### TMA, FISH and IHC analysis

FISH analysis was performed on TMA using commercially available dual-color break-apart probes *MYC*, *BCL2* and *BCL6* (Metasystems). Images were captured using a Metafer CoolCube

1 camera and Metafer software (version 3.11.8). Scoring was performed on 100 cells by a cytogenetic expert for each case. Tumors displaying break-apart signals in ≥5% of cells were considered to harbor a translocation. Gain was defined as 3-4 fused signals and amplification was defined as 5+ signals.

Three different panels were employed in the multicolor IHC: CD20-CD10-CD4-CD8-FOXP3-LAG3-PD1 (REG panel, Supplementary Table 10), CD20-CD70-CD4-CD8-FOXP3-LAG3-CD27 (CD27-CD70 panel, Supplementary Table 11) and CD20-CXCL13-CXCR5-BCL6-CD4-CD10-PD1 (TFH panel, Supplementary Table 12). TMA slides were deparaffinized in xylene, fixed for an additional 20 min in neutral buffered formalin and rinsed with dH2O. For the REG panel, antigen retrieval was first performed in a Decloaking Chamber Plus with Diva Decloaker (all products from Biocare Medical unless otherwise indicated). After appropriate blocking, the primary antibody for CD20 (mouse, clone L26, Cat. # CM004) was diluted 1/300 in DaVinci Green (DVG), applied and incubated for 30 min in an Intellipath FLX rack at room temperature, followed by detection using Mach2 mouse HRP. Visualization of CD20 was achieved using Opal Polaris 480 at 1/100 dilution (Akoya Biosciences). This was the first of 7 rounds using the Intellipath FLX automated IHC staining system. For the second round, the TMA slide was placed into AR6 buffer (PerkinElmer) and heated to boiling in a microwave oven, then incubated with the primary antibody for FOXP3 (mouse, clone 236A/E7, Abcam, Cat. # Ab20034) at 1/100 dilution in DVG, followed by detection using Mach2 mouse HRP and visualization using Opal 690 at 1/150 dilution. For the third round, the TMA slide was once again placed into AR6 buffer and heated using a microwave, then incubated with the primary antibody for CD8 (mouse, clone C8/144B, Cell Marque, Cat. # 108M-94) at 1/800 dilution in DVG, followed by detection using Mach2 mouse HRP and visualization using Opal 520 at 1/300 dilution. For the fourth round, the TMA slide was once again placed into AR6 buffer and heated using a microwave, then incubated with the primary antibody for LAG3 (rabbit, clone D2G40, Cell Signaling Technology, Cat. # 15372T) at 1/300 dilution in Renoir Red (RR), followed by detection using Mach2 rabbit HRP and visualization using Opal 540 at 1/400 dilution. For the fifth round, the TMA slide was once again placed into AR6 buffer and heated using a microwave, then incubated with the the primary antibody for CD4 (rabbit, clone EPR6855, Abcam, Cat. # Ab133616) at 1/100 dilution in DVG, followed by detection using Mach2 rabbit HRP and visualization using Opal 570 at 1/200 dilution. For the sixth round, the TMA slide was once again placed into AR6 buffer and heated using a microwave, then incubated with the primary antibody for PD1 (mouse, clone NAT105, Cell Marque, Cat. # 315M-94) at 1/200 dilution in RR, followed by detection using Mach2 mouse HRP and visualization using Opal 650 at 1/200 dilution. For the seventh and final round, the TMA slide was once again placed into AR6 buffer and heated using a microwave, then incubated with the primary antibody for CD10 (rabbit, clone E5P7S, Cell Signaling Technology, Cat. # 65534S) at 1/150 dilution in DVG, followed by detection using Mach4 rabbit HRP (10-minute polymer) and visualization using Opal 620 at 1/100 dilution. Details of the REG panel workup described above are also provided in tabular format below, along with those for the CD27-CD70 panel and TFH panel. Nuclei were visualized with DAPI staining and sections were coverslipped using Fluoro Care Anti-Fade Mountant. TMA slides were scanned using the Vectra multispectral imaging system (PerkinElmer) following the manufacturer's instructions to generate .im3 image cubes for downstream analysis. Optimal exposure times for fluorophores ranged between 50 and 200ms.

InForm image analysis software (v2.4.4; PerkinElmer) was used to analyze the spectra for all fluorophores.Cells were phenotyped as positive or negative for each of the 7 markers within a given panel. Data were merged in R by X-Y coordinates so that each cell could be assessed for all markers simultaneously. Distance and neighbor analysis were performed with the spatstat R package (v1.58-2). Specifically, we calculated the average distance between every malignant B cell to its nearest neighbor with a specific phenotype across each image and also calculated the average proportion of cells with a given phenotype within a 125-micron radius surrounding every malignant B cell.

#### Immunohistochemistry in validation cohorts

Briefly, AR6 buffer (PerkinElmer, USA) with Diva decloaker (Biocare Medical, USA) was used for microwave heat antigen retrieval in each primary antibody reaction. Secondary antibodies (Opal 620 for CD20, Opal 690 for FOXP3, Opal 520 for CD8, Opal 540 for LAG-3, Opal 570 for CD4, and Opal 650 for PD-1) were used for visualization. TMA slides were scanned using the Vectra multispectral imaging system (PerkinElmer, USA) according to manufacturer's instructions and InForm image analysis software (PerkinElmer, USA) was used to analyze all fluorophores. Cells were phenotyped as positive or negative for each of the six markers (PD-1, LAG-3, CD4, CD8, FOXP3, CD20).

#### Survival analysis

For prediction, the proportion of LAG3+/CD8+/CD20- cells out of CD20 negative cells was calculated for the R (cells with both CD4 and CD8 expressions were removed from the analysis; the mean value from the 2 cores of a patient was taken for analysis). Survival (disease specific survival, time to progression and time to transformation) and probability of transformation analysis were performed with Kaplan-Meier estimates and log-rank tests using the survminer (0.4.9) and survival (3.3-1) R packages. Multivariate analysis was performed using FLIPI score, percentage of LAG3CD8 of cells out of TME cells and FL grading as covariate. The cutoffs for the percentage of CD8+LAG3+ cells out of CD20- cells were selected as the top 25% quantile for the RB and RCVP cohort. Clinical data of these 2 cohorts are presented in Supplementary Table 9 and 10.

### Supplementary figures

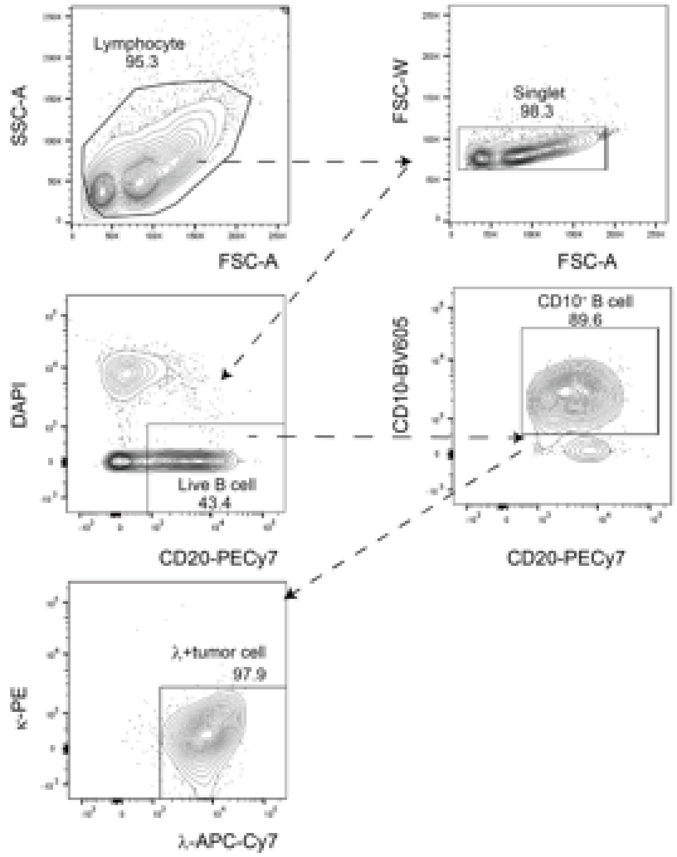


**Supplementary Figure 1.** Flow gating strategy. Viable cells (DAPI-negative) were sorted on a FACS ARIAIII or FACS Fusion (BD Biosciences) using an 85-μm nozzle. For DLP sequencing, malignant B cells were sorted based on CD20 and CD10 positive expression (as previously assessed at time of diagnosis) and based on kappa or lambda light chain expression.

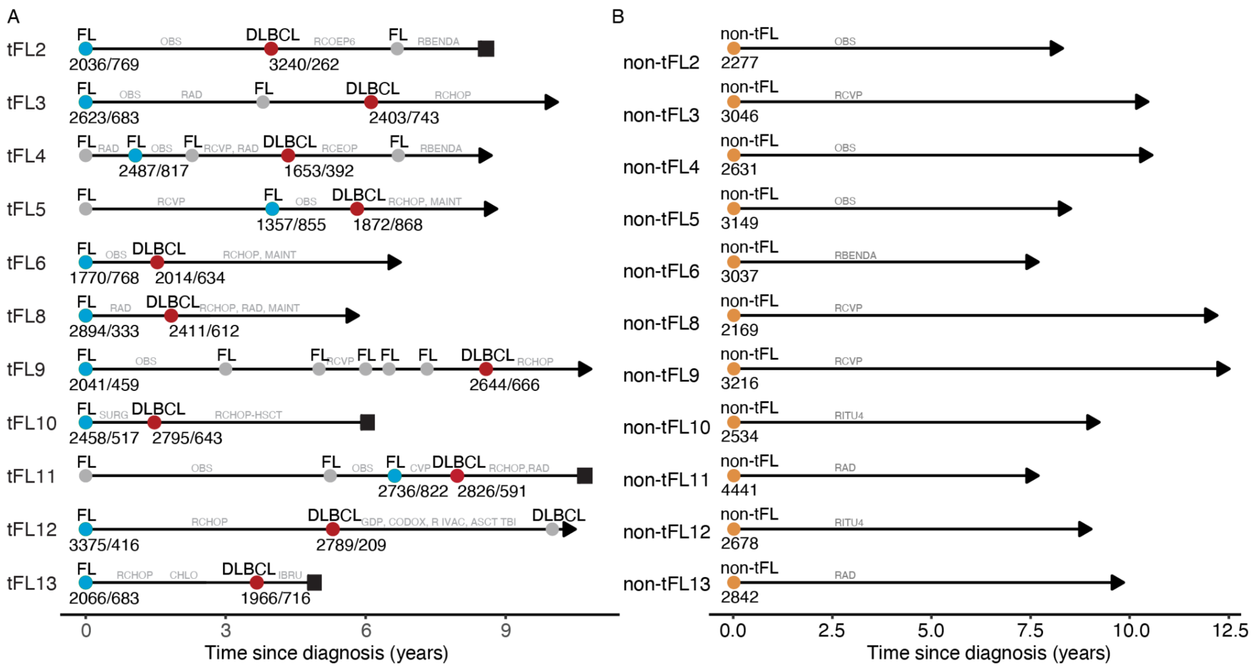


**Supplementary Figure 2. Timelines for the patient cohort.**

**Panel A:** Timelines are represented for pairs, with time in x axis. Blue dots represent the tFL-FL timepoint analyzed and red dots the tFL-DLBCL timepoint. Numbers below the dots represent the number of cells recovered in 10xRNA and in DLP respectively. The grey dots represent relapse without biopsy available. Scares represent dead patients and arrow patients still alive and in follow-up. Treatment information are provided in grey : OBS: observation, RCEOP: rituximab, Cyclophosphamide, Etoposide, Prednisolone, Vincristine; RCVP: Rituximab, Cyclophosphamide, Vincristine, Prednisolone; RCHOP: Rituximab, Cyclophosphamide, Adriamycine, Oncovin, Prednisolone. RAD: radiation; BENDA: Bendamustine; HSCT: hematopoietic stem cell transplant

**Panel B:** Timelines and treatment information for non-transformed FL samples. None of these patients relapsed.

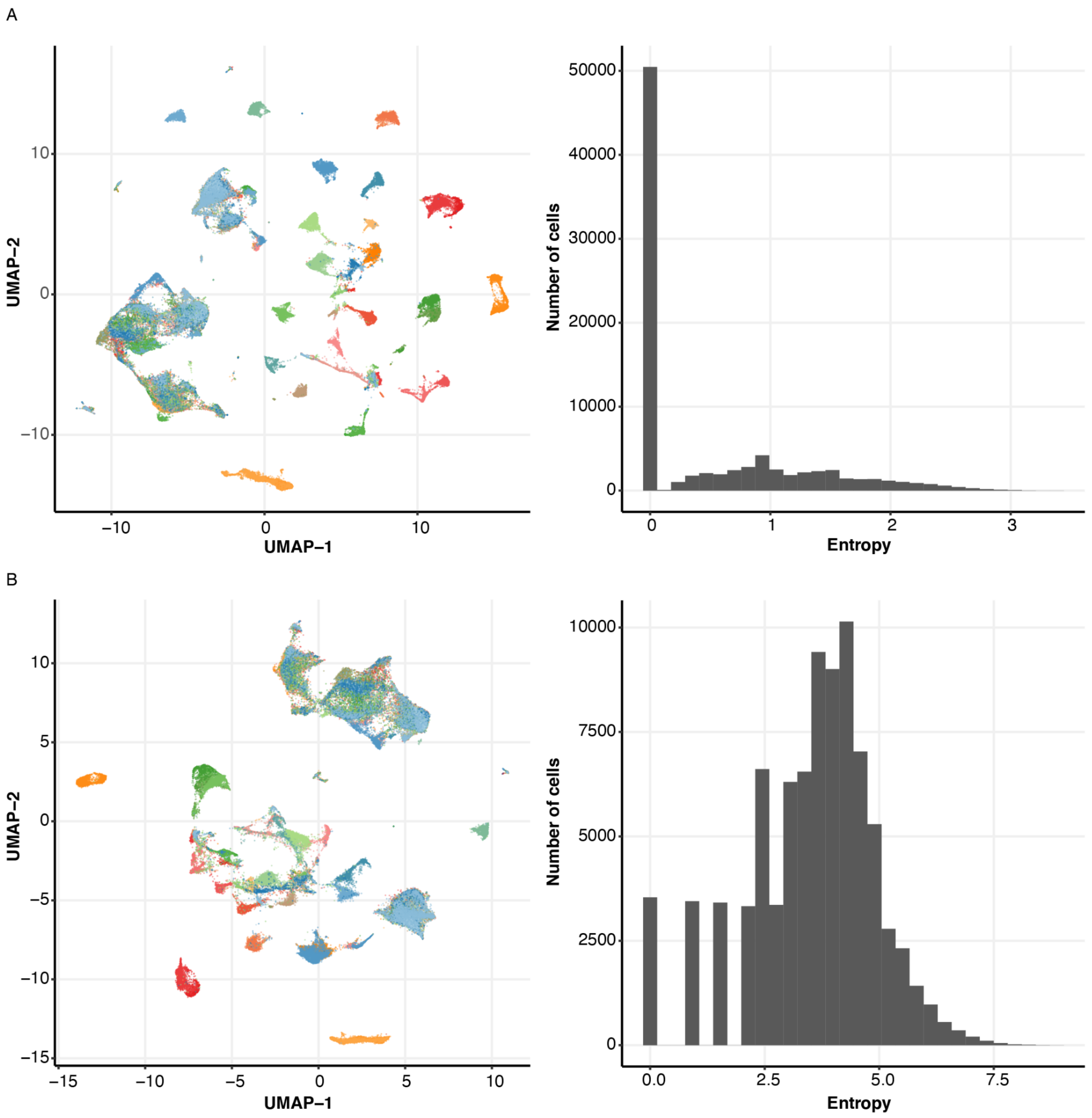


**Supplementary Figure 3.** Batch correction. In this representation, the expression data from all cells of the 11 tFL-pairs, 11 non-tFL and 2 RLN are merged (35 samples). Batch correction and normalization was performed.

**Panel A**: UMAP visualization and cell entropy before batch correction

**Panel B**: UMAP visualization and cell entropy after batch correction. Removal of batch effects (caused by single-cell isolation and library preparation in different experimental runs) resulted in improved mixing of cells across samples, as demonstrated by a significant increase in cell entropy (Wilcoxon–Mann–Whitney $P < 0.001$).

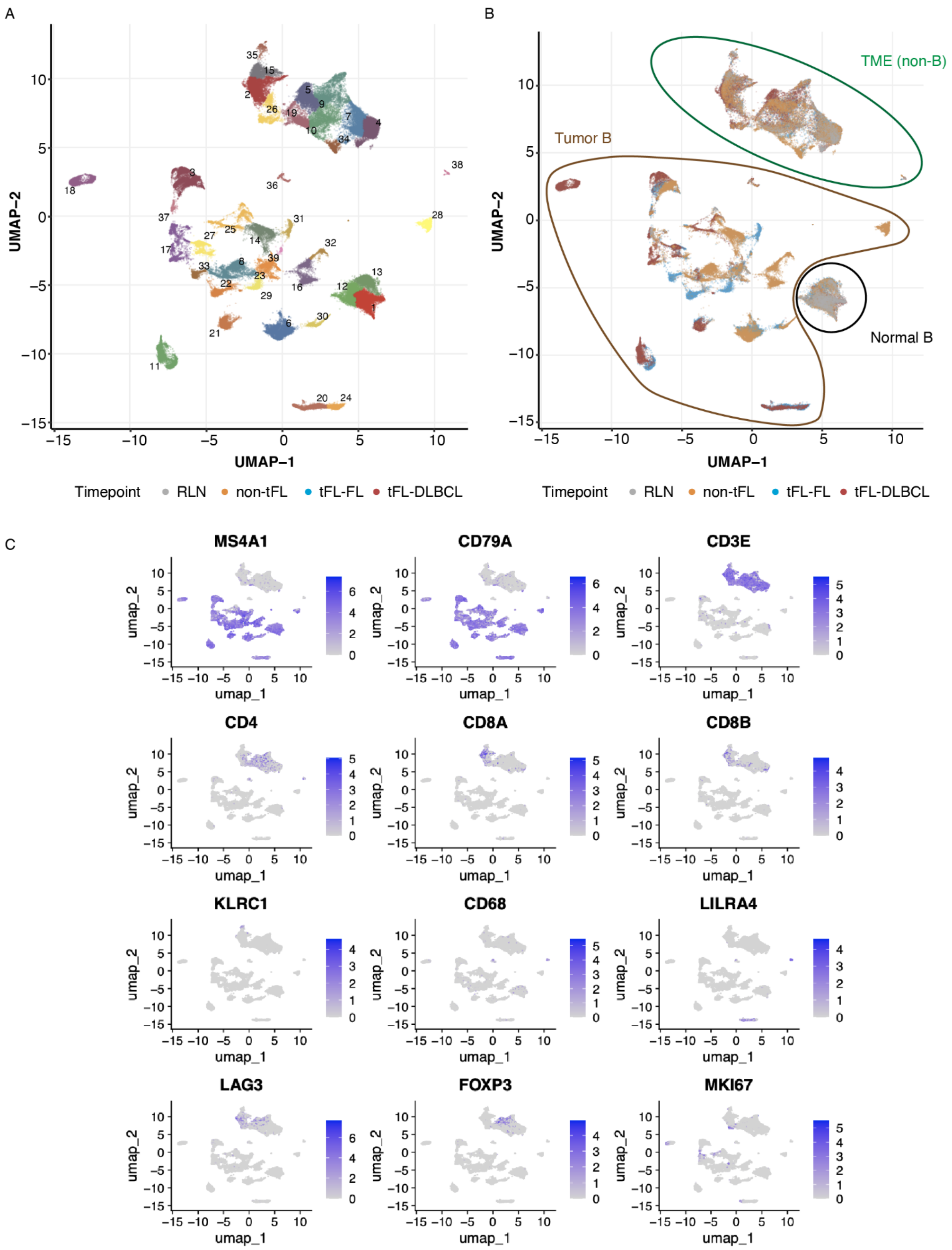


**Supplementary Figure 4: scRNA data clustering and cell type annotation.**

**Panel A:** Unsupervised clustering using Monocle3 (version 0.2.2) followed by visualization in UMAP space identified 39 expression-based cell clusters (panel A).

**Panel B:** cells are colored based on their cohort: tFL-FL in blue (follicular lymphoma timepoint within the tFL pairs)), tFL-DLBCL in red (diffuse large B cell lymphoma timepoint within the tFL pairs), non-tFL in orange (non-transformed follicular lymphoma) or reactive lymph nodes in grey. RLN B cells are clustered with few suspected normal B cells from tumor samples. TME clusters included cells from all the cohorts. Tumor B cells clusters were mainly driven by patients' heterogeneity.

**Panel C**: UMPA is colored based on the level of expression of selected genes, to identify the B cells (normal and malignant) and non-B cells (or tissue microenvironment cells). Key markers used for broad classification are as follows: MS4A1/CD79A/CD3E/CD4/CD8A/CD8B/KLRC1/CD68/LIL4R/LAG3/FOXP3/MKI67.

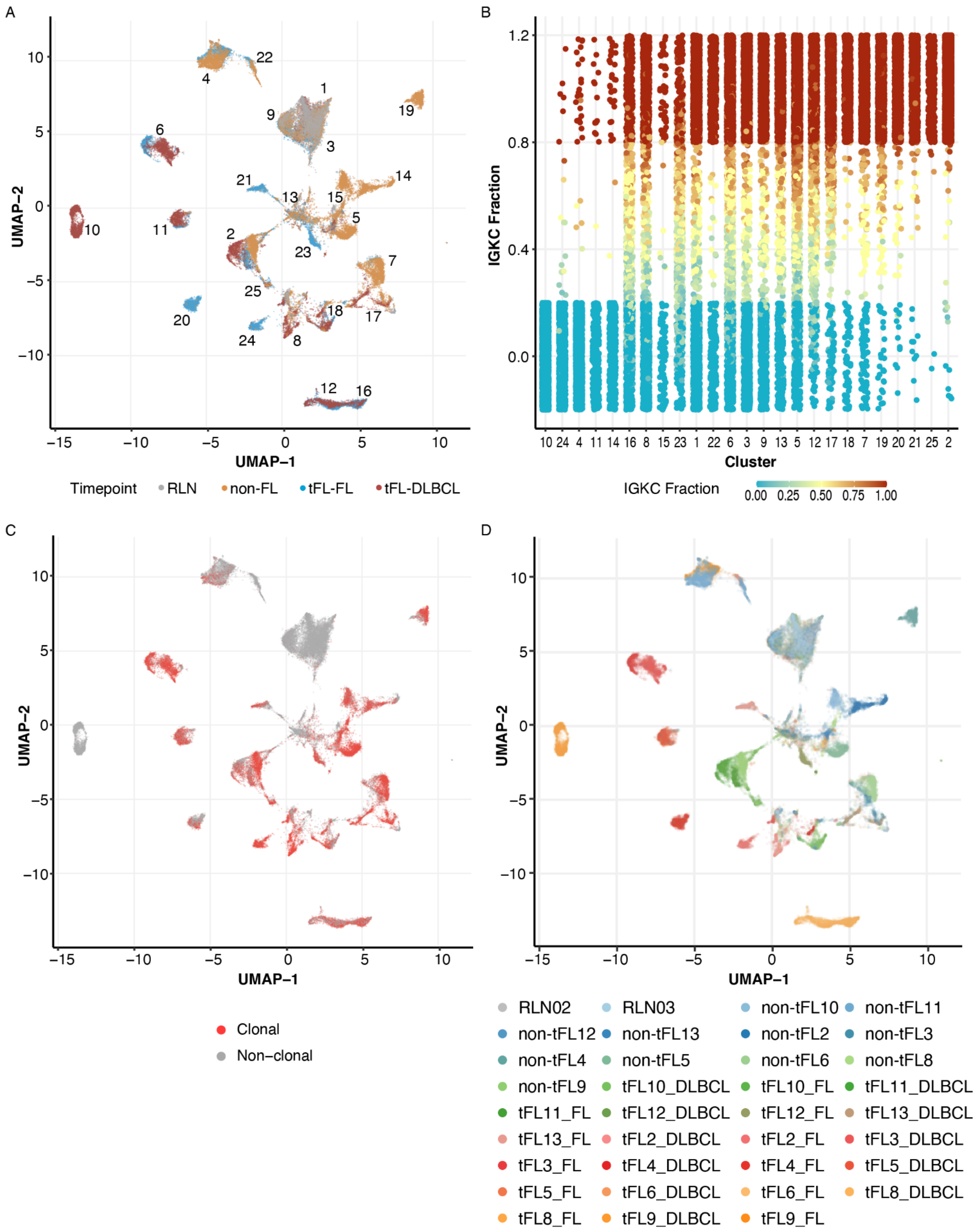


**Supplementary Figure 5. Identification of malignant and non-malignant B cells.**

**Panel A**: UMAP representation and clustering of B cells only, cells are colored based on their cohort: tFL-FL in blue (follicular lymphoma timepoint within the tFL pairs)), tFL-DLBCL in red (diffuse large B cell lymphoma timepoint within the tFL pairs), non-tFL in orange (non-transformed follicular lymphoma) or reactive lymph nodes in grey.

**Panel B**: Kappa/lambda scattered plot for each cluster identified in the UMAP. The color scale represents the IGKC fraction (y axis) defined as IGKC positive B cells out of (IGKC + IGLC) positive B cells (kappa/lambda double positive/negative cells were removed from the analysis).

Normal B cells clusters were identified as being cluster 1-3-9: including RLN B cells and polytypic B cells from tumor samples; as well as cluster 22: including RLN cells and polytypic B cells from tumor samples (nontFL-11, tFL-FL-9 and 13). None of the cells included in these clusters 1-3-9-22 had a monoclonal V-D-J rearrangement as identified in BCR sequencing.

Of note, cells in Cluster 16 included only tFL-FL-8 sample; Cluster 8 is a mix of tumor samples (majority of tFL-DLBCL-2, lambda monoclonal; the other samples are also monotypic in cluster 8); Cluster 15 contains cells from non-tFL-10 (double negative for kappa/lambda expression in 10X and Lambda clonal in BCR sequencing); Cluster 23 contains cells from tFL-FL-12 (double expressor for kappa and lambda in 10X); Cluster 6 cells from Pair 3 only (double expressor for kappa and lambda in 10X, but kappa clonal in BCR sequencing as well as in kappa in Flow Cytometry); Cluster 13 contains cells from mixed tumor samples, with almost no RLN cells (tumor sample cells are monotypic, except for the dominant sample in this cluster: tFL-FL-12 double expressor kappa lambda); and cluster 5 a mix of tumor samples from non-tFL-5 (expressing kappa in 10X) plus tFL-FL-6 (double expressor for kappa and lambda in 10X) and non-tFL-FL-13 (expressing kappa in 10X).

**Panel C:** UMAP colored by clonal V-D-J in BCR sequencing. tFL-pair-9 (cluster 10) is the only case where no clonal VDJ could be identified, likely due to the abundance of SHM and absence of productive VDJ. When looking at the kappa/lambda scatter plot, we can see that this cluster 10 contains only kappa cells. Cluster 4, containing only cells from non-tFL-11, has also only few VDJ clonal cells. But we can see that cells in this cluster 4 are all expressing the kappa chain, confirming the tumoral origine. Cluster 20, containing a vast majority of cells from tFL-FL-4, also presents only a few VDJ clonal cells. But we can see that cells in this cluster 20 are all expressing the lambda chain, confirming the tumoral origine. Finally, cluster 13 is a mix of cells from different tumor samples, with almost no RNL cells (cf panel B).

**Panel D**: UMAP colored by patients and samples.

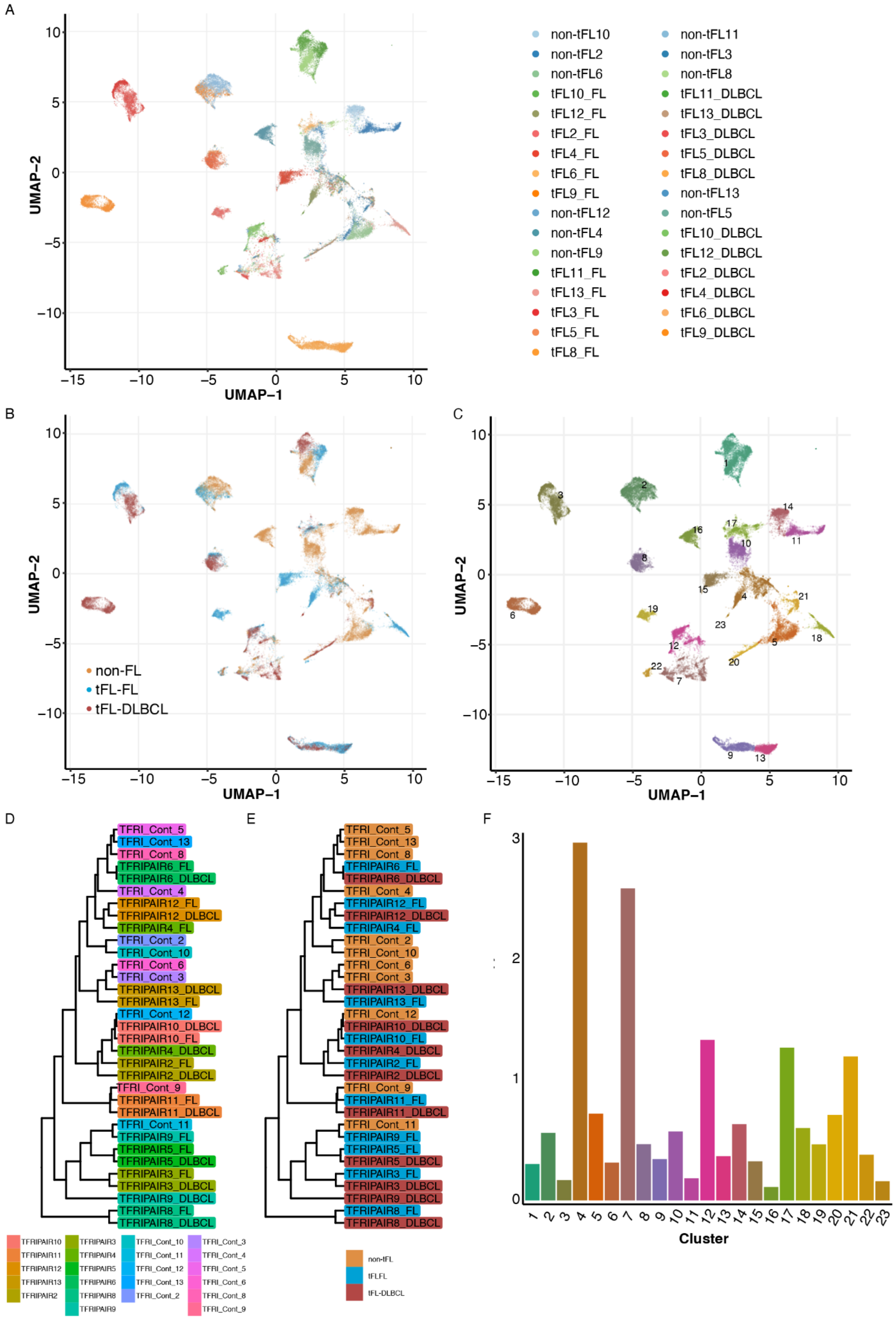

A
UMAP−1
UMAP−2
non-tFL10
non-tFL11
non-tFL2
non-tFL3
non-tFL6
non-tFL8
tFL10_FL
tFL11_DLBCL
tFL12_FL
tFL13_DLBCL
tFL2_FL
tFL3_DLBCL
tFL4_FL
tFL5_DLBCL
tFL6_FL
tFL8_DLBCL
tFL9_FL
non-tFL13
non-tFL12
non-tFL5
non-tFL4
tFL10_DLBCL
non-tFL9
tFL12_DLBCL
tFL11_FL
tFL2_DLBCL
tFL13_FL
tFL4_DLBCL
tFL3_FL
tFL6_DLBCL
tFL5_FL
tFL9_DLBCL
tFL8_FL
B
non-FL
tFL-FL
tFL-DLBCL
C
D
E
F
Cluster
non-tFL
tFLFL
tFL-DLBCL

**Supplementary Figure 6.** Malignant B cells only reclustering (with tFL-FL, tFL-DLBCL and non-tFL samples).

**Panel A**: UMAP visualization of malignant B cells, colored by sample name. To further analyze malignant B cell phenotypic evolution, the malignant B cells were then re-clustered independently leading to 23 independent clusters with a strong patient but no timepoint related clustering effect. A UMAP tumor sample distance analysis showed that tFL- FL and tFL-DLBCL samples in each PAIR were closer together than with other samples

**Panel B**: UMAP visualization of malignant B cells, colored by timepoints.

**Panel C**: UMAP visualization of malignant B cells, colored by clusters.

**Panel D**: Sample distance trees: colored by sample name.

**Panel E**: Sample distance trees: colored by timepoint.

**Panel F**: Entropy in malignant B cells.

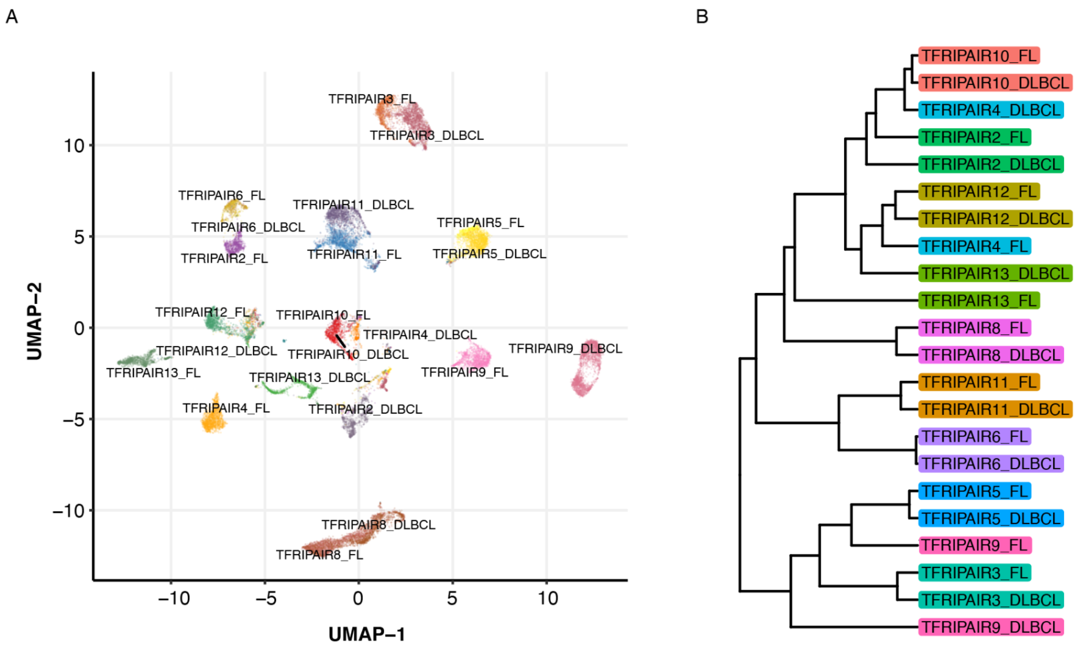


**Supplementary Figure 7:** Reclustering malignant B cells from pairs only (tFL-FL and tFL-DLBCL).

**Panel A:** UMAP visualization of malignant B cells, colored by sample name. The UMAP highlights the patients' related clustering effect.

**Panel B:** Sample distance tree showing the high inter-patient heterogeneity, and no timepoint related clustering.

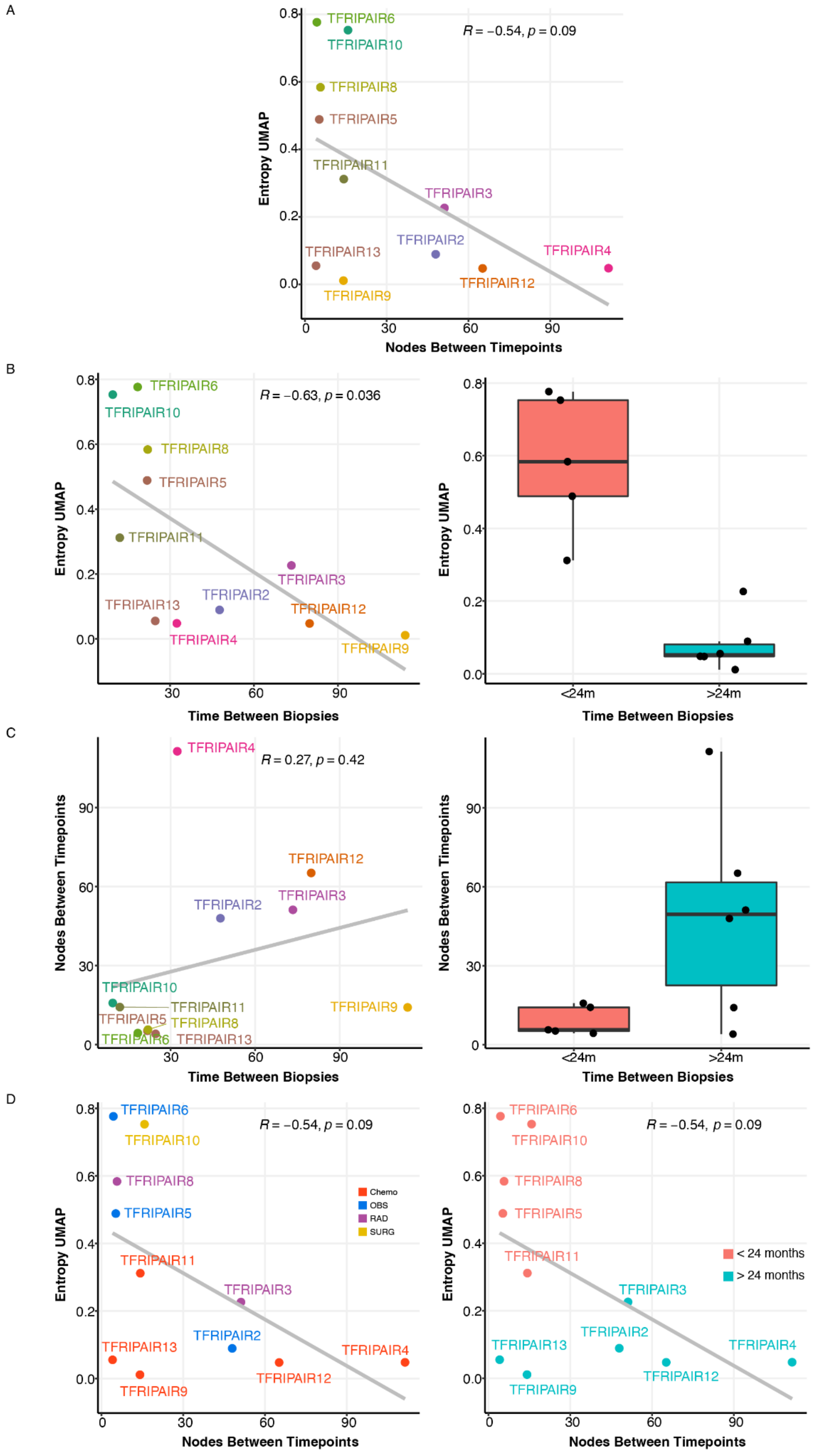
A
B
C
D
R = −0.54, p = 0.09
R = −0.63, p = 0.036
R = 0.27, p = 0.42
Entropy UMAP
Nodes Between Timepoints
Time Between Biopsies
TFRIPAIR2
TFRIPAIR3
TFRIPAIR4
TFRIPAIR5
TFRIPAIR6
TFRIPAIR8
TFRIPAIR9
TFRIPAIR10
TFRIPAIR11
TFRIPAIR12
TFRIPAIR13
<24m
>24m
Chemo
OBS
RAD
SURG
< 24 months
> 24 months

**Supplementary Figure 8.** Genomic and phenotypic evolution within each pair, using UMAP entropy and corrupt tree distance correlations. To compare and measure the clonal and phenotypic co-evolution, we defined the DLP tree distance in DLP, and the entropy UMAP in 10X data. Within each pair re-clustering, the entropy of timepoints is calculated for each cell with the proportions of tFL-FL cells and tFL-DLBCL cells within the 20 nearest neighbors of this given cell. The larger entropy values represent greater timepoint similarity, and value closer to 0 representing clear distinction between tFL-FL and tFL-DLBCL timepoint in a pair. The DLP tree distance is defined as the average number of nodes between tFL-FL and tFL-DLBCL cells in DLP.

**Panel A:** correlation between entropy UMAP (y axis) and number of nodes (x axis), for each pair. For the majority of the pairs, we observe a significant inverse correlation (R=-0.55, p=0.09): pairs with a high similarity in 10X between the 2 timepoints present a low number of nodes; except for PAIR 13 and 9 that have a very low entropy but also very few nodes between the 2 timepoints. These data suggest that the CNV analysis does not capture all the genomic modifications that will lead to phenotypic evolution in the transformation process.

Pairs with the lowest phenotypic and genomic evolution burden were those were transformation was capture early one (ie less than 2 years between the 2 biopsies: PAIR 11-10-5-6 and 8) whereas those with the highest evolution burden were those where the transformation was captured after 2 years (ie more than 2 years between tFL-FL and tFL-DLBCL biopsy, PAIR 4-3-12-2).

**Panel B:** Entropy UMAP (y axis) versus time (x axis) elapsed between the 2 biopsies for each pair. We observe a significant inverse correlation. A boxplot with a binary split for time value is also provided (less than 2 years between biopsies or more than 2 years).

**Panel C:** Number of nodes (or distance) versus time between the 2 biopsies for each PAIR. Pairs with less than 24 months between tFL-FL and tFL-DLBCL timepoint present a very close genomic profil. The majority of the pairs with more than 24 months between the 2 biopsies had a complexe evolution with a higher number of nodes between the 2 timepoints, except for PAIRS 13 and 9.

**Panel D:** Correlation between Entropy UMAP (y axis) and number of nodes (or distance, x axis), by pair, and colored by treatment type received between the 2 biopsies. Chemo = chemotherapy, OBS= observation (watch and wait), RAD= radiation. Treatment received does not seem to create a greater bias than the time interval between the 2 biopsies. In both PAIR2 and 6 a watch and wait strategy was applied, but one had a drastic phenotypic and genomic shift, and the other a unique mixed clone. As well, PAIR12 and 13 received the same R-CHOP treatment before transformation, but had different patterns of genomic evolution, and while radiation was applied in both PAIR3 and 8, these two pairs presented a very distinct UMAP entropy and nodal distance score. The other way around, 3 different treatment options were applied in PAIR2, 3 and 4 (WW, radiation and chemotherapy respectively), but these 3 pairs presented the same pattern of phenotypic and genomic evolution.

A

| Pathway | Gene ranks | NES | pval | padj |
|---|---|---|---|---|
| HALLMARK_MYC_TARGETS_V1 | | 7.90 | 1.0e−50 | 2.4e−49 |
| HALLMARK_OXIDATIVE_PHOSPHORYLATION | | 7.10 | 1.0e−50 | 2.4e−49 |
| HALLMARK_E2F_TARGETS | | 4.70 | 2.2e−37 | 3.5e−36 |
| HALLMARK_G2M_CHECKPOINT | | 3.99 | 3.0e−26 | 3.6e−25 |
| HALLMARK_MYC_TARGETS_V2 | | 3.59 | 2.0e−10 | 1.1e−09 |
| HALLMARK_MTORC1_SIGNALING | | 3.58 | 1.8e−22 | 1.7e−21 |
| HALLMARK_DNA_REPAIR | | 3.08 | 4.9e−13 | 3.4e−12 |
| HALLMARK_UNFOLDED_PROTEIN_RESPONSE | | 2.84 | 9.7e−10 | 4.7e−09 |
| HALLMARK_GLYCOLYSIS | | 2.67 | 1.2e−10 | 7.4e−10 |
| HALLMARK_REACTIVE_OXYGEN_SPECIES_PATHWAY | | 2.62 | 4.4e−06 | 1.3e−05 |
| HALLMARK_FATTY_ACID_METABOLISM | | 2.45 | 3.8e−08 | 1.4e−07 |
| HALLMARK_PROTEIN_SECRETION | | 2.24 | 2.6e−05 | 6.2e−05 |
| HALLMARK_ADIPOGENESIS | | 2.19 | 1.1e−06 | 3.4e−06 |
| HALLMARK_PI3K_AKT_MTOR_SIGNALING | | 1.82 | 7.6e−04 | 1.5e−03 |
| HALLMARK_PEROXISOME | | 1.66 | 4.2e−03 | 7.2e−03 |
| HALLMARK_MITOTIC_SPINDLE | | 1.65 | 2.9e−04 | 6.1e−04 |
| HALLMARK_INTERFERON_ALPHA_RESPONSE | | 1.19 | 1.9e−01 | 2.6e−01 |
| | 0 3000 6000 9000 12000 | | | |

B

| Pathway | Gene ranks | NES | pval | padj |
|---|---|---|---|---|
| HALLMARK_TNFA_SIGNALING_VIA_NFKB | | −2.64 | 1.5e−13 | 1.2e−12 |
| HALLMARK_EPITHELIAL_MESENCHYMAL_TRANSITION | | −2.50 | 7.8e−09 | 3.1e−08 |
| HALLMARK_INFLAMMATORY_RESPONSE | | −2.39 | 2.6e−09 | 1.1e−08 |
| HALLMARK_KRAS_SIGNALING_DN | | −2.37 | 3.2e−07 | 1.1e−06 |
| HALLMARK_COAGULATION | | −2.22 | 4.8e−06 | 1.3e−05 |
| HALLMARK_KRAS_SIGNALING_UP | | −2.10 | 4.9e−06 | 1.3e−05 |
| HALLMARK_MYOGENESIS | | −2.06 | 1.2e−05 | 3.0e−05 |
| HALLMARK_IL6_JAK_STAT3_SIGNALING | | −2.06 | 1.0e−04 | 2.2e−04 |
| HALLMARK_APICAL_SURFACE | | −1.93 | 1.9e−03 | 3.5e−03 |
| HALLMARK_ALLOGRAFT_REJECTION | | −1.88 | 4.3e−05 | 9.9e−05 |
| HALLMARK_ESTROGEN_RESPONSE_EARLY | | −1.73 | 5.8e−04 | 1.2e−03 |
| HALLMARK_IL2_STAT5_SIGNALING | | −1.62 | 2.5e−03 | 4.5e−03 |
| HALLMARK_TGF_BETA_SIGNALING | | −1.46 | 6.0e−02 | 9.2e−02 |
| HALLMARK_HEDGEHOG_SIGNALING | | −1.46 | 8.0e−02 | 1.2e−01 |
| HALLMARK_HYPOXIA | | −1.43 | 3.6e−02 | 5.9e−02 |
| HALLMARK_UV_RESPONSE_DN | | −1.41 | 5.6e−02 | 9.0e−02 |
| HALLMARK_WNT_BETA_CATENIN_SIGNALING | | −1.26 | 1.9e−01 | 2.6e−01 |
| HALLMARK_COMPLEMENT | | −1.23 | 1.5e−01 | 2.2e−01 |
| HALLMARK_P53_PATHWAY | | −1.19 | 2.0e−01 | 2.6e−01 |
| HALLMARK_ESTROGEN_RESPONSE_LATE | | −1.16 | 2.5e−01 | 3.2e−01 |
| | 0 3000 6000 9000 12000 | | | |

C

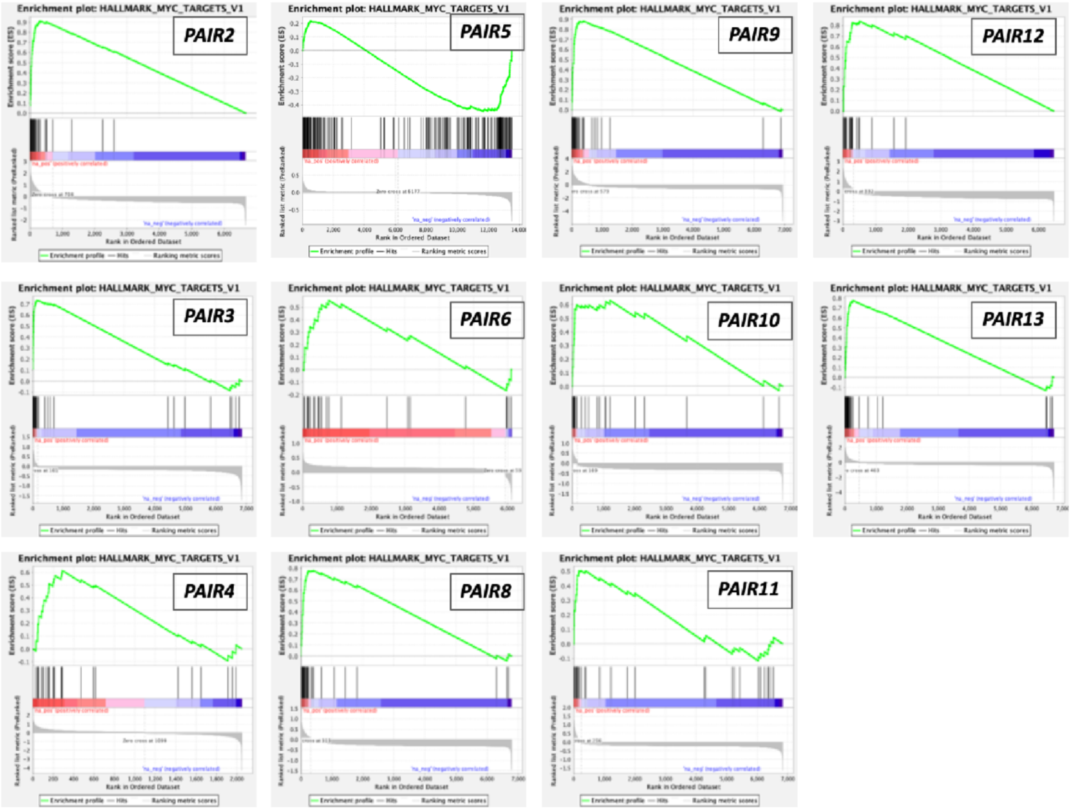

**Supplementary Figure 9.** Gene set enrichment analysis based on a differential expression analysis in tFL-FL cells versus tFL-DLBCL cells. 13303 genes are differentially expressed between tFL-FL and tFL-DLBCL tumor B cells.

**Panel A**: Enrichment plot for the top 20 most significant pathways in the Hallmark data-base positively enriched in the tFL-DLBCL versus tFL-FL cells. Genes whose expression is positively enriched in tFL-DLBCL cells are on the left side of the plot, and in tFL-FL on the right.

**Panel B**: Enrichment plot for the top most significant pathways in the Hallmark data-base negatively enriched in the tFL-DLBCL versus tFL-FL cells DE analysis, i.e. enriched in the tFL-FL samples.

**Panel C**: for each pair, enrichment plot for the MYC targets V1 gene sets in Hallmark database for the tFL-FL versus tFL-DLBCL differential expression (HALLMARK_MYC_TARGETS_V1, M5926).

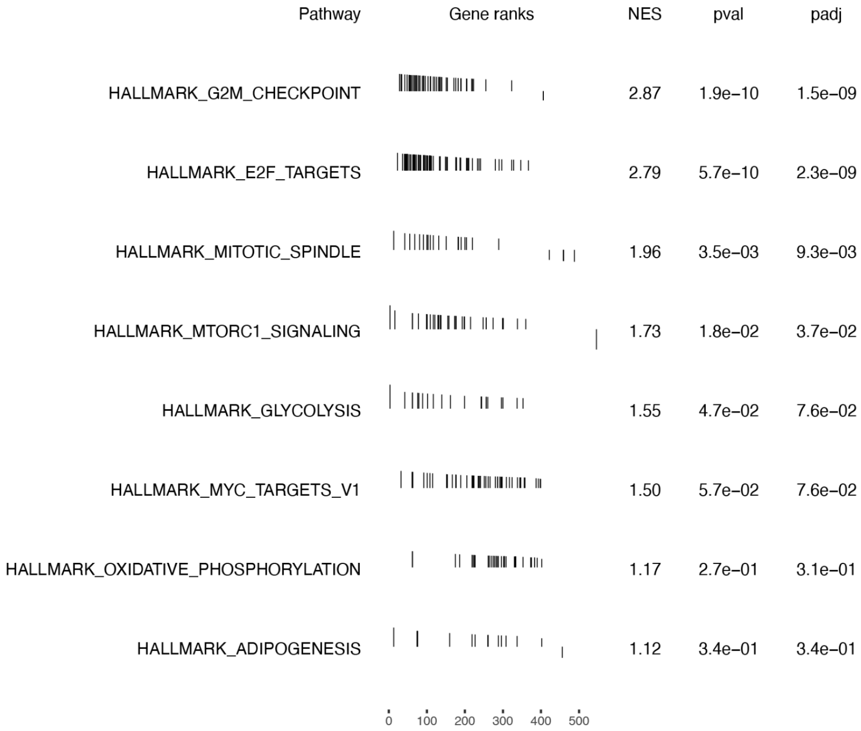


**Supplementary Figure 10.** GSEA from pseudo-bulk differential expression between tFL-FL and tFL-DLBCL timepoint. In this DE, all the cells are included (ie normal B, tumor B, and non-B TME cells) and only 576 genes are differentially expressed. The sensitivity is much lower and biological pathways enriched are less significant. Pathways upregulated in tFL-DLBCL versus tFL-FL using Hallmark data-base: Only G2M and E2F pathways are significantly enriched, reflecting cell proliferation. No pathway were found as significantly downregulated in tFL-DLBCL versus tFL-FL using Hallmark data-base.

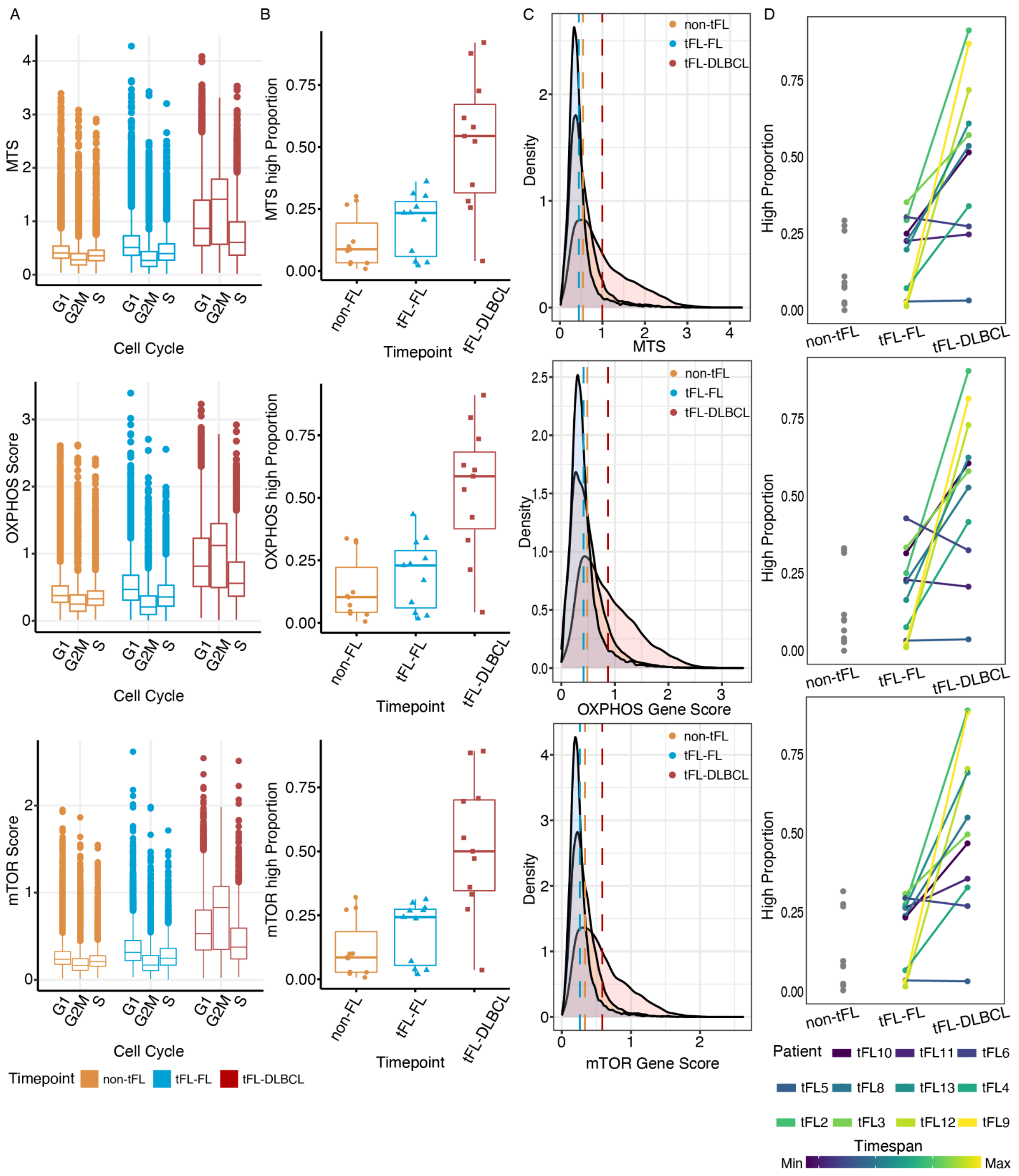


**Supplementary Figure 11.** MTS/OXPHOS/mTOR scores in tFL-FL, tFL-DLBCL and non-tFL samples.

**Panel A:** Boxplot and statistical comparison of mean level of expression between the 3 timepoints (boxplot tFL-FL vs tFL-DLBCL vs non-tFL): the mean MYC target genes score (MTS), was 0.45 vs 0.55 vs 0.99 in non-tFL vs tFL-FL vs tFL-DLBCL (p value <0.001 for each comparison). The mean OXPHOS pathway score was 0.41 vs 0.48 vs 0.87 in non-tFL vs tFL-FL vs tFL-DLBCL (p value <0.001 for each comparison). The mean mTOR pathway score was 0.26 vs 0.33 vs 0.56 in non-tFL vs tFL-FL vs tFL-DLBCL (p value <0.001 for each comparison).

**Panel B**: Boxplot showing within each sample, the proportion of cells with a high score using a 75% quantile threshold.

**Panel C**: Density plot of each score (MTS, OXPHOS and mTOR), the dotted lines represent the mean per data-set.

**Panel D**: Line plot showing the evolution of the proportion of cells with a high "MYC target score" MTS, oxphos or mTOR score, from tFL-FL to tFL-DLBCL, within each pair.

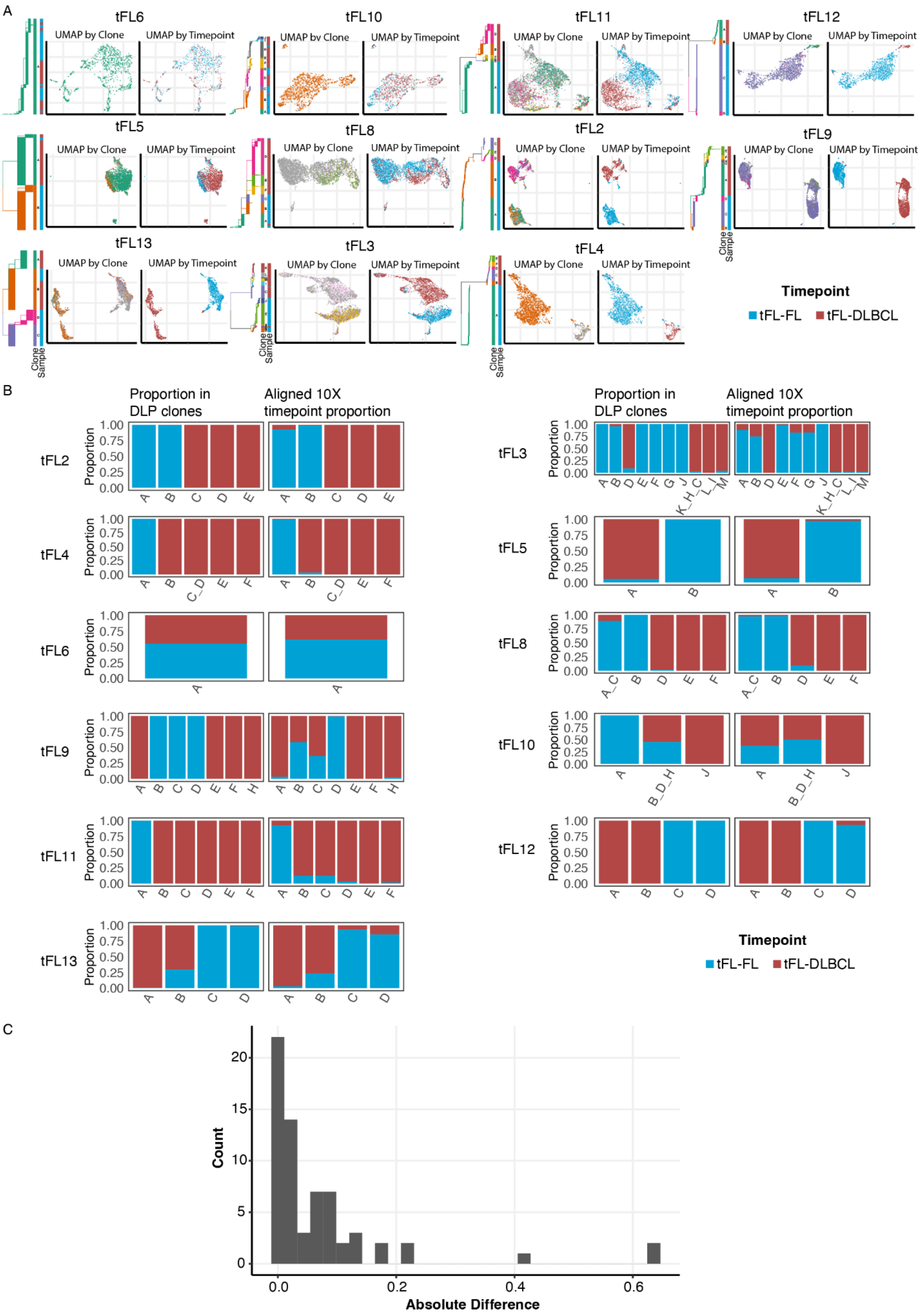
A
tFL6
tFL10
tFL11
tFL12
tFL5
tFL8
tFL2
tFL9
tFL13
tFL3
tFL4
UMAP by Clone
UMAP by Timepoint
Clone
Sample
Timepoint
tFL-FL
tFL-DLBCL
B
Proportion in DLP clones
Aligned 10X timepoint proportion
Proportion
1.00
0.75
0.50
0.25
0.00
C
Count
Absolute Difference
0.0
0.2
0.4
0.6

**Supplementary Figure 12.** clonealign analysis of pairs.

**Panel A:** Each pair is represented, with a phylogeny tree colored by DLP clone, a vertical bar representing the proportion of cells from each DLP clone, and timepoints; and 2 UMAP representations of the re-clustering of each pair. In the first UMAP, cells that are aligned to a DLP clone are colored accordingly, in the second, cells are colored by timepoints (cells from tFL-FL sample are in blue and tFL-DLBCL in red).

**Panel B**: Barplot representation of proportion of tFL-FL versus tFL-DLBCL cells for each pair, within each DLP clone first (left part) and after clone-alignment of the 5GEX cells to a given clone. For each pair, the left barplot shows the proportion of each timepoint by clone (DLP data) and the right barplot shows the proportion of each timepoint in the 5GEX cells that are assigned to a given clone (by clonealign).

**Panel C:** For each clone, we compared the timepoint proportion of cells within DLP clones, and within 5GEX cells aligned to the given DLP clone and represent the absolute difference. A perfect alignment should result in the absence of difference. Given the repartition (mean (0.0845); median (0.0274); range (0-0.635)), N=3 outlier with a difference greater than 0.4 were removed from further analysis, and considered to be failure of clone align: clone A pair 10 (75 cells), clone B and C pair 9 (47 and 339 cells respectively) where assigned cells in fully FL timepoint clone were both from FL and DLBCL timepoint in 10X.

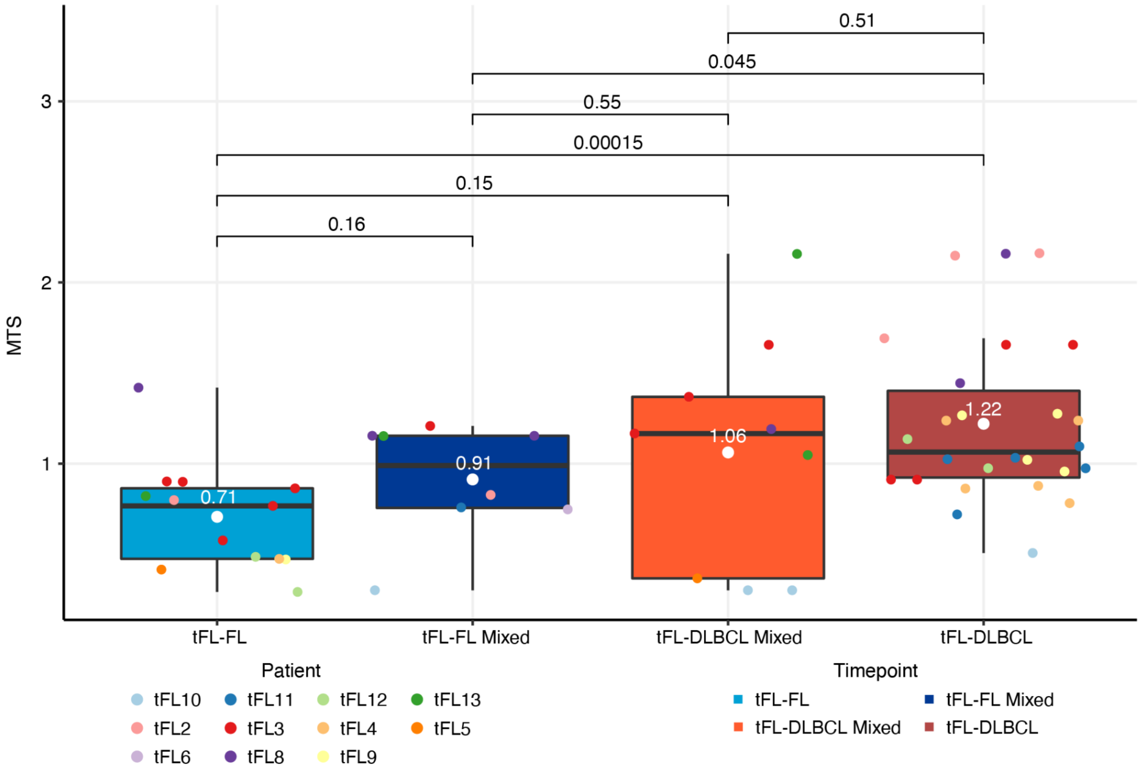


**Supplementary Figure 13.** MYC Target score (MTS) in each DLP clone.

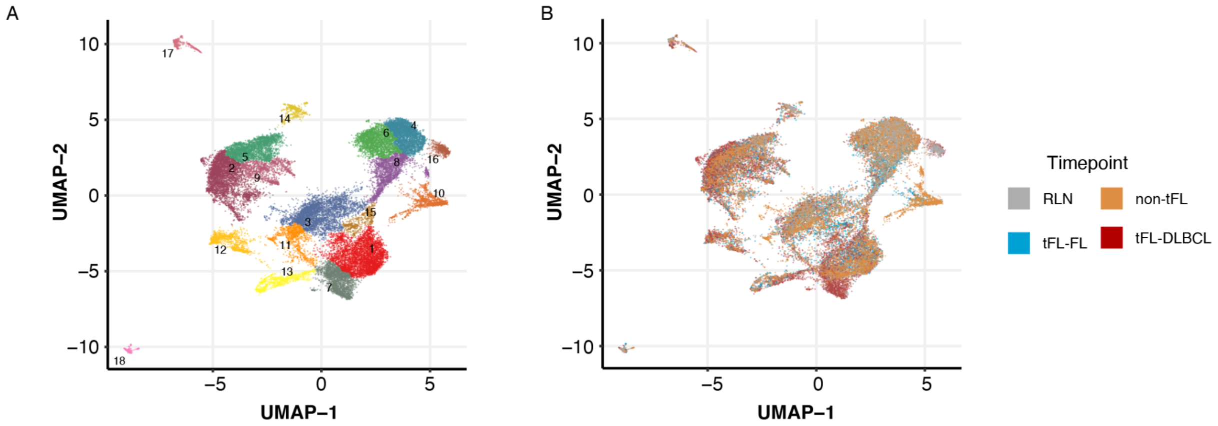


**Supplementary Figure 14.** Tumor microenvironment clustering.

**Panel A**: UMAP visualization of non-B TME cells colored by the 18 clusters.

**Panel B**: UMAP visualization of non-B TME cells colored by timepoints highlighting a shift in UMAP space, from reactive lymph nodes (RLN) cells to tFL-DLBCL cells. RLN cells are colored in grey, non-tFL in orange, tFL-FL in blue and tFL-DLBCL in red.

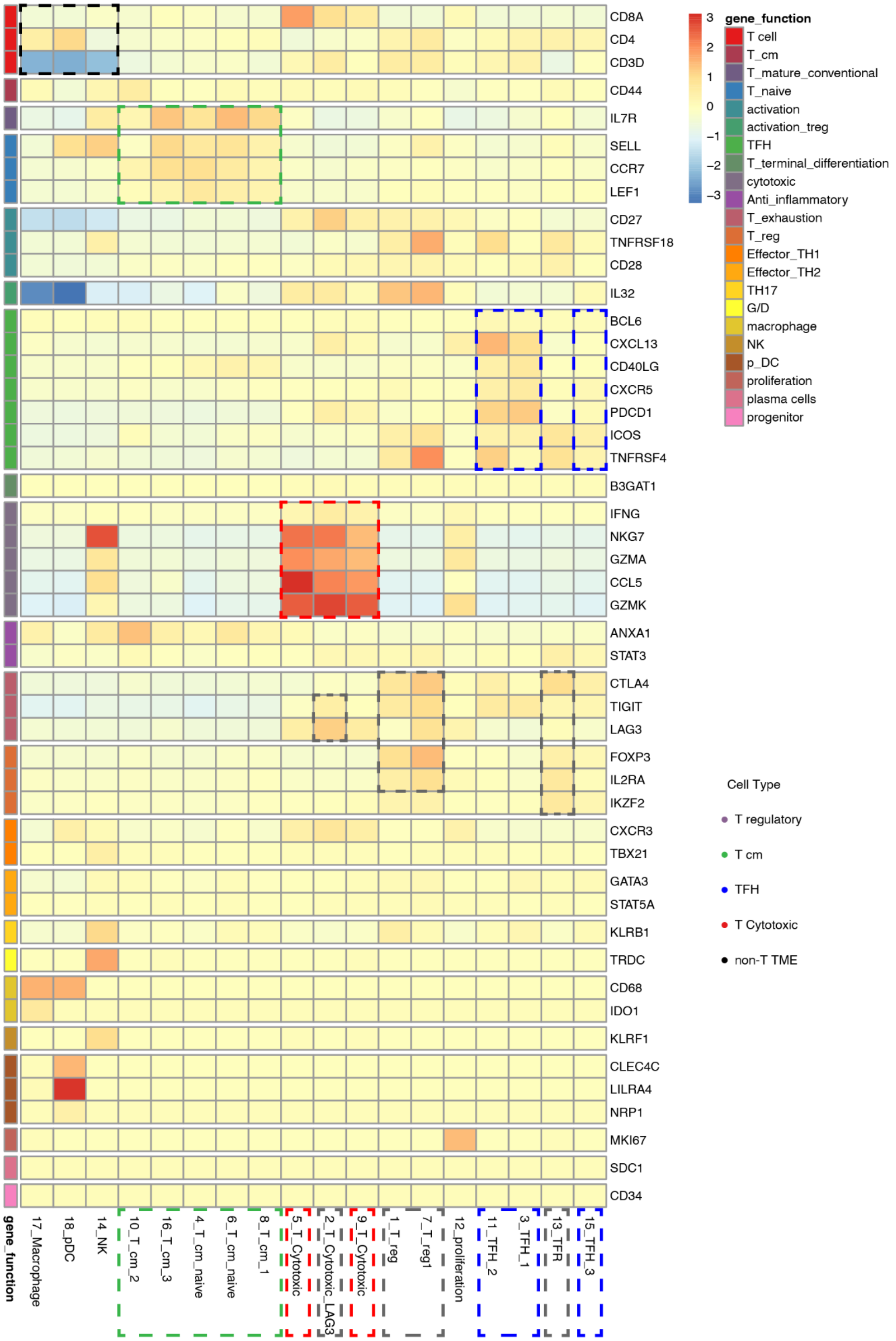


**Supplementary Figure 15**. Gene expression heatmap. Genes were selected based on literature data (Supplementary Table 3). The 5 different naïve/central-memory-like clusters had a high level of expression of *CCR7, LEF1, IL7R, CD52*; one was more enriched in naïve *LEF1* cells (cluster 4), two in cm cells (higher *IL32* expression, cluster 6 and 8), one expressed more AP1 family genes (cluster 10) and one IFN family genes and *SELL* (cluster 16). The 4 T_exh clusters included one CD4+ Treg cluster (cluster 1), one CD4+ Treg1 (high *LAG3* expression, cluster 7), one LAG3+CD8+ cells cluster (cluster 2), and one *ICOS+TNFRSF18+CD4+PD1+* TFR cells cluster (cluster 13). The proliferation cluster (cluster 12) presented a high expression level of *MKi67*, but also *GZMA/K,* and *LAG3*. Finally, the 2 cytotoxic clusters (cluster 5 and 9) were enriched in the expression level of *GZMK/A, CCL5/4* and also *LAG3* or *CD27*, but with a lower level than cluster 2.

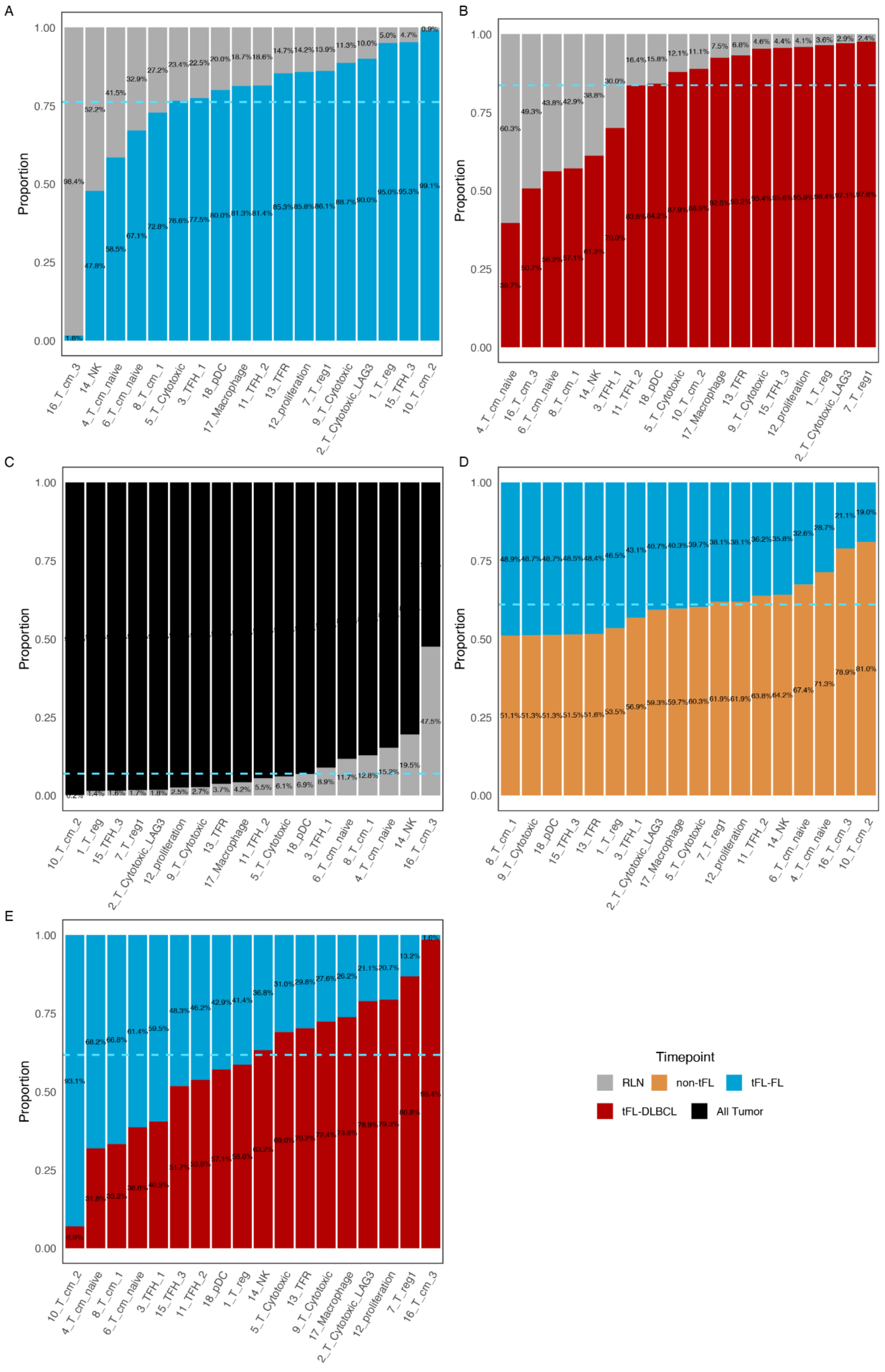
A
B
C
D
E
Proportion
Timepoint
RLN
non-tFL
tFL-FL
tFL-DLBCL
All Tumor

**Supplementary Figure 16.** Barplot for evolution of TME at the cohort level. For each side by side comparison, proportions are represented as the number of cells from each timepoint in the cluster, out of the total number of cells in the cluster. The dotted line represents the expected ratio in each cluster, based on the ratio of total number of non-B TME cells by timepoint.

**Panel A**: tFL-FL versus reactive lymph nodes.

**Panel B**: tFL-DLBCL versus reactive lymph nodes.

**Panel C**: All tumor samples versus reactive lymph nodes.

**Panel D:** tFL-FL versus non-tFL.

**Panel E:** tFL-FL versus tFL-DLBCL.

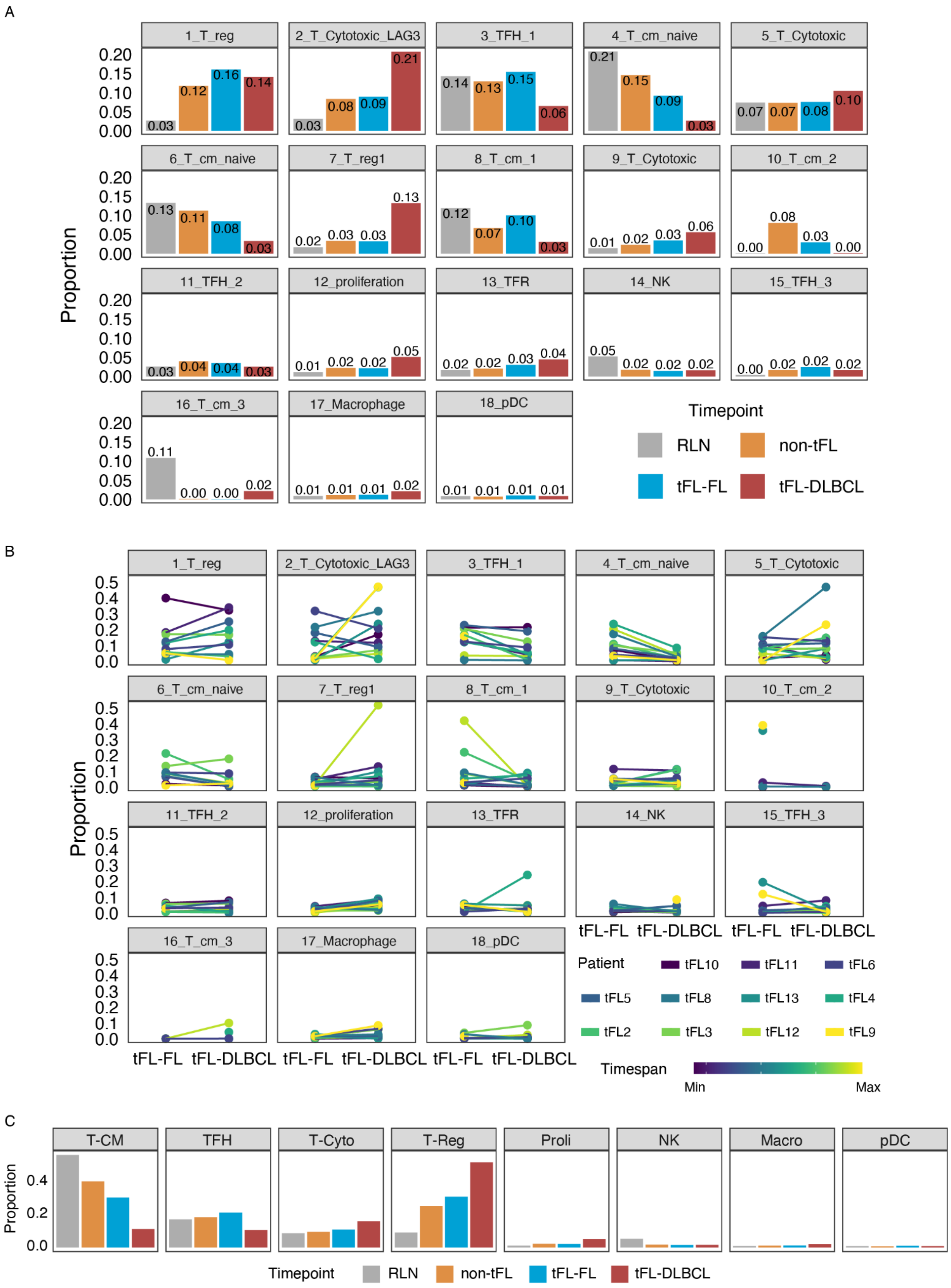
A
B
C
Proportion
1_T_reg
2_T_Cytotoxic_LAG3
3_TFH_1
4_T_cm_naive
5_T_Cytotoxic
6_T_cm_naive
7_T_reg1
8_T_cm_1
9_T_Cytotoxic
10_T_cm_2
11_TFH_2
12_proliferation
13_TFR
14_NK
15_TFH_3
16_T_cm_3
17_Macrophage
18_pDC
Timepoint
RLN
non-tFL
tFL-FL
tFL-DLBCL
Patient
tFL10
tFL11
tFL6
tFL5
tFL8
tFL13
tFL4
tFL2
tFL3
tFL12
tFL9
Timespan
Min
Max
T-CM
TFH
T-Cyto
T-Reg
Proli
NK
Macro
pDC

**Supplementary Figure 17.** Evolution of TME composition, within cohorts and in each pair.

**Panel A**: Barplot representing the proportion of cells from each TME cluster out of all cells from RLN, non-tFL, tFL_FL and tFL-DLBCL samples.

**Panel B:** Line plot representing the evolution of proportion of cells from each TME cluster out of all tFL-FL and tFL-DLBCL cells. The color scale represents the time elapsed between the 2 biopsies.

**Panel C:** Barplot representing the proportion of cells from each TME cluster out of all cells from RLN, non-tFL, tFL_FL and tFL-DLBCL samples. T cell clusters are grouped for further analysis: T-cm represent cells included in cluster with T central memory markers identified (ie cluster 4,6,8,10,16), TFH cells included in cluster with TFH markers identified (ie cluster 3,11,15), T-cyto cells included in cluster with T cytotoxic markers identified (ie cluster 5, 9), T-exh cells included in cluster with exhaustion or regulatory markers identified (ie cluster 1,2,7,13). Cells included in cluster 12 (characterized by high MKI67 expression level), NK (natural killer cells cluster 14), Macro (macrophage cells cluster 17) and pDC (plasmacytoid dendritic cells cluster 18) are shown.

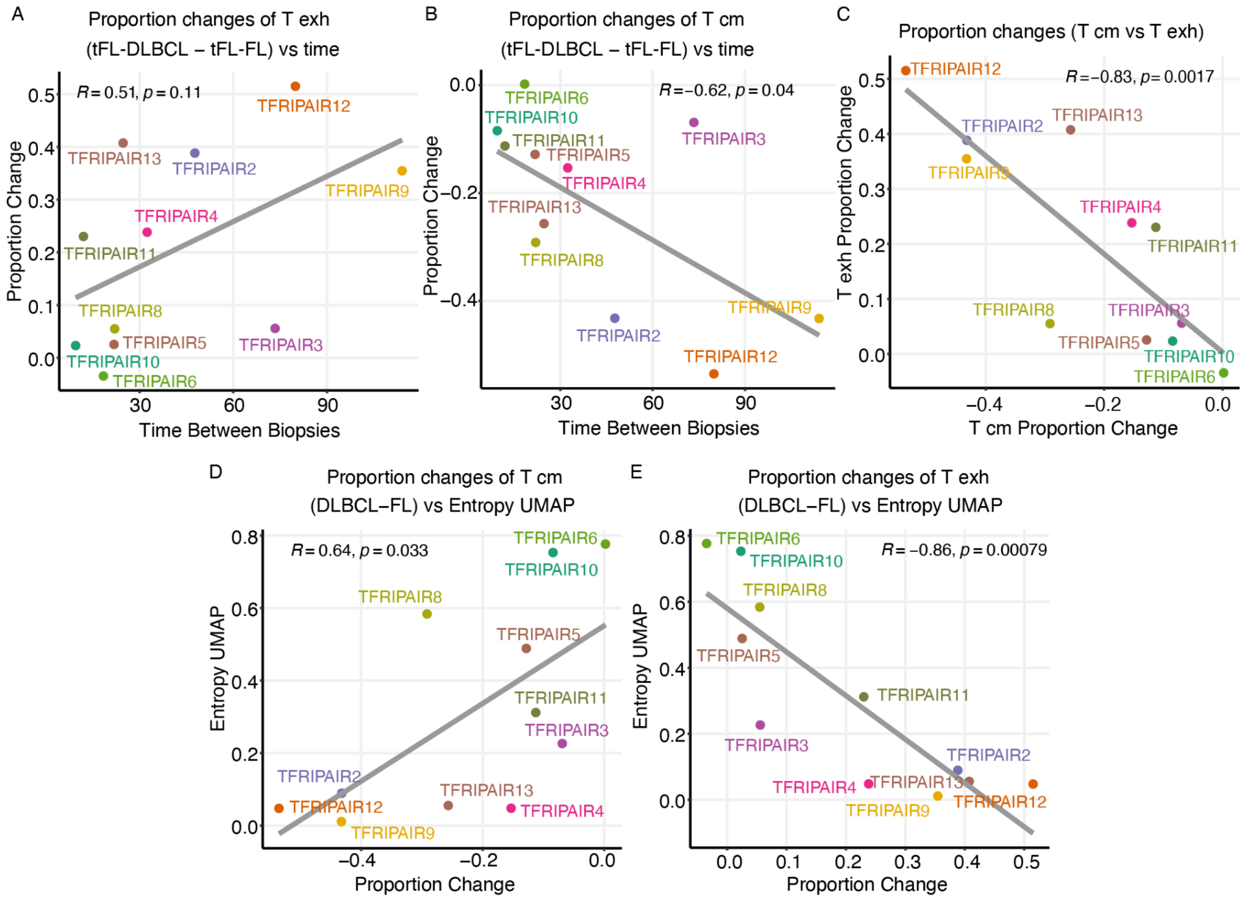


**Supplementary Figure 18.** Magnitude of tumor B cells versus TME evolution in each pair and according to time between the tFL-FL and tFL-DLBCL biopsies. The proportion changes (defined as the changes in cluster proportion out of all TME cells in each timepoints) between tFL-FL and tFL-DLBCL for T_exh and Tcm clusters is represented versus time elapsed between the 2 biopsies and the UMAP entropy.

**Panel A:** Proportion changes of T_exh clusters versus time between the 2 biopsies, p value = 0.12

**Panel B:** Proportion changes of Tcm clusters versus time between the 2 biopsies, p value=0.038

**Panel C:** Proportion changes of Tcm clusters versus T_exh clusters;

**Panel D:** Correlation between UMAP entropy and proportion of change in the Tcm population

**Panel E:** Correlation between UMAP entropy and proportion of change in T_exh clusters

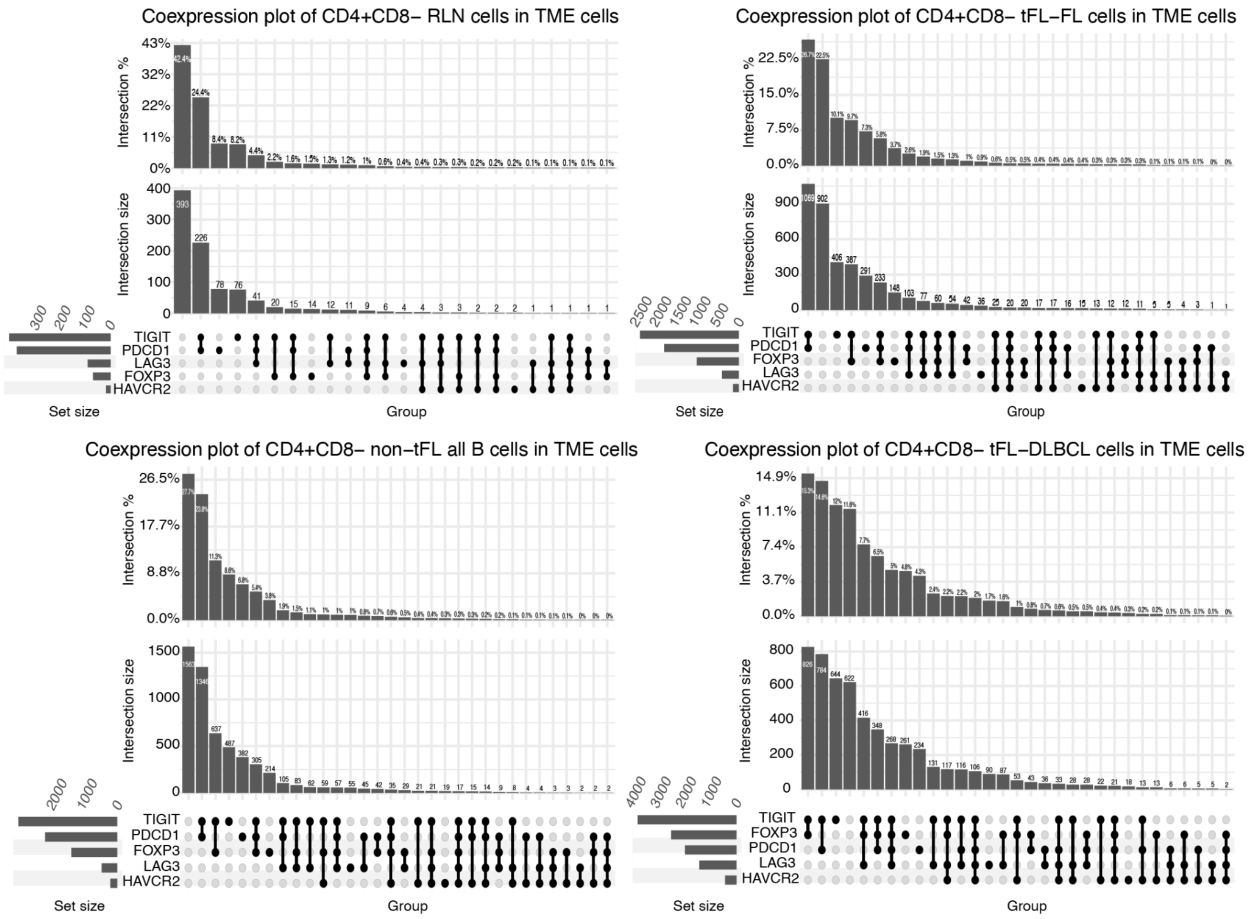


**Supplementary Figure 19.** Co-expression plot of regulatory/exhaustion genes from 10X single cell gene expression data, within CD4 cells, in RLN, nontFL, tFL-FL and tFL-DLBCL samples (CD4/CD8 double positive cells were excluded from this analysis). The plot highlights an increase in FOXP3-TIGIT and LAG3 positive cells in tFL-DLBCL samples, as compared to FL, with a decrease in the TIGIT-PD1 population. Of note, the global increase in the co-expression of genes associated with an exhausted phenotype, within CD4 cells, is gradual from RLN to FL, tFL-FL and tFL-DLBCL.

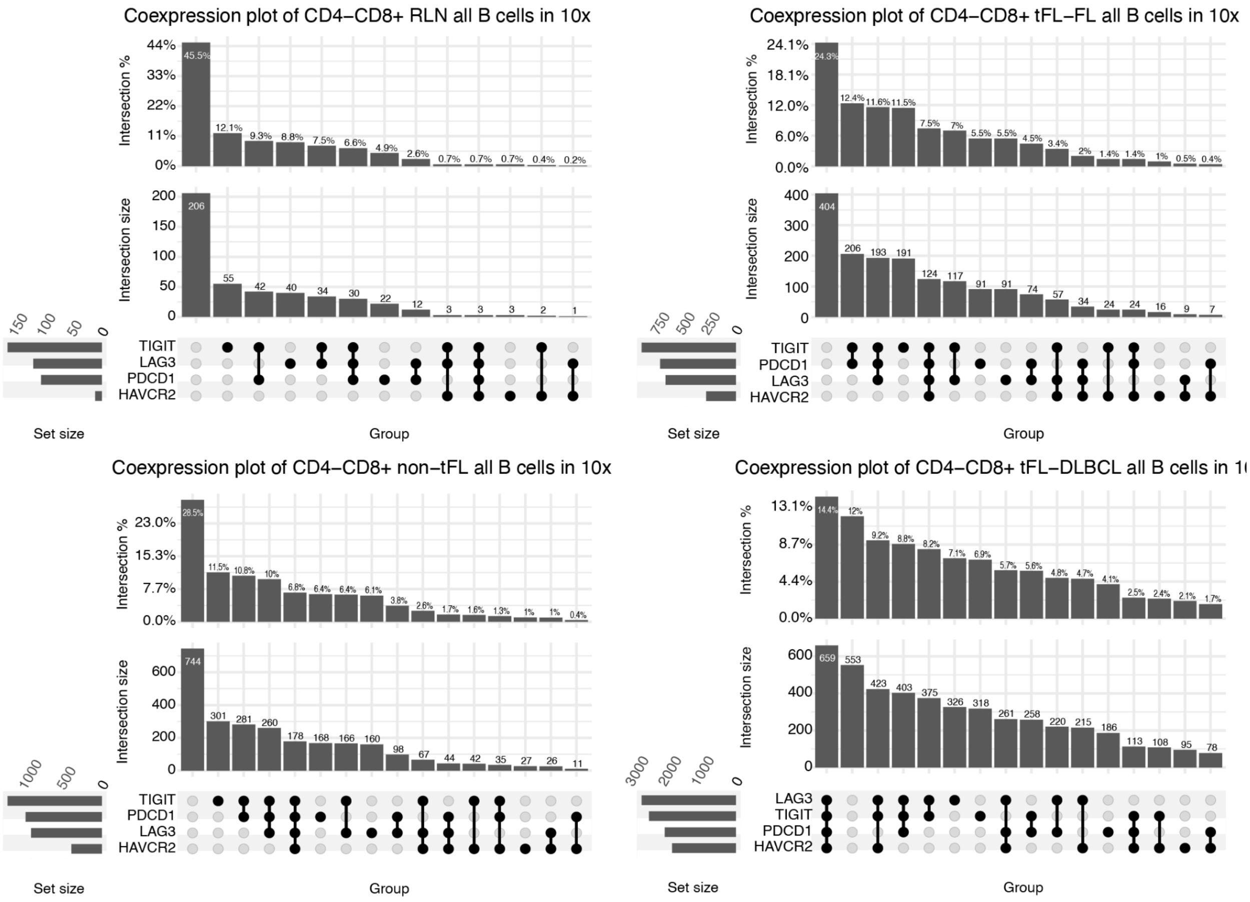


**Supplementary Figure 20.** Co-expression plot of regulatory/exhaustion genes from 10X single cell gene expression data, within CD8 cells, in RLN, nontFL, tFL-FL and tFL-DLBCL samples (CD4/CD8 double positive cells were excluded from this analysis). The plot highlights a clear shift in the co-expression of LAG3+, TIGIT+, PDCD1+, HAVCR2+ with an important increase in the number and proportion of cells with a "full" exhausted CD8 phenotype in tFL-DLBCLs versus FLs. These cells are almost absent in RLN samples. Of note, the global increase in the co-expression of genes associated with an exhausted phenotype, within CD8 cells, is gradual from RLN to FL, tFL-FL and tFL-DLBCL.

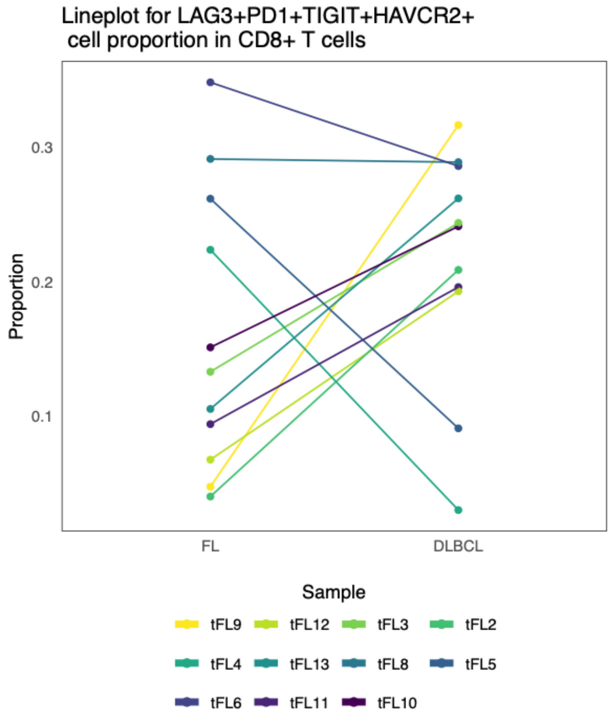


**Supplementary Figure 21.** Lineplot for proportion change of LAG3/PD1/TIGIT/HAVCR2 in CD8 cells between tFL-FL and tFL-DLBCL.

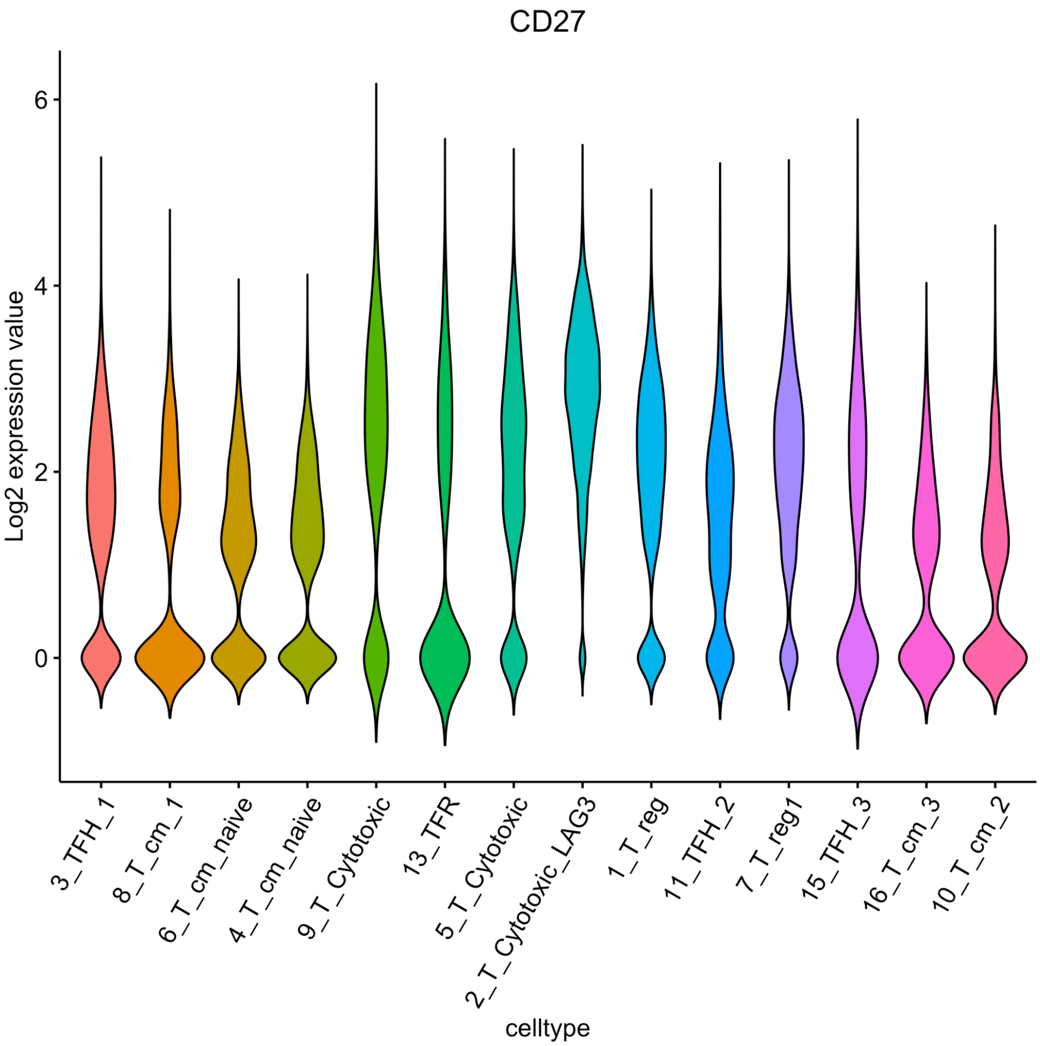


**Supplementary Figure 22.** CD27 expression in the different TME clusters in 10X.

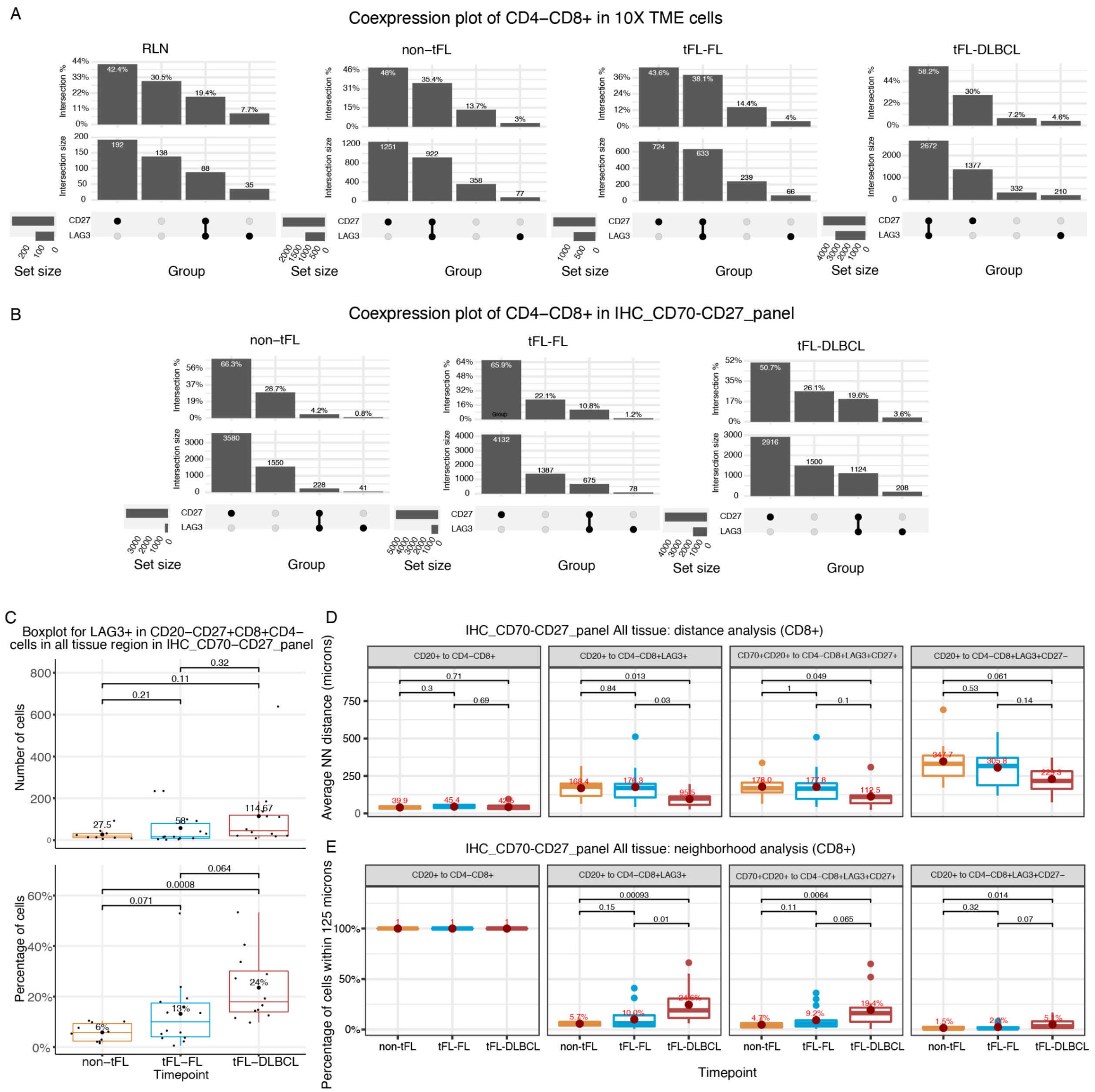


**Supplementary Figure 23.** LAG3 and CD27 expression in CD8 cells, and distance to CD20/CD70 cells during transformation.

**Panel A**:10X coexpression plot of CD27 and LAG3 within CD8+CD4- and CD20- cells.

**Panel B**: MC-IHC plot of CD27 and LAG3 within CD8+CD4- and CD20- cells.

**Panel C**: Percentage of LAG3 positivity within CD8+CD27+ cells in non-tFL, FL and DLBCL.

**Panel D**: Distance analysis between B cells (CD20+) and CD8+ cells with a given phenotype.

**Panel E**: Percentage of CD8LAG3+, CD8LAG3CD27+ cells out of CD8 cells within 125 microns from CD20/CD70 cells, and CD8LAG3C27- cells out of CD8 cells within 125 microns from CD20 cells.

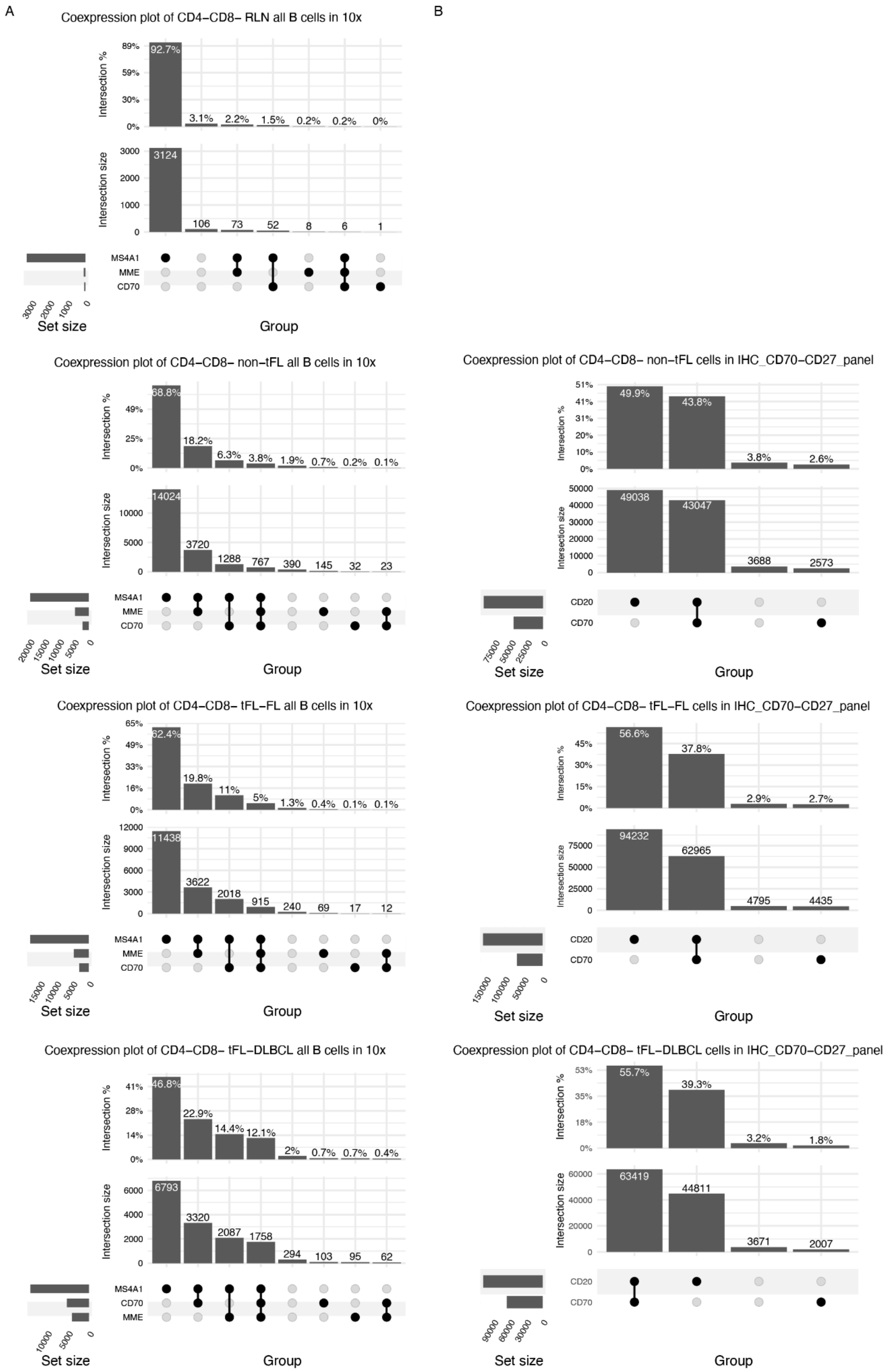
A
Coexpression plot of CD4−CD8− RLN all B cells in 10x
Intersection %
92.7%
3.1%
2.2%
1.5%
0.2%
0.2%
0%
Intersection size
3124
106
73
52
8
6
1
MS4A1
MME
CD70
Set size
Group
B
Coexpression plot of CD4−CD8− non−tFL all B cells in 10x
68.8%
18.2%
6.3%
3.8%
1.9%
0.7%
0.2%
0.1%
14024
3720
1288
767
390
145
32
23
Coexpression plot of CD4−CD8− non−tFL cells in IHC_CD70−CD27_panel
49.9%
43.8%
3.8%
2.6%
49038
43047
3688
2573
CD20
CD70
Coexpression plot of CD4−CD8− tFL−FL all B cells in 10x
62.4%
19.8%
11%
5%
1.3%
0.4%
0.1%
0.1%
11438
3622
2018
915
240
69
17
12
Coexpression plot of CD4−CD8− tFL−FL cells in IHC_CD70−CD27_panel
56.6%
37.8%
2.9%
2.7%
94232
62965
4795
4435
Coexpression plot of CD4−CD8− tFL−DLBCL all B cells in 10x
46.8%
22.9%
14.4%
12.1%
2%
0.7%
0.7%
0.4%
6793
3320
2087
1758
294
103
95
62
Coexpression plot of CD4−CD8− tFL−DLBCL cells in IHC_CD70−CD27_panel
55.7%
39.3%
3.2%
1.8%
63419
44811
3671
2007

**Supplementary Figure 24.** CD70 expression in  B cells.

**Panel A:** 10X co-expression plot of MS4A1, MME and CD70 in all B cells clusters (CD4-CD8- cells were selected within the B cell clusters (cf Supplementary Figure 4B, including both normal and malignant B cells)).

**Panel B:** Co-expression of CD70 and CD20 in immuno-histo-chemestry, in tFL-FL, tFL-DLBCL and non-tFL samples. As for panel A, only cells with a negative expression for both CD4 and CD8 were analyzed.

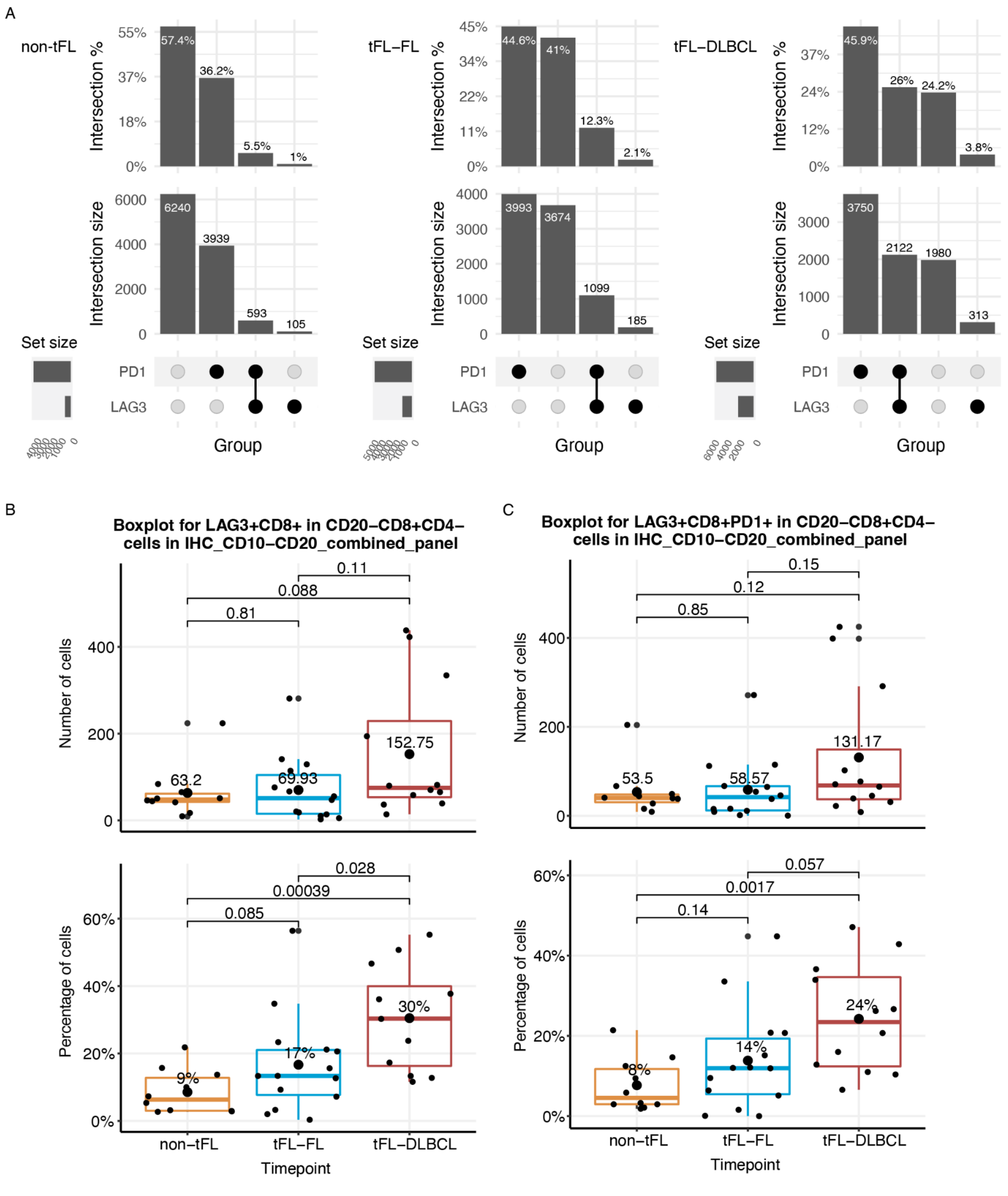


**Supplementary Figure 25.** LAG3 and PD1 expression in CD8 cells in FL and during transformation.

**Panel A**: Co-expression plot for LAG3 and PD1 expression, within CD8+ (and CD4-CD20-) cells, in MC- IHC in the 3 different cohorts (FL-nontFL, tFL-FL and tFL-DLBCL).

**Panel B**: IHC expression of LAG3 in CD8 positive cells (and CD4-CD20-) in the 3 different cohorts (FL-nontFL, tFL-FL and tFL-DLBCL). The % of LAG3 + cells among CD8 + cells is represented.

**Panel C**: IHC expression of LAG3PD1 in CD8 positive (and CD4-CD20-) cells in the 3 different cohorts (FL-nontFL, tFL-FL and tFL-DLBCL). The % of LAG3PD1 co-expressing + cells among CD8 + cells is represented.

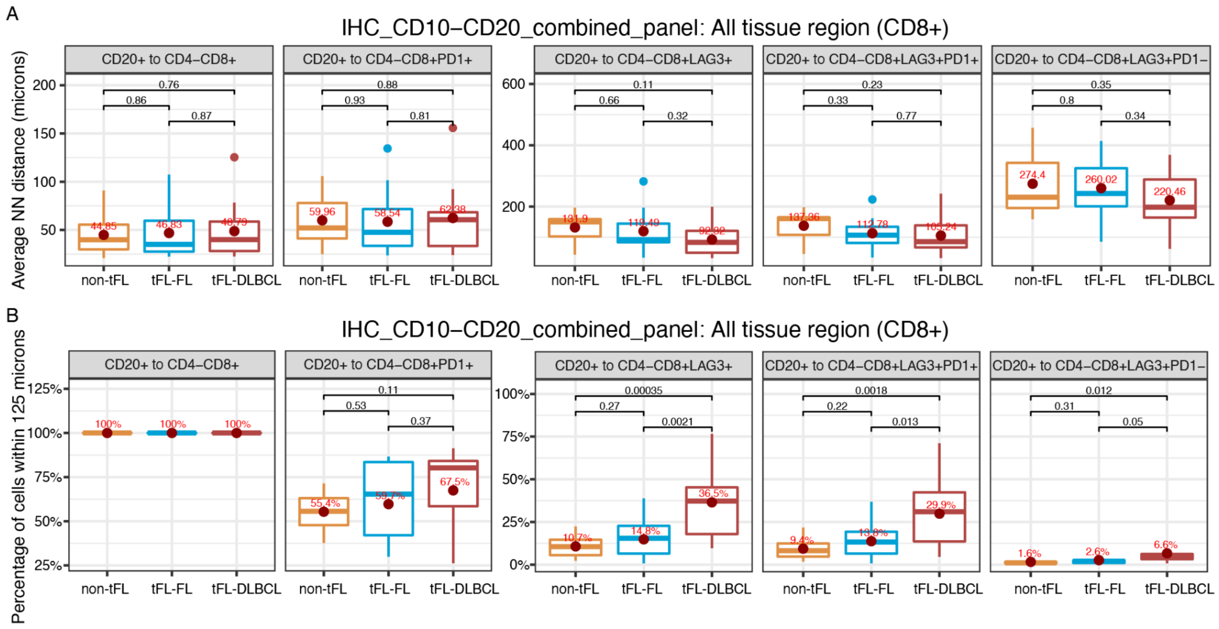


**Supplementary Figure 26.** Distance and neighborhood analysis between CD8+/LAG3+ cells and CD20+ cells. Within all tissue regions, the distance and neighborhood analysis were performed between CD8+, CD8+PD1+, CD8+LAG3+, CD8+LAG3+PD1+, CD8+LAG3+PD1- and B cells. CD8+ cells were selected as CD4- and CD20-.

**Panel A**: Distance between the different types of CD8 cells and CD20 cells, within each cohort. The dot represents the mean distance and the bar the median. The p value represents the statistical testing between the groups.

**Panel B:** CD8 cells and CD20 neighborhood analysis representing the percentage of CD8 cells with the given phenotype within CD8+ cells, in 125 microns from any B cells.

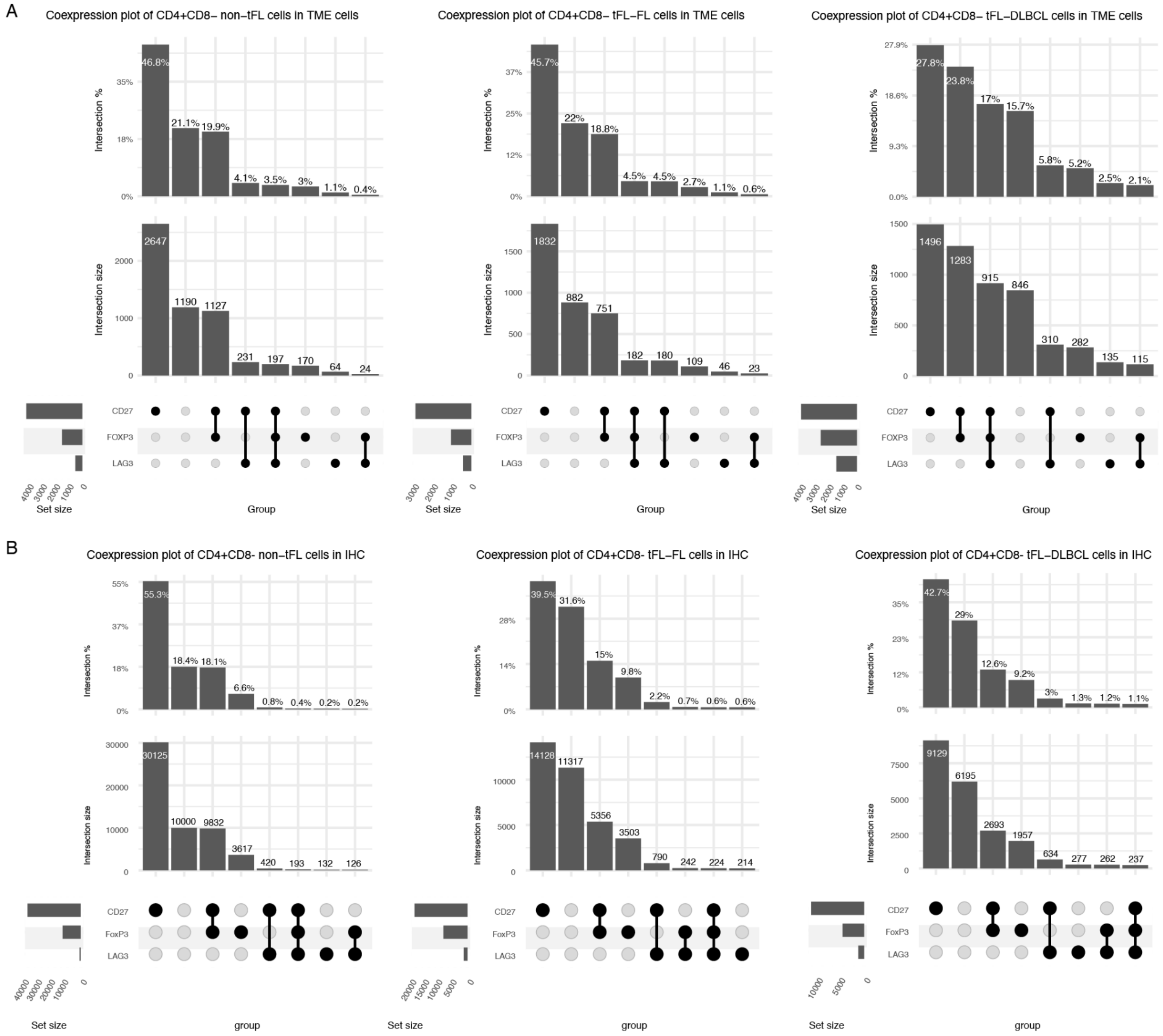


**Supplementary Figure 27.** Coexpression plots of CD4+ cells in 10X and IHC.

**Panel A**: 10X coexpression plot of *CD27, FOXP3* and *LAG3* within CD8-CD4+ and CD20- cells.

**Panel B**: MC-IHC plot of CD27, FOXP3 and LAG3 within CD8-CD4+ and CD20- cells.

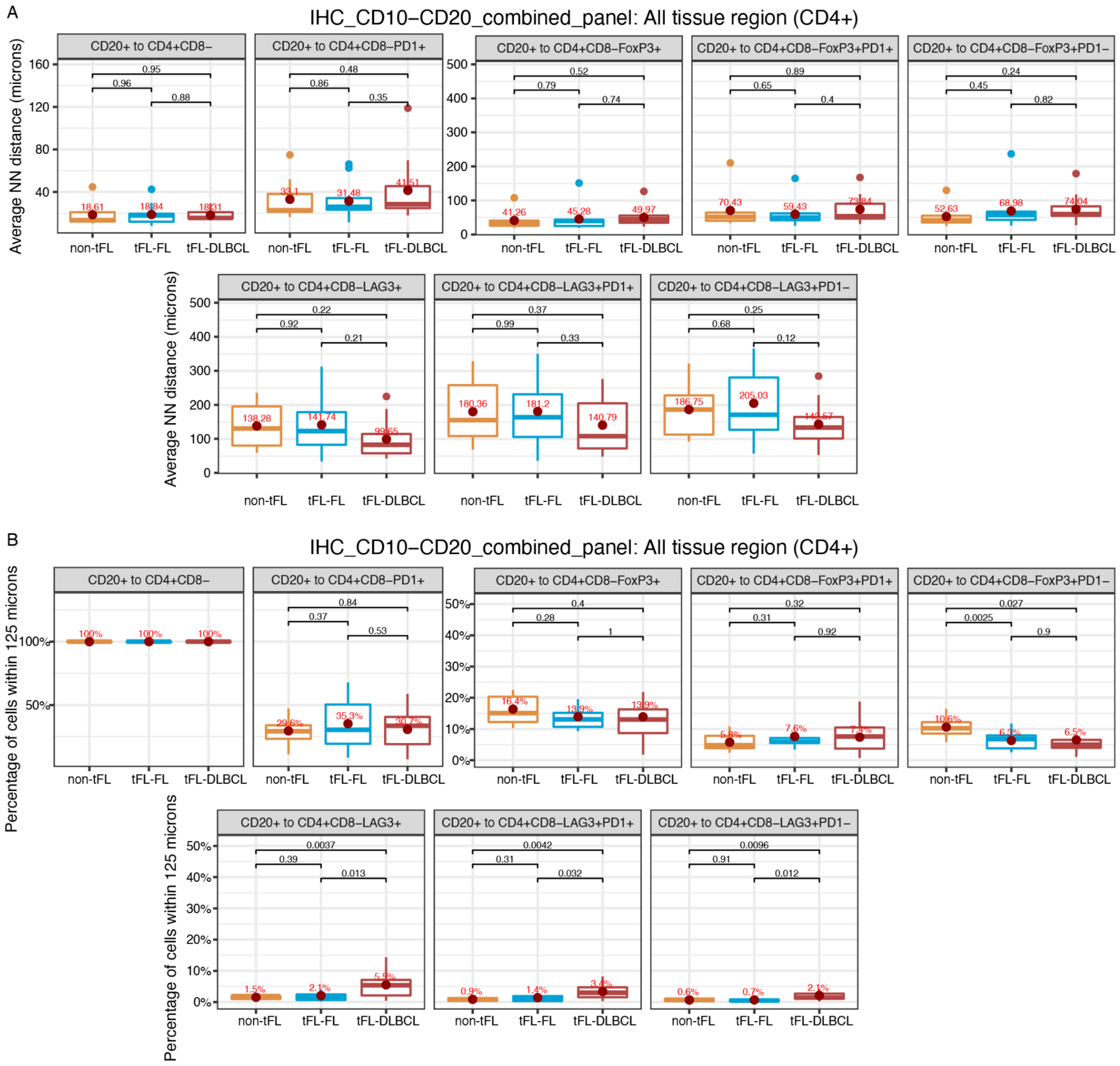


**Supplementary Figure 28.** Distance and neighborhood analysis between CD4+ cells and B cells. Within all tissue region, the distance and neighborhood analysis were performed between CD4+, CD4+PD1+, CD4+LAG3+, CD4+LAG3+PD1+, CD4+LAG3+PD1-, CD4+FOXP3+PD1+, CD4+FOXP3+PD1-, and B cells. CD4+ cells were selected as CD8- and CD20-.

**Panel A** : Distance between the different types of CD4 cells and CD20 cells, within each cohort. The dot represents the mean distance and the bar the median. The p value represents the statistical testing between the groups.

**Panel B**: CD4 cells and CD20 neighborhood analysis representing the percentage of CD4 cells with the given phenotype within CD4+ cells, in 125 microns from any B cells.

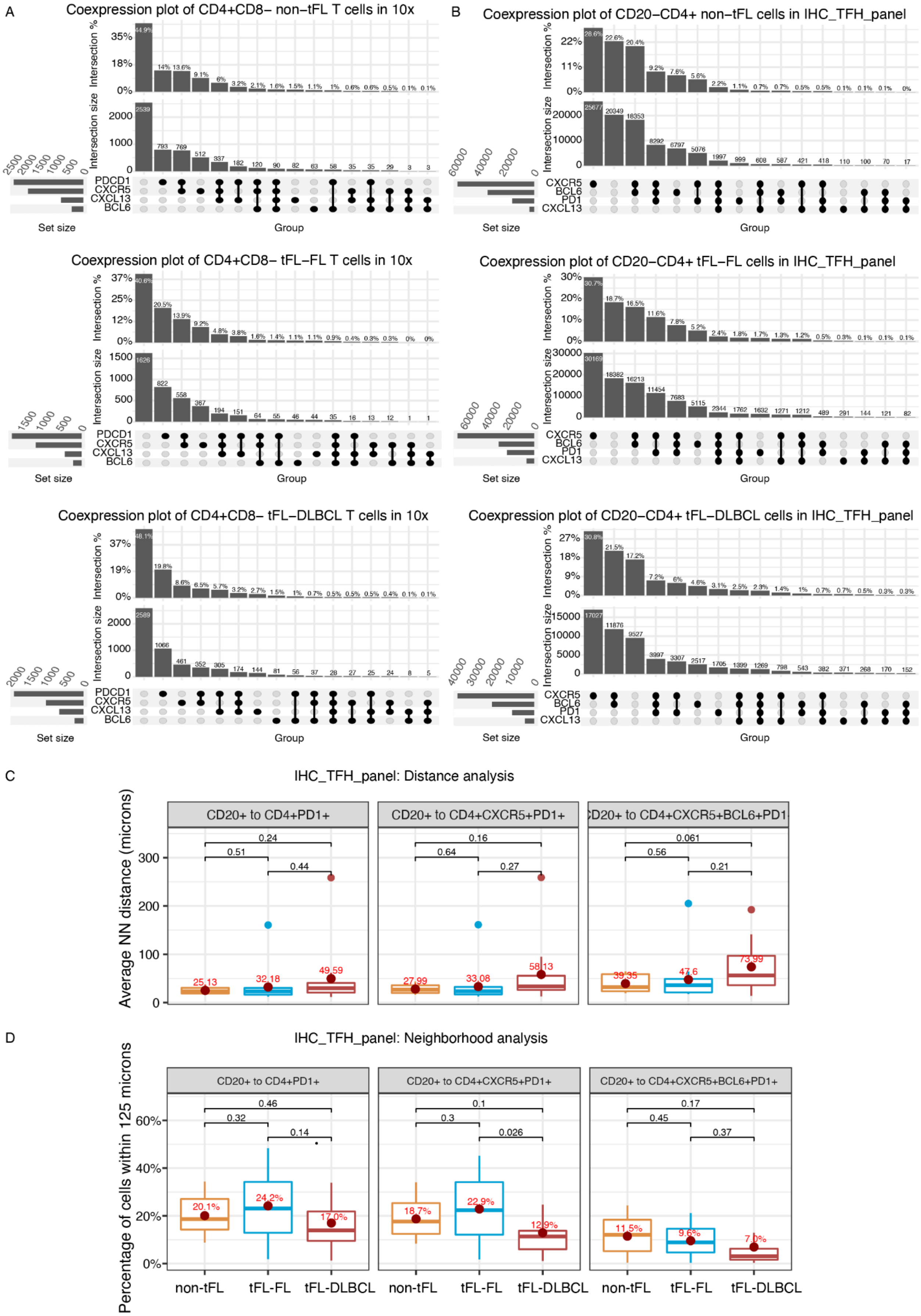

A
Coexpression plot of CD4+CD8− non−tFL T cells in 10x
B
Coexpression plot of CD20−CD4+ non−tFL cells in IHC_TFH_panel
Coexpression plot of CD4+CD8− tFL−FL T cells in 10x
Coexpression plot of CD20−CD4+ tFL−FL cells in IHC_TFH_panel
Coexpression plot of CD4+CD8− tFL−DLBCL T cells in 10x
Coexpression plot of CD20−CD4+ tFL−DLBCL cells in IHC_TFH_panel
Intersection %
Intersection size
Set size
Group
PDCD1
CXCR5
CXCL13
BCL6
PD1
C
IHC_TFH_panel: Distance analysis
Average NN distance (microns)
CD20+ to CD4+PD1+
CD20+ to CD4+CXCR5+PD1+
CD20+ to CD4+CXCR5+BCL6+PD1+
D
IHC_TFH_panel: Neighborhood analysis
Percentage of cells within 125 microns
non-tFL
tFL-FL
tFL-DLBCL

**Supplementary Figure 29.** TFH component in FL and transformation.

**Panel A**: Co-expression plot for CD4 cells (and CD8 negative) with a TFH phenotype in 10X data (ie expression of BCL6, CXCR5, PD1, CXCL13), within the different cohorts (non-tFL, tFL-FL, tFL-DLBCL).

**Panel B**: Co-expression plot of CD4 cells with a TFH phenotype (ie surface expression of BCL6, CXCR5, PD1, CXCL13), within the different cohorts (non-tFL, tFL-FL, tFL-DLBCL).

**Panel C**: Distance analysis with mean distance between B cells (CD20+) and TFH cells with a given phenotype showing that TFH cells are more distant to B cells in tFL-DLBCL as compared to tFL-FL samples.

**Panel D**: percentage of TFH cells out of CD4 cells within 125 microns from B cells, showing a decrease in TFH cells in tFL-DLBCL samples as compared to FLs.

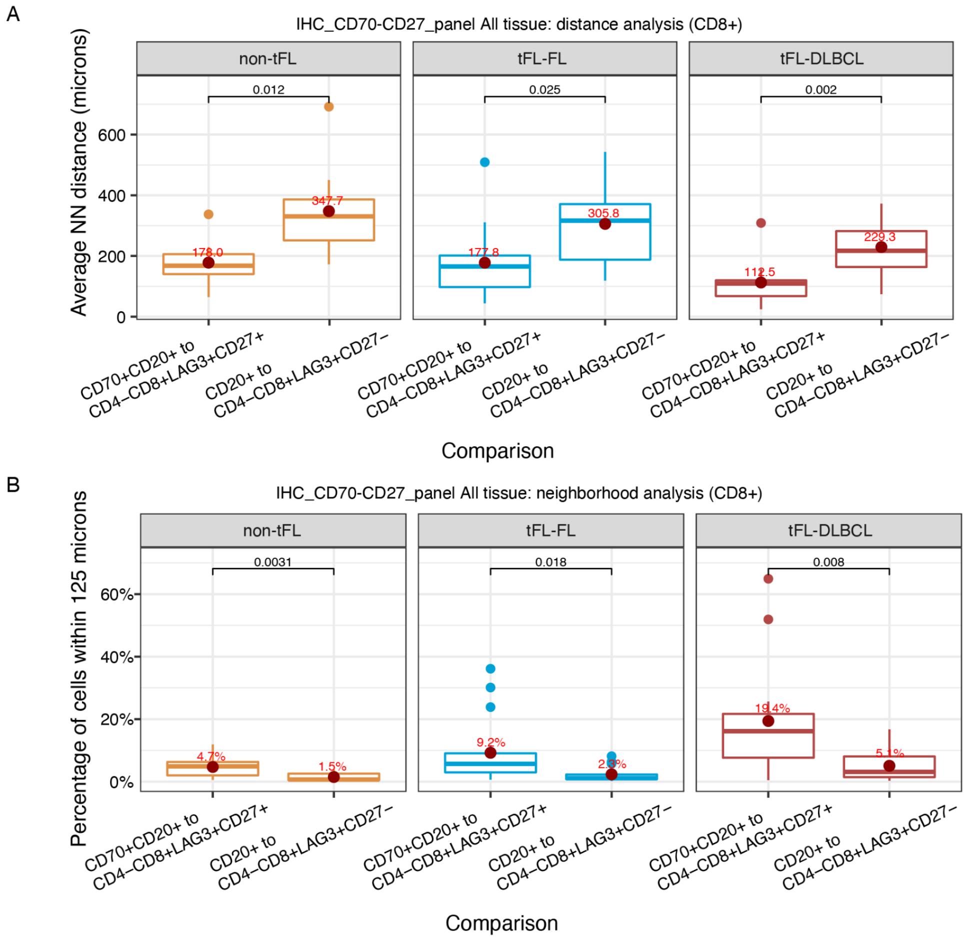


**Supplementary Figure 30**. Distance and neighborhood analysis of CD27LAG3CD8 cells to CD20CD70 cells.

**Panel A:** Analysis, within each timepoint, of the distance between CD20CD70 positive cells and CD27CD8LAG3 positive cells, versus CD27 negative cells.

**panel B:** Percentage of CD8LAG3CD27 cells out of CD8 cells, within 125 microns from CD20CD70 cells.

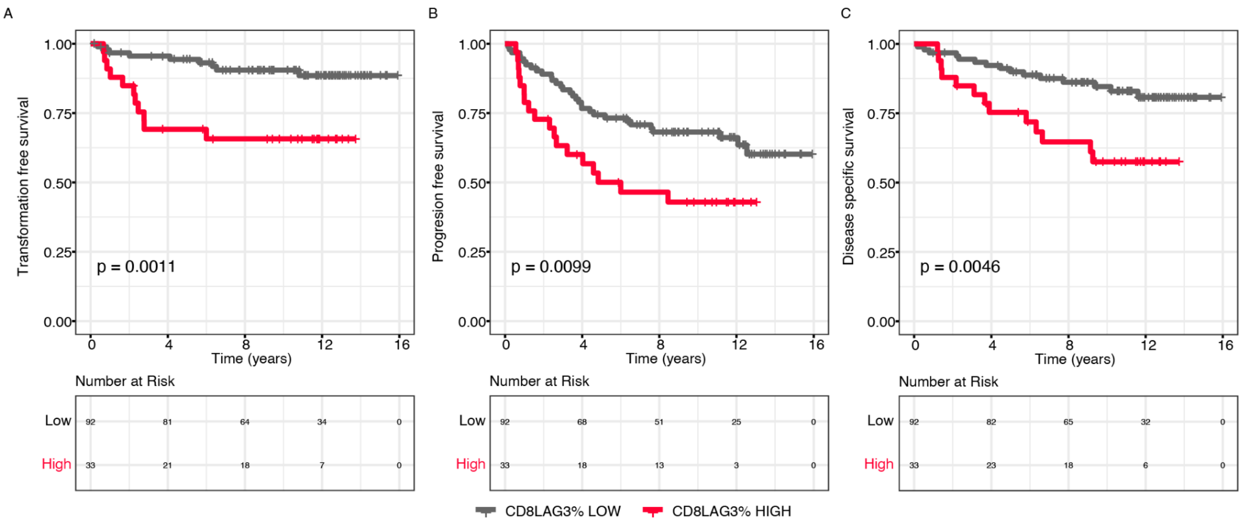


**Supplementary Figure 31:** Survival analysis in R-CVP cohort (rituximab, cyclophosphamide, vincristine, prednisone). N=125 patients were included in the analysis.

**Panel A:** Survival curves for time to transformation (transformation free survival).

The cohort is grouped based on high versus low percentage of CD8LAG3 positive cells out of TME cells (ie CD20 negative cells) in pre-treatment biopsies. Threshold was defined as the highest quartile cutoff value (1.1%).

**Panel B:** Survival curves for time to progression (progression free survival).

**Panel C:** Survival curves for disease specific survival.

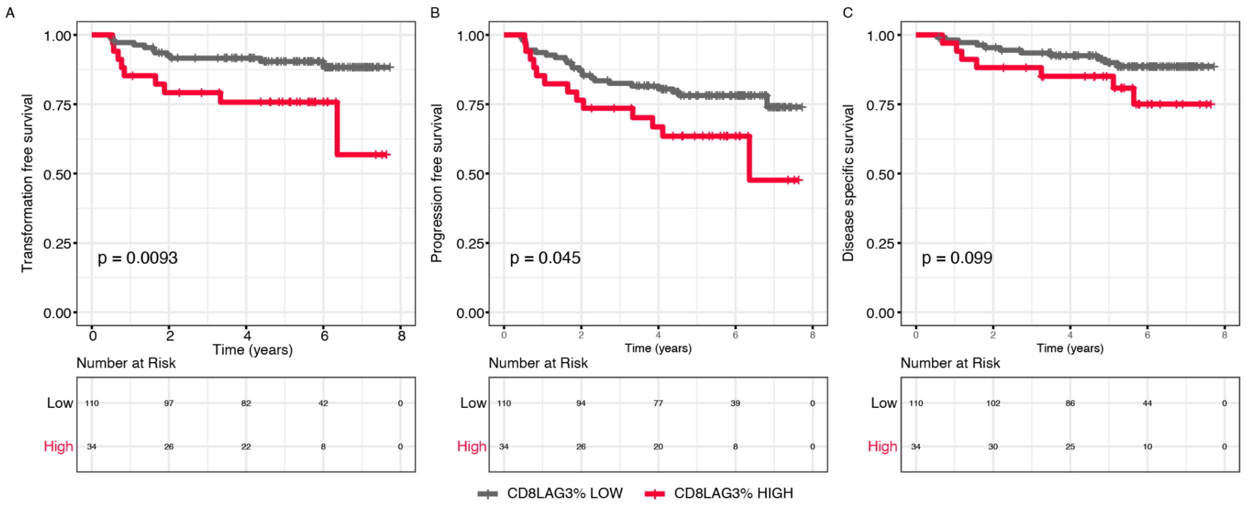


**Supplementary Figure 32.** Survival analysis in Rituximab-Bendamustine cohort.

N=144 patients are included in the analysis.

**Panel A:** Survival curves for time to transformation.

The cohort is grouped based on high versus low percentage of CD8LAG3 positive cells out of TME cells (ie CD20 negative cells) in pre-treatement biopsies. Threshold was defined as the highest quartile cutoff value (1.8%).

**Panel B:** Survival curves for time to progression.

**Panel C**: Survival curves for disease specific survival.

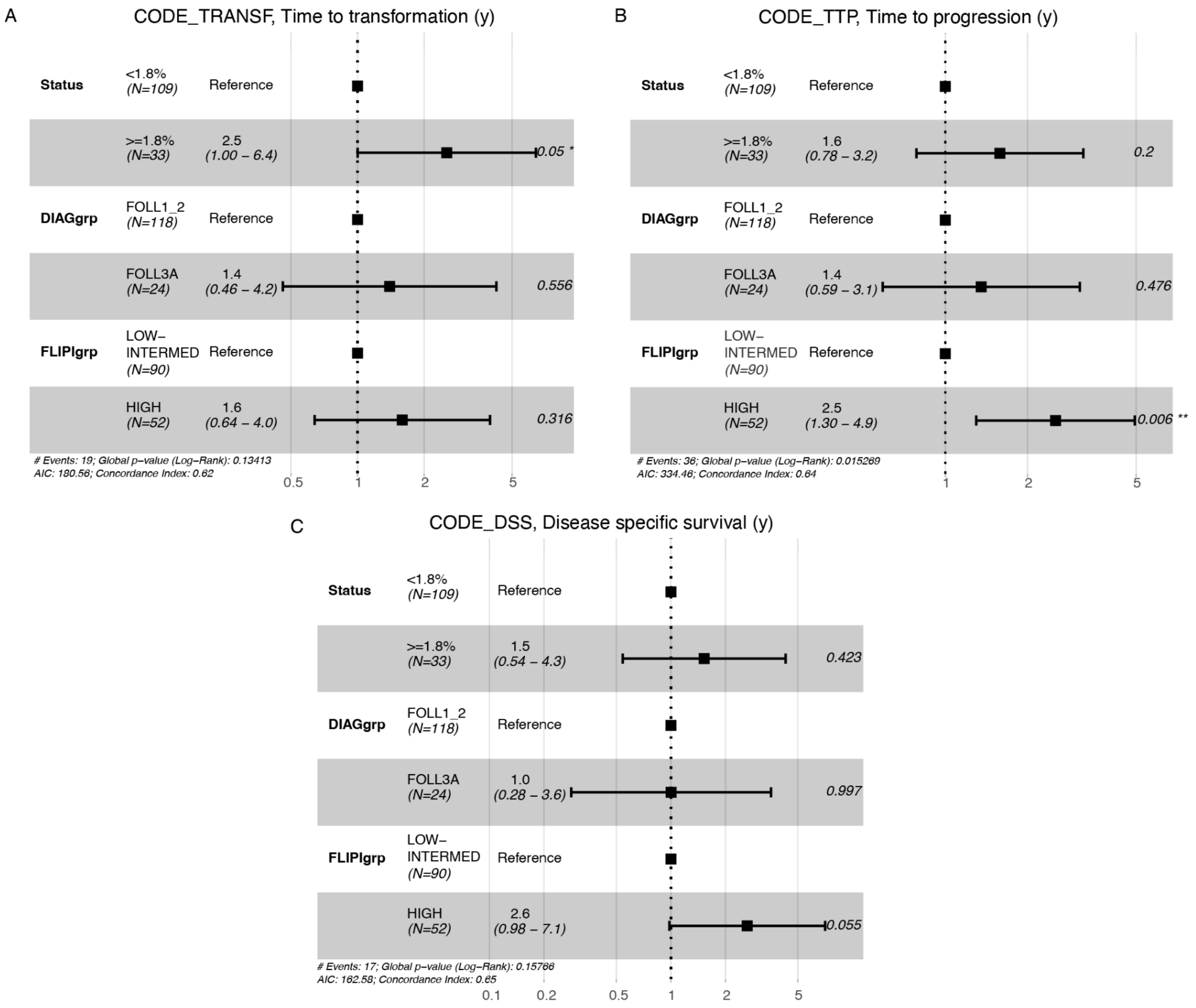


**Supplementary Figure 33:** Multivariate analysis in R-B cohort. The model included FLIPI score (low, versus intermediate, versus high), pathological grading (1-2 versus 3A) and CD8LAG3 infiltrate (low versus high)

**Panel A:** Cox model for time to transformation.

**Panel B:** Cox model for time to progression.

**Panel C:** Cox model for disease specific survival.

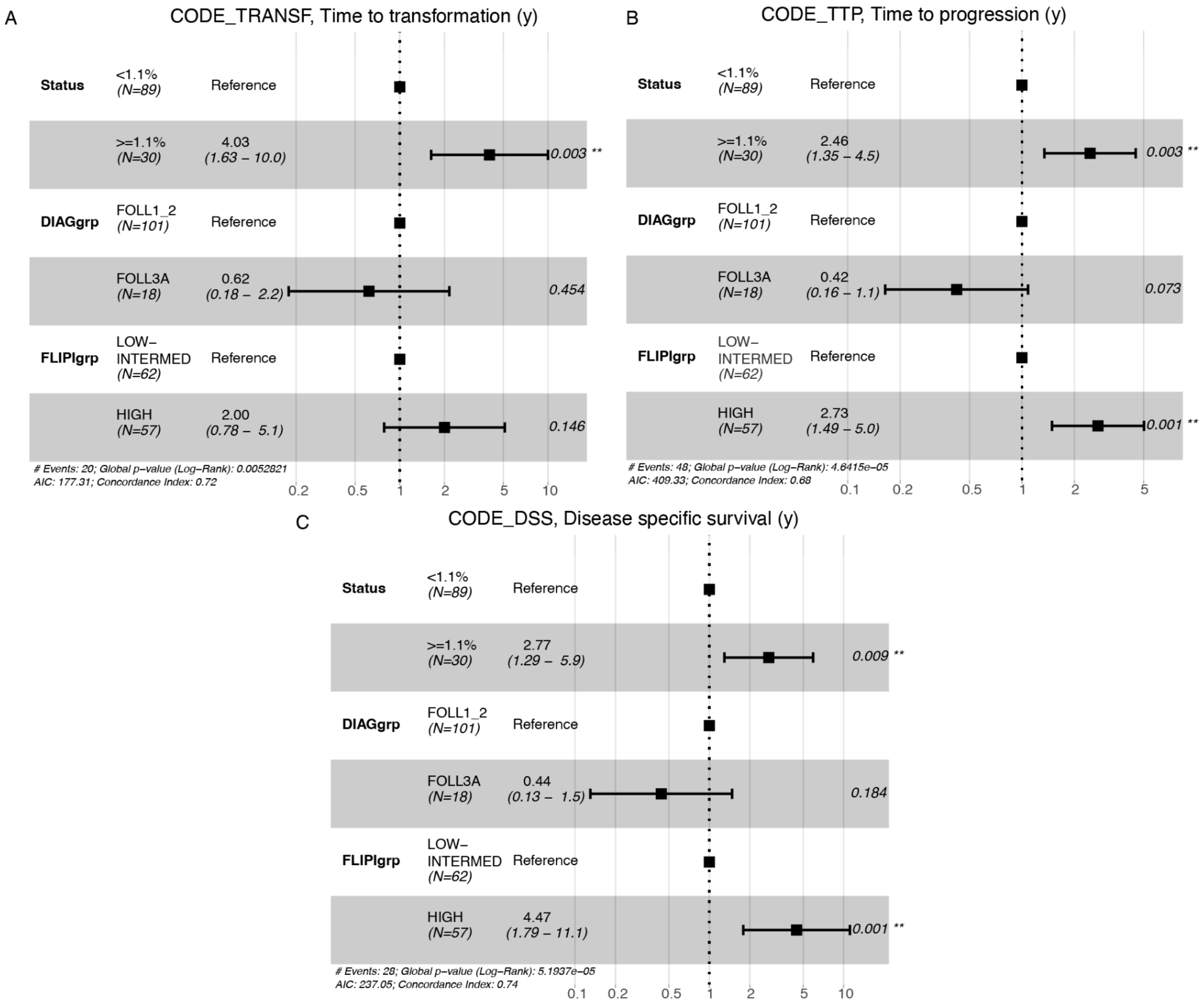


**Supplementary Figure 34:** Multivariate analysis in R-CVP cohort. The model included FLIPI score (low, versus intermediate, versus high), pathological grading (1-2 versus 3A) and CD8LAG3 infiltrate (low versus high).

**Panel A:** Cox model for time to transformation.

**Panel B:** Cox model for time to progression.

**Panel C:** Cox model for disease specific survival.

### References

Aoki, T., Chong, L.C., Takata, K., Milne, K., Hav, M., Colombo, A., Chavez, E.A., Nissen, M., Wang, X., Miyata-Takata, T., et al. (2020). Single-Cell Transcriptome Analysis Reveals Disease-Defining T-cell Subsets in the Tumor Microenvironment of Classic Hodgkin Lymphoma. Cancer Discov. *10*, 406–421. .

Campbell, K.R., Steif, A., Laks, E., Zahn, H., Lai, D., McPherson, A., Farahani, H., Kabeer, F., O'Flanagan, C., Biele, J., et al. (2019). clonealign: statistical integration of independent single-cell RNA and DNA sequencing data from human cancers. Genome Biol. *20*, 54. .

Freeman, C.L., Kridel, R., Moccia, A.A., Savage, K.J., Villa, D.R., Scott, D.W., Gerrie, A.S., Ferguson, D., Cafferty, F., Slack, G.W., et al. (2019). Early progression after bendamustine-rituximab is associated with high risk of transformation in advanced stage follicular lymphoma. Blood *134*, 761–764. .

Hie, B., Bryson, B., and Berger, B. (2019). Efficient integration of heterogeneous single-cell transcriptomes using Scanorama. Nat. Biotechnol. *37*, 685–691. .

Jin, S., Guerrero-Juarez, C.F., Zhang, L., Chang, I., Ramos, R., Kuan, C.-H., Myung, P., Plikus, M.V., and Nie, Q. (2021). Inference and analysis of cell-cell communication using CellChat. Nat. Commun. *12*, 1088. .

Kridel, R., Xerri, L., Gelas-Dore, B., Tan, K., Feugier, P., Vawda, A., Canioni, D., Farinha, P., Boussetta, S., Moccia, A.A., et al. (2015). The Prognostic Impact of CD163-Positive Macrophages in Follicular Lymphoma: A Study from the BC Cancer Agency and the Lymphoma Study Association. Clin. Cancer Res. *21*, 3428–3435. .

Salehi, S., Dorri, F., Chern, K., Kabeer, F., Rusk, N., Funnell, T., Williams, M.J., Lai, D., Andronescu, M., Campbell, K.R., et al. Cancer phylogenetic tree inference at scale from 1000s of single cell genomes. https://doi.org/10.1101/2020.05.06.058180.

Zahn, H., Steif, A., Laks, E., Eirew, P., VanInsberghe, M., Shah, S.P., Aparicio, S., and Hansen, C.L. (2017). Scalable whole-genome single-cell library preparation without preamplification.

Nat. Methods *14*, 167–173. .
